# Memory from variability: Heritable short-term cellular memory emerges from stochastic biochemical reaction networks

**DOI:** 10.64898/2025.12.31.694479

**Authors:** Mark S. Aronson, Brian Y. Zhou, James E. Fitzgerald, Allyson E. Sgro

## Abstract

Cells exhibit a mysterious form of selective heritable short-term memory, influencing outcomes as diverse as cell fate decisions in embryos and environmental responses in cancer cells and bacteria. Here, we present a simple theoretical framework explaining how this selective memory can arise from the reactions regulating molecular levels in cells. Our key insight is that related cells retain more similar molecular concentrations relative to random cells when a greater variance of possible concentration states is created during a single cell generation than is created by cell division across a population. This persistence of molecular similarity down a lineage constitutes a form of heritable short-term memory. We identify the biochemical networks that produce, modify, and degrade molecules as an underexplored source of these additional molecular concentration states. Using experimentally informed simulations, we find that the strength and duration of molecular similarity down a lineage depend on tunable network properties, explaining why some cellular traits persist only briefly while others last generations. These contributions to molecular concentration variance from biochemical reaction networks act in concert with gene expression and other regulatory processes to shape the protein composition of cells. Our framework yields clear, testable predictions for determining how biochemical network architectures drive non-genetic cellular inheritance.

## Introduction

Cells across all kingdoms of life organize into multicellular communities of genetically identical individuals, ranging from bacteria in biofilms to complex tissues in organisms (1, 2). Within these communities, cells often functionally specialize, either temporarily adopting distinct gene expression programs or traits for a few generations or permanently committing to a cell fate (3–5). Recent experimental studies have highlighted the importance of relatedness among cells in some of these temporary and permanent specializations. For example, bacteria inherit protein levels and traits that persist for 2–10 generations (6) and maintain a short-term memory of environmental conditions that influences their bias toward biofilm formation or other behaviors (7–10), cancer cells inherit gene expression patterns that can contribute to drug resistance (11), and embryonic cells inherit signaling molecule levels that control their eventual fate (12). However, this situation presents a paradox: similarity persists because amounts of some molecules remain correlated after cell division, yet fluctuations in molecule counts after cell division are thought to introduce heterogeneity rather than create similarity (6, 12–17).

An underexplored potential contributor to multigenerational correlations in levels of regulatory molecules is how cells regulate molecules during a single cell generation. The biochemical reactions that create and consume regulatory molecules are inherently stochastic and are layered into complex networks that activate gene expression programs in response to environmental inputs (18–20). Yet, the properties of these reactions are difficult to directly measure: protein and gene expression measurements are often taken at fixed time points, where information about fluctuations within a single cell generation is lost and fluctuations that last multiple generations must be inferred from population-wide measurements (11, 21, 22). Thus, as a first step toward explaining experimentally observed patterns of similarity, a theoretical framework is needed to identify how these stochastic reactions lead to similar molecular concentrations down lineages, as if restrained by their parentage (6), while other molecular levels remain uncorrelated.

Here, we develop such a framework and show how stochastic biochemical reactions can generate correlations in regulatory molecule levels that persist for multiple generations, thus acting as a form of short-term memory or a molecular form of “lineage-associated similarity” (Figure S1). This framework predicts that similarity arises when the variance of possible molecular concentrations created by stochasticity in the biochemical reactions that occur during a single cell generation exceeds the variance introduced during cell division. The magnitude and persistence of similarity are controlled by the amplification and turnover rates of these reactions, as well as the regulatory network structure. We validate these findings across multiple reaction types and network architectures, demonstrating the potential of this mechanism to function as a form of cellular memory driving trait inheritance. Importantly, this stochasticity driven by signaling networks upstream of gene expression is predicted to work in tandem with gene expression regulation and noise to create this short-term heritable non-genetic memory. Finally, to guide future experiments, our framework makes clear, testable predictions for how amplification, turnover, and pathway architecture mod-ulate similarity through generations, informing efforts to understand and potentially tune cellular memory through selectively heritable molecular similarity.

## Results

### A framework to connect intracellular signaling pathway structure to lineage-associated similarity

Lineageassociated similarity for a given molecule indicates the extent to which the molecule’s concentration in a given cell is more similar to the concentration in its sister cell than in a randomly chosen cell. This molecular similarity can drive behavioral and phenotypic similarity. Mathematically, if sister cells have similar concentrations of a molecule, the variance of concentration differences between pairs of sister cells is smaller than the variance of concentration differences between pairs of random cells in the population (Figure 1A). To quantify this in a single metric, we introduce the lineage-associated similarity (LAS) index:

**Fig. 1.**
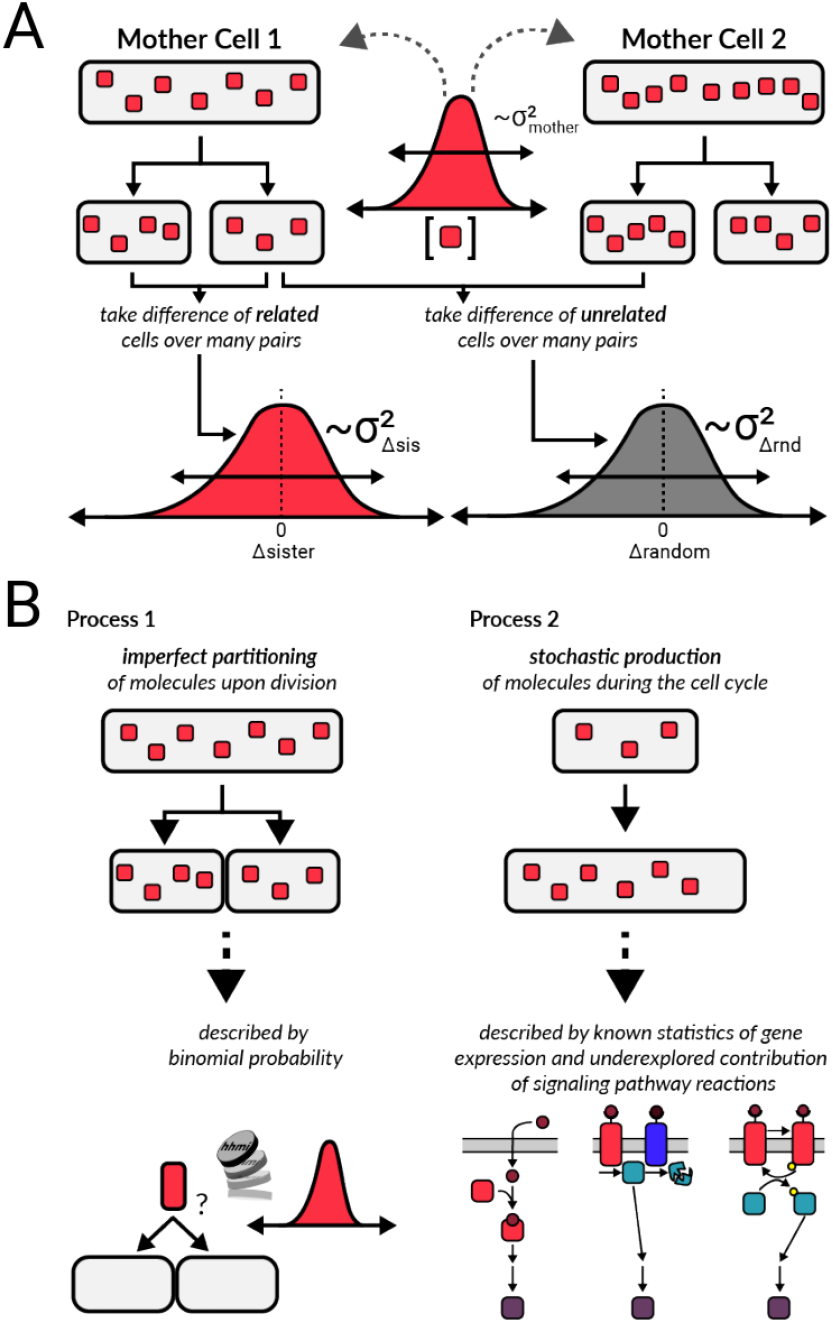
Lineage-associated similarity occurs when the molecular counts of sister cells are more similar to one another than random pairs of cells. **(A)** Schematic for comparing numbers of a molecule to create a metric for lineage association. Sister cell comparisons are made by taking the difference in the number of a specific molecule between cells arising from the same mother cell (Δ*sister*), while random cell comparisons are made between cells arising from different mother cells (Δ*random*). Comparisons over many pairs build up distributions of differ-ences characterized by a variance (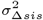 and 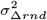). **(B)** Two key processes shape the distributions of molecules inside cells. The first process is cell division, where most molecules are divided at random. This leads to imperfect partitioning of molecules and is well described by a binomial distribution, where the probability of a molecule ending up in either daughter cell is 50%. The second key process is stochastic production and degradation of molecules during a single cell generation, controlled by signaling pathways and gene expression. In this work, we focus on the underexplored contribution of biochemical reactions by examining signaling pathway reactions that feed into gene expression and how these reactions influence molecular concentration distributions and thus lineage-associated similarity.

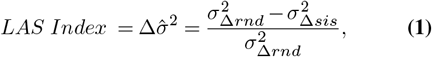

where 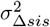 and 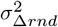 are the variance of the differences between sister cells and random cell pairs, respectively. Values near zero indicate a lack of lineage-associated similarity (as 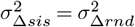), whereas larger values indicate the presence of lineage-associated similarity (as 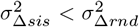). This metric is 0 when sister and random pairs of cells are equally variable and 1 when sister cells are identical to one another but not to other cells in a population.

This metric can be applied to experimental data and used in combination with analytical derivations and simulated experiments to gain insight into which features of biochemical networks, such as signaling networks, create similarity between cells (see SI Appendix A for how this metric extends to multiple generations and compares to the Pearson correlation coefficient). To this latter end, we formulated a conceptual framework to identify which processes and properties determine the amount of a specific molecule in a cell. This framework informs our analytical derivations for both the simplest biochemical reaction that we consider as well as our simulations of more complex reactions and signaling networks (see SI Appendix B for the full conceptual framework description, SI Appendix C for the simulation construction, and SI Appendix D for the analytical derivations). Briefly, molecular concentrations are controlled by the production and degradation of molecules through biochemical reactions between cellular divisions as well as partitioning of these molecules at cell division (Figure 1B). It has been well established that most molecular species are divided by random chance, as if by a coin flip, at cell division (23, 24). During the interdivisional time, molecules are created, modified, and destroyed by simple biochemical reaction motifs such as production, degradation, binding, phosphorylation, and gene expression. While the variability of gene expression has been well explored across the kingdoms of life and can range from sub-to super-Poissonian (25–28), the contributions of other processes to molecular distributions have not been well described. The likelihood of any molecule being created is either set by a fixed probability, as observed in many experiments quantifying gene expression, or determined by a more complex reaction with some rate constant (see SI Appendix C.1 for parameter ranges). By mathematically modeling each of these motifs and layering them into complex networks, we can explore how the properties of biochemical reactions affect molecular concentrations down lineages.

### Lineage-associated similarity is present only when molecules have a larger variance in possible concentrations generated during a single cell generation compared with those generated by cell division

Considering a simple toy biochemical reaction helps illuminate the origins of lineage-associated similarity. Consider a production and degradation reaction in which some enzyme A (produced at a fixed probability *P*_*prod,A*_) produces a product B at some rate *k*_*cat,A*_ during a single cell generation time (cell cycle) of length *T*_*cc*_ until cell division, at which point we simulate division of the mother cell and compare the molecular concentrations between sister cells and random cells. A second enzyme C can also be created with a fixed probability (*P*_*prod,C*_) and degrade product B at a rate set by a maximal rate constant (*k*_*cat,C*_) and half-saturation constant (*K*_*M,B*_) to consider the role of turnover in similarity (Figure 2A). We start by considering the toy model with just A and B, as this saturated production portion of the system is simple enough to explore both analytically and through simulations. Analytically, we can compute the variances of the differences between sister and random cells, 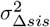 and 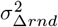, for both enzyme A and product molecule B immediately after a division event (Figure 2A, Snapshot Analyses). For enzyme A, we find the following:

**Fig. 2.**
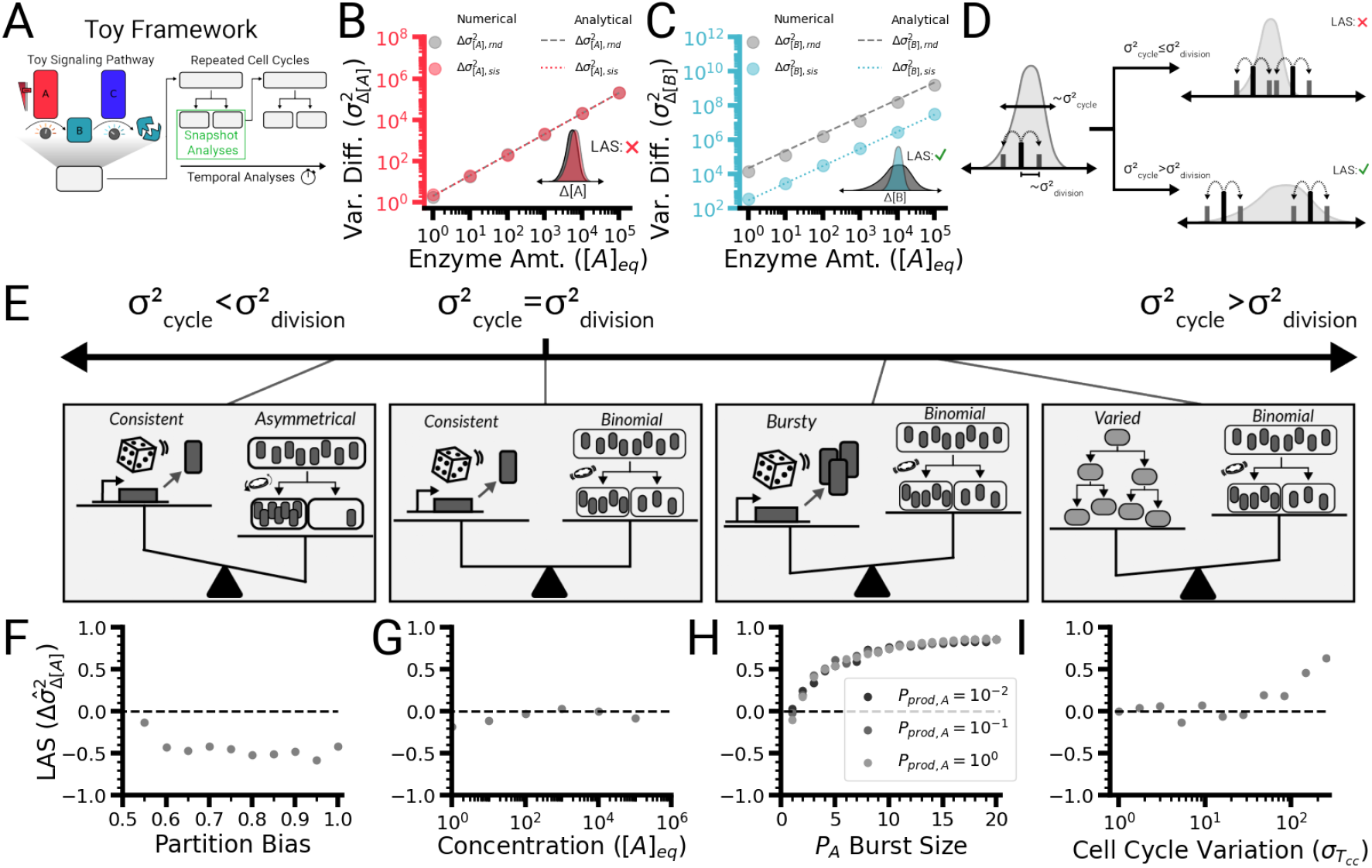
A toy model reveals the features of biochemical reactions that tune the amplitude and duration of lineage-associated similarity. **(A)** Schematic depicting a simple production and degradation motif in which enzyme A produces product molecule B and enzyme C degrades molecule B. The amounts of enzyme A can be tuned, as well as the reaction rates of each enzyme. **(B)** Variance of pairwise differences in the concentrations of enzyme A for sister and random cell pairs at various enzyme concentrations for a saturated production motif, where enzyme A is making product molecule B at a fixed rate. Dotted lines represent analytical derivations (see SI Appendix D.3), and circular markers represent numerical simulations. Note that the results for the sister and random cell pairs are identical and thus overlap each other in the plot. **(C)** Variance of pairwise differences in the concentrations of product molecule B for sister and random cell pairs at various enzyme concentrations for a saturated production motif, where enzyme A is making product molecule B at a fixed rate. Dotted lines represent analytical derivations (see SI Appendices C.9 and C.10), and circular markers represent numerical simulations. **(D)** Conceptual model for how lineage-associated similarity is created by molecular variance. Schematic representing the variance of a given molecule across a population of cells. When a mother cell (dark, vertical marker on axis) divides, binomial partitioning “pushes” each daughter cell (shorter, lighter vertical markers) in equal and opposite directions on the concentration axis. The width of the overall distribution is related to the variance in molecular concentration from processes during a single cell generation 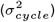, while the distance between the daughter cells and the mother cell is related to the variance in molecular concentration from division 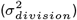. If the variance from a single cell generation is less than that from division, the narrow distribution leads to daughter cells from different mothers being close to each other in the concentration space (top). If the variance from a single cell generation is larger than that from division, daughter cells from different mothers are more likely to be far away from each other in the concentration space (bottom). **(E)** Spectrum of the relationship between variance generated during a single cell generation and variance generated during division. When the variance generated during a single cell generation is low, as in the case of consistent gene expression and asymmetric partitioning, lineage-associated dissimilarity occurs, where random pairs of cells are more likely to have similar molecular concentrations than related pairs **(F)**. When the variance generated during a single cell generation is equal to the variance generated upon division, as with consistent gene expression and binomial partitioning, sister pairs are as similar to each other as any random pair in the population **(G)**. When the variance generated during a single cell generation is greater than the variance generated upon division, lineage-associated similarity is present. This occurs in the case of bursty gene expression and binomial partitioning **(H)** as well as varied single cell generation times and binomial partitioning **(I)**. Each variance value is calculated from 1000 pairs of sister cells and 1000 pairs of random cells. See Appendix G for simulation parameters.

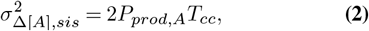

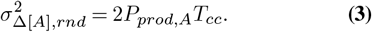

As these two difference distributions have identical variances for enzyme A (Figure 2B; for full distributions, see Figure S2), our similarity metric, 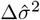, is 0 (see SI Appendix D.3 for the derivation). This result indicates that lineage-associated similarity does not exist for enzyme A at any concentration. Our analytical solutions provide an explanation for this phenomenon: they highlight that the variance in the molecule A count generated during a single cell generation (from the Poisson production) and the variance generated upon division (from the binomial partitioning) perfectly counteract each other (SI Appendix D.3), as has been noted elsewhere (15). Similarly, our simulations show that while the coefficient of variation in A’s concentration decreases with increasing concentration of A, the relative partitioning error also decreases (Figure S3).

However, the difference distributions for the product molecule B in sister cells and random pairs are not identical (Figure 2C; for full distributions, see Figure S2). Analytically, we have the following:

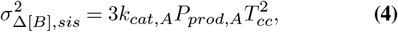

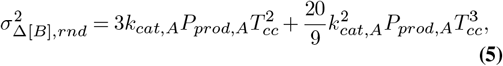

so that the lineage-associated similarity in product molecule B is

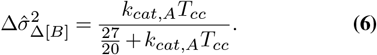

This expression indicates that not only does lineage-associated similarity exist in product molecule B, its level is independent of the amount of enzyme A (see SI Appendices D.9 and D.10 for a derivation). This result arises because the production of B is a doubly-stochastic process, which leads to a wider-than-Poisson variance that is not counteracted by the variance generated upon division. This doubly-stochastic process is conceptually similar to the well-studied Cox processes (29), where the doubly-stochastic variable always has a larger variance than the singly-stochastic variable.

More generally, this finding suggests a simple explanation for the origin of lineage-associated similarity (Figure 2D and E). Specifically, the amount of any molecule in a cell is shaped by the amount inherited during cell division, as well as how it is created and consumed. Thus, the similarity in molecular levels between cells is a function of the variance stemming from each of these processes. When the variance generated by biochemical reactions and other processes that create and consume molecules is larger than the variance generated upon division, related cells are more similar to each other than random cells in the population, resulting in lineage-associated similarity. When these variances are equal, there is no lineage-associated similarity. The variance introduced by division can also exceed the variance from the biochemical reactions creating and destroying molecules, making related cells more dissimilar to each other than random cells in the population. For example, mechanisms that asymmetrically partition molecules upon division (16, 30–32) can increase variance from division and lead to lineage-associated dissimilarity (Figure 2F). When the production and division processes have equal variances such as observed in enzyme A in our toy model, there is no lineage-associated similarity (Figure 2G). Other sources of variance such as bursty gene expression and varying single cell generation times increase variance in molecule counts during a single cell generation and create lineage-associated similarity (Figures 2H and I). As the variance generated by gene expression noise has been well described elsewhere (33) and the division of most molecules is well established as binomial (23, 24), here we focus on the unexplored contributions of biochemical reactions and network structure to molecular distribution variance and thus lineage-associated similarity.

### Enzymatic reaction amplification is a key modulator of the magnitude of lineage-associated similarity

Given that the amount of enzyme A does not appear to change the magnitude of lineage-associated similarity in product molecule B (Figure 2C), what parameters could tune its magnitude? Our analytical derivations indicate that in addition to the enzyme amount, the reaction rate of enzyme A (*k*_*cat,A*_) and the generation time between cell divisions (cell cycle time, *T*_*cc*_) are also critical parameters (SI Appendices D.9 and D.10). Note that the concentration of the enzyme (*P*_*prod,A*_*T*_*cc*_) tunes both the random and sister cell difference distributions by the same amount; thus, it does not affect similarity. In contrast, the reaction rate of A (*k*_*cat,A*_) and the cell cycle time (*T*_*cc*_) both affect similarity. The combined value (*k*_*cat,A*_*T*_*cc*_) is the average number of product B molecules created by one enzyme A at saturation during each cell cycle. Thus, this value can be considered the amount of amplification in the motif or the “amplification factor.” Focusing again on only a saturated production reaction (Figure 2A), the lineage-associated similarity of product molecule B is a function of the amplification factor (Figure 3A, solid black line) with a maximum possible value approaching 1 (i.e., as the amplification factor increases, sister cells approach being identical). Supporting our analytical derivation, our numerical simulations, sweeping through either component parameter (enzyme maximal reaction rate, *k*_*cat,A*_, or cell cycle time, *T*_*cc*_), show similar results (Figure 3A, scatter plot; see Figure S4 for sweeps covering the full range of amplification factors and Figure S5 for a two-dimensional sweep of both parameters). This finding suggests that amplification is an important modulator of lineage-associated similarity, with a higher amplification creating wider possible molecular distributions and thus more space for cells to remain similar to their siblings (Figure 3B).

**Fig. 3.**
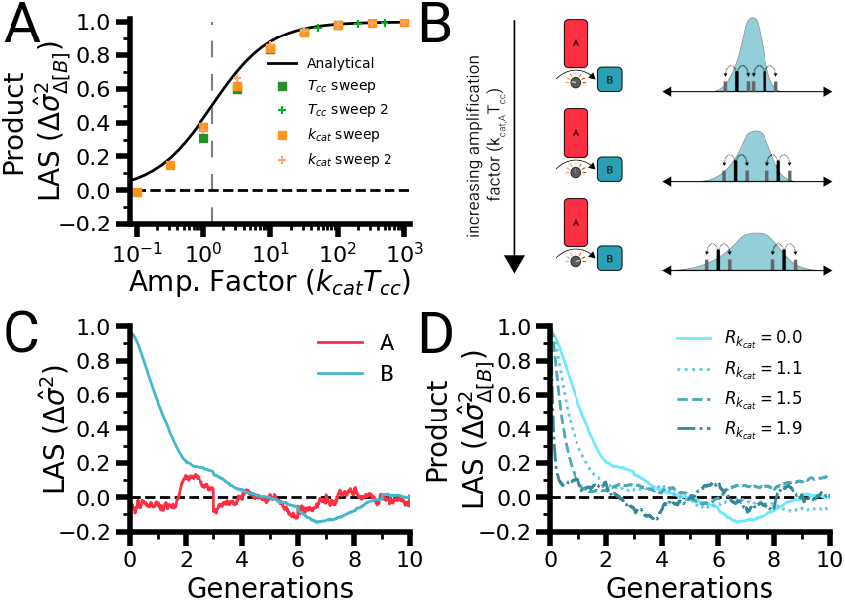
Factors that change the magnitude and duration of lineage-associated similarity. **(A)** Lineage-associated similarity for product molecule B at various values for the amplification factor, which is the combined value of the reaction rate of enzyme A and the cell cycle time (*k*_*cat,A*_*T*_*cc*_). The analytical solution (Equation 6) is shown (solid black line, inflection value of 1.35 indicated by a vertical gray dashed line) alongside computed values for simulations sweeping both contributors to the amplification factor: cell cycle time (*T*_*cc*_, green squares and crosses) and reaction rate (*k*_*cat,A*_, orange squares and crosses). The fitted inflection point for the *T*_*cc*_ sweep is 2.14 and for the *k*_*cat,A*_ sweep is 1.85. See Figure S4 for extended parameter sweep results. **(B)** As the amplification factor increases, the distribution of possible molecular concentrations widens, creating more space for sister cells to retain similar molecular counts after division. **(C)** Lineage-associated similarity (as measured by the normalized difference of variances) over ten cell generations for both components of the saturated production motif. **(D)** Lineage-associated similarity of the product molecule B (as measured by the normalized difference of vari-ances) over ten cell generations for different values of the 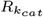 ratio, which alters the turnover rate of the product molecule in the motif. See SI Appendix G for all simulation parameter values.

In addition to reaction rate and cell cycle time, the amount of reactant can also tune the amount of amplification in a reaction by not fully saturating the reaction’s production potential. By fixing the amount of reactant, we can demonstrate that the amount of reactant tunes the amount of lineage-associated similarity along the same curve (Figure S6). Similarly, we explored binding motifs in which two molecules associate together to produce a bound product. In our analyses of both reversible and irreversible binding motifs (SI Appendices C.5 and C.6), we found no similarity in any component (Figure S7). Together, these findings reinforce the notion that creating a wide variance of possible molecular states is critical for lineage-associated similarity: amplification creates a wider distribution.

### Molecular turnover rate determines the duration of lineage-associated similarity

Thus far, we have explored which properties of biochemical reactions create and modulate lineage-associated similarity in the generation immediately after cell division. Our framework explains experimental observations of similarity that persist past a single generation, such as how different levels of phosphorylated ERK that build up over a single cell generation persist after division and drive cell fate decisions (12). However, some traits appear to persist over multiple generations without a clear cell fate commitment mechanism (6). Thus, we extended our framework beyond a single cell division (see SI Appendices A and B) to develop an intuition for which parameters could generate and tune multigenerational lineage-associated similarity. As expected from the single-generation results, enzyme A exhibited no similarity (Figure 3C, red, and Figures S8B–D^*′*^) while the similarity metric of product molecule B decayed from its initial value to near zero over 2–3 cell generations (Figure 3C, cyan). Our simulation result matches a previous experimental observation in bacteria (6), in which the fluorescence of green fluorescent protein (GFP) molecules expressed from a constitutive promoter between related *Escherichia coli* cells exhibited short-term correlation (6). If we treat the messenger RNA (mRNA) of the GFP as molecule A, as it is not consumed when it produces product molecule B, and the protein as the product molecule B in our saturated production motif, our framework perfectly predicts the two generations of correlation measured in these cells.

Which, if any, of the parameters affect the duration of similarity? We find that the amount of enzyme and reaction rate do not affect the number of generations of similarity (Figures S8E and F). However, an as-yet unexplored parameter is the molecular turnover rate, which is controlled by the cell division frequency and any active degradation of the molecule (34). To explore the impact of the molecular turnover rate, we added active degradation to our saturated production motif (see SI Appendices B and C.9). Specifically, we added a degrading enzyme C that increases the turnover rate of the product molecule B. We can simultaneously alter the maximal reaction rates for both the producing enzyme A and the degrading enzyme C, which we describe with a ratio 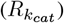:

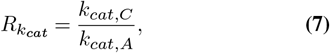

where *k*_*cat,C*_ is the maximal reaction rate of the degrading enzyme C and *k*_*cat,A*_ is the maximal reaction rate of the producing enzyme A. This ratio ranges from 0, indicating no degradation activity, to 2, the maximal amount of degradation activity that maintains a nonzero amount of product molecule B. As the degradation rate increases, the correlation time of the product molecule B decreases (Figure 3D; see Figure S9 for a full sweep). As both increasing the cell division frequency (Figures S8G and G^*′*^) and increasing degradation lead to a reduced correlation time, our findings suggest that the molecular turnover rate sets the duration of lineage-associated similarity for a given molecule.

### Lineage-associated similarity in multiple biologically realistic pathways and parameter regimes

Our previous cases focused on an idealized toy model that provides insight into the properties that tune the magnitude and duration of lineage-associated similarity. In reality, biochemical reactions do not have unlimited amounts of reactant substrate, but operate at or near saturation (Figure 4A) (35) with reactant amounts in the range of 100 *µ*M–10 mM and reactant:enzyme ratios of 1:1 – 100:1 (36). To explore these near- or at-saturation behaviors in all the molecules involved in a reaction, we simulated a more realistic production motif in which the substrate is produced, like the enzyme, by a Poisson process (Figure 4A, SI Appendix C.4) and analyzed the lineage-associated similarity for each component across a range of substrate amounts. As with the saturated production motif, we observed no similarity in the amount of enzyme A (Figure 4B, red). However, we observed similarity in the reactant near the point of saturation that then decreased sharply after saturation (Figure 4B, purple). We also observed a monotonic, switch-like increase in similarity of the product with respect to the amount of reactant starting near the point of saturation and leveling off in the fully saturated regime (Figure 4B, blue), suggesting that lineage-associated similarity in the product exhibits zero-order ultrasensitivity to the amount of reactant (see SI Appendix E for a further exploration of the relationship between ultrasensitivity and lineage-associated similarity). We observed a similar dependency on reactant saturation for lineage-associated similarity in a simple phosphorylation motif (Figure S10) and in a more biologically realistic phosphorylation motif with dual kinase-phosphatase activity (Figure S11).

**Fig. 4.**
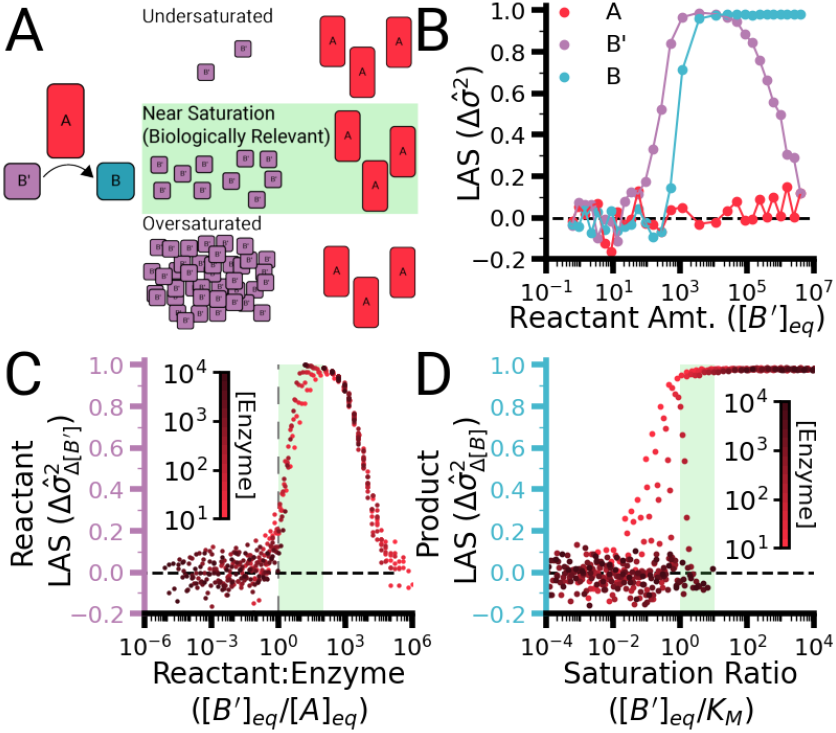
Lineage-associated similarity in the reactant and product molecules exists in biologically realistic regimes. **(A)** Schematic depicting the production motif, in which a reactant molecule (*B*^*′*^) is converted into a product molecule (*B*) by an enzyme (*A*) and the different regimes of reactant and enzyme ratios. **(B)** Lineage-associated similarity of each component of the production motif for varying reactant concentration values (generated by sweeping the reactant production probability, 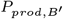). **(C)** Lineage-associated similarity of the reactant molecule (*B*^*′*^) as a function of the mean reactant-to-enzyme concentration ratio (ratio of 1:1 is indicated by a vertical dashed line) over a sweep of enzyme A concentrations spanning three orders of magnitude (colorbar). Shading indicates a physiologic stoichiometric ratio range of 1:1–100:1 (36). **(D)** Lineage-associated similarity of the product molecule (*B*) as a function of the saturation ratio over a sweep of enzyme A concentrations spanning three orders of magnitude (red slider). Shading indicates a physiologic range where the ratio of reactant concentration to its *K*_*M*_ value is 1:1–10:1. See Appendix G for all simulation parameter values

To better understand if lineage-associated similarity exists in these biologically realistic regimes, it is helpful to consider the natural relationship between reactants and enzymes. When considering how reactant concentrations are shaped, the reactant:enzyme ratio is the most important factor because the enzyme is the component that modifies the reactant concentration. These ratios are in the range of 1:1 – 100:1 (36), and we see that lineage-associated similarity begins to spike in this regime (Figure 4C). Similarly, the ratio between the amount of reactant and the half-saturation constant of the enzyme dictates the amount of product produced because the reactant is turned into the product at some rate related to the half-saturation constant of the enzyme. In the 1:1–10:1 regime found *in vivo* (36), lineage-associated similarity reaches a maximum (Figure 4D). Multiple mechanisms create wide variances in possible molecular concentrations near and at saturation in the reactant and product, which we explore in SI Appendix F. Our findings suggest that the capacity for a reaction to be saturated is important for creating wide variance distributions and thus creating lineage-associated similarity.

### Linear composition of biochemical reactions in signal pathways leads to an additive correlation time of lineage-associated similarity

Signaling pathways driving traits, behaviors, and cell fates within cells are composed of multiple biochemical reactions similar to the ones we have analyzed. To confirm that lineage-associated similarity can affect traits over multiple generations in real biological signaling networks, we turned to three critical classes of bacterial intracellular signaling pathways that mediate intracellular gene expression and thus behavioral changes in response to extracellular changes (20): a diffusible ligand circuit, a second messenger system, and a two-component system.

Our diffusible ligand circuit is composed of an irreversible binding motif, where a ligand that diffuses into the cell binds to a transcription factor, followed by a saturated production motif where the transcription factor drives expression of a protein gene product (Figure 5A, SI Appendix C.10). While the ligand and both unbound and bound transcription factors have no lineage-associated similarity, the gene product has 2– 3 generations of lineage-associated similarity, as previously observed in the saturated production motif where one component amplifies the production of a second component (Figure 5B).

**Fig. 5.**
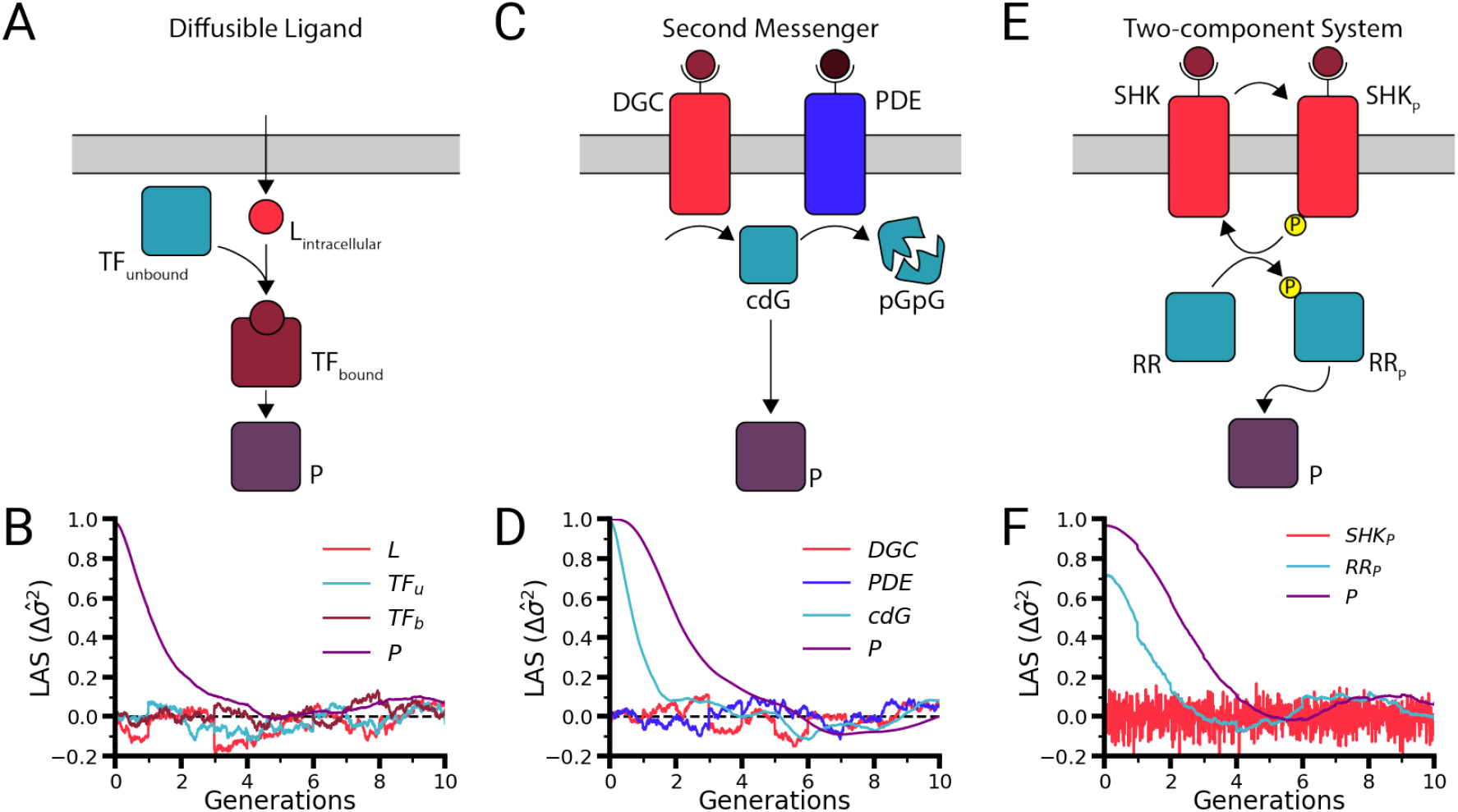
The lineage-associated similarity correlation time of a signal pathway output is dictated by the addition of individual reaction similarity correlation times. **(A)** Schematic depicting a diffusible ligand–transcription factor pathway, which is composed of a binding motif followed by a saturated production motif. Ligand molecule L diffuses into a cell, where it irreversibly binds with unbound transcription factor (*TF*_*unbound*_) to form a bound complex (*TF*_*bound*_). This bound complex drives the gene expression of protein product *P*. **(B)** Lineage-associated similarity (as measured by the normalized difference of variances) over ten cell generations for the components of the diffusible ligand pathway. **(C)** Schematic depicting a second messenger pathway (shown here as a cyclic di-GMP pathway), which is composed of a production and degradation motif followed by a saturated production motif. Here, the diguanylate cyclase enzyme (DGC) produces a cyclic di-GMP product molecule (cdG), which is degraded into pGpG by the phosphodiesterase (PDE). cdG drives the gene expression of protein product P. **(D)** Lineage-associated similarity over ten cell generations for the components of the second messenger pathway (here, 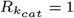; see Figure S9 for all values). **(E)** Schematic depicting a two-component system, composed of a phosphorylation motif followed by a saturated production motif. A sensor histidine kinase (SHK) autophosphorylates (producing *SHK*_*P*_), which then transphosphorylates an unphosphorylated response regulator (RR), producing a phosphorylated response regulator (*RR*_*P*_) that drives the gene expression of a protein product (*P*). **(F)** Lineage-associated similarity over ten cell generations for the components of the two-component system pathway. See SI Appendix G for all simulation parameter values.

To understand how a second messenger pathway would behave, we simulated a simple cyclic di-GMP circuit (37). Here, the signal molecule, cyclic di-GMP (cdG), is produced by a diguanylate cyclase (DGC) and degraded by a phosphodiesterase (PDE). cdG activates the gene expression of a protein product P (Figure 5C, SI Appendix C.11). As expected for a production and degradation reaction, no lineage-associated similarity was measured for DGC or PDE, yet a moderate correlation time of almost two generations was observed for cdG (Figure 5D, cyan). The saturated production motif of the protein product then extended the similarity time to over three generations for the protein product P (Figure 5D, purple). This result suggests that correlation times are additive when reactions are arranged in linear pathways.

Lastly, we analyzed the monofunctional sensor histidine kinase two-component system circuit, which consists of a phosphorylation motif followed by a saturated production motif (Figure 5E, SI Appendix C.12). As expected for the phosphorylation motif operating at sensor histidine kinase saturation (Figure S10), lineage-associated similarity was absent in the phosphorylated sensor histidine kinase (Figure 5F, red) and present for two generations in the phosphorylated response regulator (Figure 5F, cyan). The saturated production motif generating protein product P resulted in two additional generations of correlation, leading to four generations of lineage-associated similarity in the protein product P (Figure 5F, purple), reconfirming that a linear composition of the signal pathway motif leads to additive durations of lineage-associated similarity. We further validated this additive property by simulating a five-layer cascade, finding that each layer added approximately two generations of lineage-associated similarity (Figure S12).

### Molecules exhibiting lineage-associated similarity create regional concentration similarity in spatially structured communities

As multicellular communities grow, they also organize and develop a spatial structure; thus, we sought to determine how lineage-associated similarity persists not only over generations but also over space. To demonstrate how multigenerational lineage-associated similarity in a signaling pathway impacts the spatial structure of traits in a multicellular community, we extended our lineage-associated similarity model to consider space. Briefly, we simulated the growth and division of cells spatially, as has been previously done for bacterial biofilms ((39); see Methods and Figure S13 for details), to produce a growing collective of non-motile cells, where cells remain close to their kin (Figure 6A; Figures S14 and S15). To understand how relatedness impacts the spatiotemporal distribution of traits over time, we considered a community of cells where each cell contains a saturated signal relay circuit (Figure 6A), similar to the second messenger circuit without degradation (Figure 5C), where an enzyme produces a molecule that then drives production of the protein product. This condition creates a community in which each molecule, and thus the trait that it creates, has a different pattern in space that results from the different lengths of lineage-associated similarity for that molecule (Figure 6B). As expected from our previous results (Figure 5D), each additional layer in a simple saturated production circuit results in an additional two generations of similarity, for a total of approximately four generations of similarity in the final product molecule (Figure 6C). Over ten generations, the increasing multigenerational correlation times in molecules farther down the pathway create larger spatial neighborhoods of correlated cells (Figure 6D). Importantly, these results demonstrate that stochastic biochemical reactions occurring at the molecular level impact the spatiotemporal distribution of traits in a multicellular collective, providing insight into how molecular similarity can spread throughout a multicellular system.

**Fig. 6.**
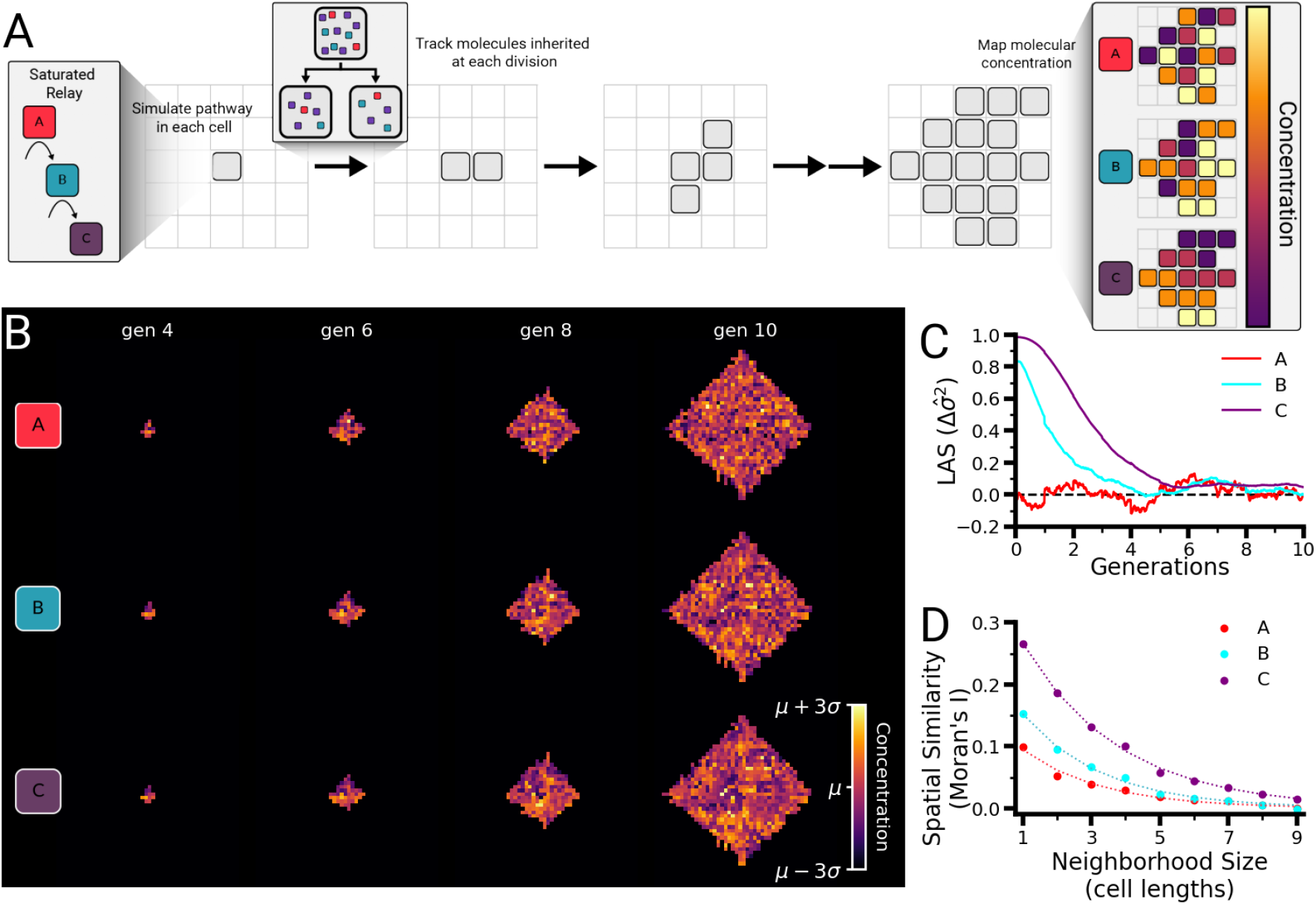
Spatial similarity of molecular concentration in a multicellular community arises from lineage-associated similarity. **(A)** Schematic of spatial lineage-associated similarity simulations. A saturated relay pathway is simulated in each cell of the simulation, starting from a single cell that replicates to produce a microcolony of cells. At each division, each molecular species is binomially partitioned into the resulting daughter cells. At each simulation time point, the concentrations of each molecule can be visualized across the entire lineage. **(B)** Temporal snapshots of concentration maps of the components of the saturated relay pathway across a simulated multicellular community. **(C)** Lineage-associated similarity over ten cell generations for the components of the saturated relay pathway. **(D)** Moran’s I quantification (38) of local spatial similarity for each component of the saturated relay pathway (see Figure S16 for neighborhood definition). Values were computed from the final time frame of the simulation depicted in **(D)**. See SI Appendix G for all simulation parameter values.

## Discussion

The analyses and simulations presented above have demonstrated how stochasticity in biochemical reactions can confer molecular similarity and thus trait similarity down a lineage. This lineage-associated similarity mechanism may explain a wide range of experimental observations, ranging from inherited molecular levels to the emergence of spatiotemporal patterning in multicellular systems. This similarity arises when the variance generated in a molecule’s concentration during a cell generation exceeds the variance generated upon division (Figures 2D–I). Different molecules in a cell will have different magnitudes and durations of lineage-associated similarity based on the amount of variance generated by the biochemical reactions that produce and degrade the molecule, and this similarity is additive down pathways.

### Stochasticity as a mechanism for short-term heritable gene expression patterns

In this work, we have focused on the unbranched structures of common bacterial signaling pathways that are conducive to building simple models for analyzing inheritance patterns (20). Importantly, mammalian signaling pathways have been shown to exhibit input–output linearity, making these linear bacterial pathways good approximations of some more complex mammalian regulatory networks and allowing our results to be generalized across kingdoms of life (40). Additionally, more complex regulatory pathways likely employ other feedback or degradation mechanisms similar to those explored in Figure 3D to increase the turnover rate of molecules. A high turnover rate reduces the duration of inherited molecular states and has been previously suggested as a strategy that cells could employ to limit deviations from the population average due to unequal partitioning, but at a high metabolic cost (15). Our results demonstrate that this biological strategy would keep each individual cell’s molecular concentration close to the population average, thus increasing overall population similarity while decreasing lineage-associated similarity.

For the simplest signaling network, where a product is produced at saturation, we have illustrated how stochasticity drives two generations of lineage-associated similarity (see Figure 3C). This circuit is analogous to a gene such as GFP being expressed by a constitutive promoter (where the pro-duction of the mRNA transcript is akin to the production of molecule A and the production of a protein is akin to the production of molecule B), and our estimate is in line with experimental observations of the duration of GFP expression similarity in bacteria (6). Thus, our framework sheds light onto the factors governing this similarity in a biological system; in turn, experiments will provide opportunities to further test and validate our framework as tools for quantifying protein counts in known regulatory networks become more readily available. Other observations that could be explained by our framework include the various levels of correlation observed for different genes down a lineage in mouse embryonic stem cells (41) or the emergence of spatial similarity in gene expression in various cancer cell lines (11).

Our proposed mechanism, where a noisier molecular concentration leads to a greater capacity for lineage-associated similarity, is an instantiation of the concept of biological noise (42). This mechanism represents another example of stochasticity amplifying a signal in a biological process, which has been demonstrated in the phenomena of stochastic focusing (43) and stochastic resonance (44). We also found that when we chain multiple reactions together into signaling pathways, downstream components have longer similarity times (Figure S12). This occurs because, without explicit noise-dampening mechanisms, downstream components have a higher variance and thus higher similarity. Previous experimental studies have found that multi-step signaling pathways increase noise in downstream products, a phenomenon termed “noise propagation” (45–47). Our results demonstrate that heritable gene expression fluctuations can be driven by this noise propagation in regulatory networks, leading to lineage-associated similarity.

### Lineage-associated similarity as a reinforcement mechanism in multicellular communities

Our spatial simulations show how stochasticity in biochemical reactions can create regional pockets of cells with similar molecular contents in non-migratory cell populations (Figure 6), which have also been described in (21) and experimentally observed in (11). These molecular-level similarities among nearby cells could drive similar responses to changes in extracellular environments, a phenomenon that has been proposed as a potential explanation of spatial responses to insulin growth factor 1 among spatially proximal fetal pulmonary arterial endothelial cells (48).

Furthermore, cells modify their own environments by emitting extracellular signaling molecules that act over only a few cell lengths, controlling cell-level properties as diverse as cell growth and horizontal gene transfer, which have important implications for the entire community (49–51). As a result, when cells are not very motile, signaling occurs between closely related cells with similar molecular states. If the signal activates instead of suppresses a pathway, it could further reinforce any spatially observed behavioral similarity. The range and strength of any behavioral correlations within uniform environments would be set by both the level of relatedness among cells in the spatial neighborhood (i.e., how motile they are) and the level of similarity among regulatory molecules that control the behavior in question.

### Verifying and exploiting stochasticity-induced lineage-associated similarity

Once the signaling network architecture that produces regulatory molecules governing a behavior or specific gene products is known, we can computationally simulate this structure to predict the duration of lineage-associated similarity for each molecule and resulting behavior. These predictions, based on verified signaling network architectures, provide a way to distinguish between our proposed mechanism and alternative explanations such as shared transcription factors or transient epigenetic modifications. While the exact signaling network architecture is often unknown and it is not yet possible to experimentally measure all of the factors contributing to transcription factor activity, estimates should still generate distinct predictions for the duration and pattern of gene products exhibiting similarity to distinguish between potential mechanisms. Importantly, we suggest that observing deviations from the duration of similarity predicted by fully accounting for sources of stochasticity in signaling networks and downstream gene regulation suggests active investment in either dissimilarity or similarity.

Emerging experimental techniques will permit us to explore how this mechanism for lineage-associated similarity may control specific trait assignments or developmental lineage decisions. For example, in mammalian embryonic development, early gene transcripts such as *Carm1* and *Cdx2*, which drive cell lineage differentiation toward the inner cell mass or trophectoderm, respectively, are binomially partitioned during the first zygotic cleavage event, consistent with our model here. Any partitioning errors would then bias differentiation toward these cell types (52). As new experimental techniques begin to permit both robust lineage tracking and the measurement of molecular counts or enzyme activity down generations, it will become possible to directly measure molecular distributions before and after division and identify when this mechanism influences key processes such as developmental fate.

In addition to explaining experimental findings in natural systems, the role of signaling network stochasticity in creating multigenerational similarity can also be both validated by and applied in synthetic systems: we can construct signaling pathways with known structures, such as synthetic networks of protein circuits (53) or phosphorylation cascades (54), and visualize reporters down entire cell lineages using microfluidic devices (6, 55). Ultimately, synthetic networks with tunable similarity durations for different molecules could enable the design of multicellular systems with heritable spatially defined gene expression patterns.

In summary, we suggest that stochasticity-induced lineage-associated similarity is a key contributor to many puzzling experimentally observed spatiotemporal gene expression patterns and fate commitments. We encourage the development of new quantitative techniques to directly connect the effects of stochasticity at the level of biochemical reactions to observations of lineage-associated similarity in multicellular systems.

## Methods

### Simulation of a cell generation

The basic algorithm for our simulation is as follows: a cell starts with a volume of 1 and a starting amount of each molecule involved in the given motif or pathway being simulated (see SI Appendix B for the conceptual framework). Initial molecule amounts are typically given as their production probability parameter multiplied by the cell cycle duration or a rough estimate of their steady-state value. A cell generation proceeds via the Gillespie algorithm (56) to select which biochemical reaction will occur in the next time step (see SI Appendix C for the details of all reactions involved in each motif or pathway). Following the Gillespie algorithm, after the next time step’s reaction is selected, a time step is calculated (SI Appendix C.16). This time step is used to calculate the volumetric growth of the cell during that time step, which is a linear growth in volume of 1 to 2, based on the 1-to 2-fL volume of *E. coli* (57). Stochastic biochemical reactions are chosen until the cell volume reaches 2, which triggers a cell division event. Upon cell division, each molecular species is partitioned into the new daughter cells using binomial random numbers. The cell volume is set to 1, and the cycle then repeats.

### Seed cell simulation

To generate pools of possible mother cells for comparisons, a starting cell is initiated, and the simulation is run for an initial 20 cell generations so that each molecular species involved in the selected motif or pathway reaches a steady-state concentration. Then, the simulation is run for a given number of cell generations (typically 1000), generating 1000 possible mother cell states (Figure S17). To make comparisons of related cells, each mother cell state is binomially partitioned using a random generator, and the difference in molecular amounts between the resulting daughter cells is computed. To make comparisons between unrelated pairs, each mother cell state is binomially partitioned, a random second mother cell from the pool (selected by a random generator) is partitioned, and the difference is calculated between the cell from the first mother and the cell from the second mother. Note that for all cells in a given simulation, the production probabilities are held constant, simulating a uniform environment for the various pathways.

### Temporal comparisons

For the temporal calculations of lineage-associated similarity (Figures 3C and D, Figure 5, and Figure 6C), the daughter cells resulting from the seed cell divisions are used to initiate new simulations that run for 10 cell generations. To make comparisons between values at stochastically decided time-steps, the concentration traces are first down-sampled to 1/100th of a cell generation. The time-steps closest to these linearly spaced times are chosen for comparisons. As with the initial values, differences are taken between the down-sampled traces of related and unrelated cells to calculate the normalized difference of variance over time.

### Spatial simulations

For the spatial simulations (Figure 6), a single cell starts at the midpoint of a Cartesian grid. Cell generation simulations are run as described above. Upon cell generation completion, the molecular contents are divided, and a new cell is formed in an adjacent location on the Cartesian grid (Figure S13). Cells divide to fill empty spaces in the cardinal directions. Once a cell is surrounded by neighboring cells, the direction with the fewest cells is chosen, and all of the cells in that row or column are pushed outward. In the case of a tie between directions, a random number generator is used to choose a direction. Cell cycle times are decided from a normal distribution around a mean such that cell division does not occur simultaneously in every cell present. Cousin calculations are performed by finding the last common ancestor between two cells and counting to the present cells (e.g., if the last common ancestor was two generations prior, then the two cells are first cousins). When cells are compared between two different generations, the lower number is chosen (i.e., cells that are first cousins once removed are treated as first cousins). For Moran’s I calculations, the neighborhood is defined as all cells within cardinal steps (Figure S16, Disc).

### Code writing

All code was written in Python 3.9.12 using Spyder 5.3.3 for an IDE and compiled using IPython 7.31.1. The packages used were NumPy 1.22.4 (58), SciPy 1.8.1 (59), Matplotlib 3.5.2 (60), and OpenCV 4.7.0.72 (61).

### Random number generation

Random number generation was performed using NumPy’s random generator function with seed 1000, unless otherwise noted.

### Simulation hardware

Code for shorter simulations and analyses were run locally on a Dell Latitude 5420 running Windows 10 Enterprise 21H1 and using an Intel 11th Gen Core i7 processor (3.0 GHz). Code for longer simulations was run on the Janelia Compute Cluster, which runs on Oracle Linux 8.3 and uses IBM Spectrum LSF 10.1 for job management. Cores were composed of either Intel SkyLake (2.7GHz Platinum 8168) or Intel Cascade Lake (3.0 GHz Gold 6284R) processors.

## Code availability

Python code for all simulations and results presented in this manuscript is detailed in SI Appendix H and available on GitHub at https://github.com/sgrolab/cellsimilaritymodel

## ACKNOWLEDGEMENTS

The authors thank members of the Sgro Lab, Dr. Oleg Igoshin, Dr. Hernan Garcia, and Dr. Archishman Raju for their helpful guidance and critical review of our work. This work was supported by the Howard Hughes Medical Institute. JEF also acknowledges support from the National Institute for Theory and Mathematics in Biology through the National Science Foundation (grant number DMS-2235451) and the Simons Foundation (grant number MPTMPS-00005320).

## Bibliography

1. Carey D. Nadell, Knut Drescher, and Kevin R. Foster. Spatial structure, cooperation and competition in biofilms. Nature Reviews Microbiology, 14(9):589–600, 2016. doi: 10.1038/nrmicro.2016.84.

2. Karin Sauer, Paul Stoodley, Darla M. Goeres, Luanne Hall-Stoodley, Mette Burmølle, Philip S. Stewart, and Thomas Bjarnsholt. The biofilm life cycle: expanding the conceptual model of biofilm formation. Nature Reviews Microbiology, pages 1–13, 2022. doi: 10.1038/s41579-022-00767-0.

3. Anissa Guillemin and Michael P. H. Stumpf. Noise and the molecular processes underlying cell fate decision-making. Physical Biology, 18(1):011002, 2020. doi: 10.1088/1478-3975/abc9d1.

4. Mark S. Aronson, Chiara Ricci-Tam, Xinwen Zhu, and Allyson E. Sgro. Exploiting noise to engineer adaptability in synthetic multicellular systems. Current Opinion in Biomedical Engineering, 16:52–60, 2020. doi: 10.1016/j.cobme.2020.100251.

5. Thomas M. Norman, Nathan D. Lord, Johan Paulsson, and Richard Losick. Stochastic Switching of Cell Fate in Microbes. Annual Review of Microbiology, 69(1):381–403, 2015. doi: 10.1146/annurev-micro-091213-112852.

6. Harsh Vashistha, Maryam Kohram, and Hanna Salman. Non-genetic inheritance restraint of cell-to-cell variation. eLife, 10:e64779, 2021. doi: 10.7554/eLife.64779.

7. Anna Kuchina, Lorena Espinar, Tolga Çağ atay, Alejandro O Balbin, Fang Zhang, Alma Alvarado, Jordi Garcia-Ojalvo, and Gürol M Süel. Temporal competition between differentiation programs determines cell fate choice. Molecular Systems Biology, 7(1):557, 2011. doi: 10.1038/msb.2011.88.

8. Joe H. Levine, Michelle E. Fontes, Jonathan Dworkin, and Michael B. Elowitz. Pulsed Feedback Defers Cellular Differentiation. PLOS Biology, 10(1):e1001252, 2012. doi: 10.1371/journal.pbio.1001252.

9. Omri Gilhar, Liat Rahamim Ben-Navi, Tsviya Olender, Asaph Aharoni, Jonathan Friedman, and Ilana Kolodkin-Gal. Multigenerational inheritance drives symbiotic interactions of the bacterium Bacillus subtilis with its plant host. Microbiological Research, 286:127814, 2024. doi: 10.1016/j.micres.2024.127814.

10. Lam Vo, Fotios Avgidis, Henry H. Mattingly, Karah Edmonds, Isabel Burger, Ravi Balasubramanian, Thomas S. Shimizu, Barbara I. Kazmierczak, and Thierry Emonet. Nongenetic adaptation by collective migration. Proceedings of the National Academy of Sciences, 122 (8):e2423774122, 2025. doi: 10.1073/pnas.2423774122.

11. Sydney M. Shaffer, Benjamin L. Emert, Raúl A. Reyes Hueros, Christopher Cote, Guillaume Harmange, Dylan L. Schaff, Ann E. Sizemore, Rohit Gupte, Eduardo Torre, Abhyudai Singh, Danielle S. Bassett, and Arjun Raj. Memory Sequencing Reveals Heritable Single-Cell Gene Expression Programs Associated with Distinct Cellular Behaviors. Cell, 182(4):947–959.e17, 2020. doi: 10.1016/j.cell.2020.07.003.

12. Michael J. Pokrass, Kathleen A. Ryan, Tianchi Xin, Brittany Pielstick, Winston Timp, Valentina Greco, and Sergi Regot. Cell-Cycle-Dependent ERK Signaling Dynamics Direct Fate Specification in the Mammalian Preimplantation Embryo. Developmental Cell, 55(3): 328–340.e5, 2020. doi: 10.1016/j.devcel.2020.09.013.

13. Maroš Pleška, David Jordan, Zak Frentz, BingKan Xue, and Stanislas Leibler. Nongenetic individuality, changeability, and inheritance in bacterial behavior. Proceedings of the National Academy of Sciences, 118(13):e2023322118, 2021. doi: 10.1073/pnas.2023322118.

14. Arnab Bandyopadhyay, Huijing Wang, and J. Christian J. Ray. Lineage space and the propensity of bacterial cells to undergo growth transitions. PLOS Computational Biology, 14(8):e1006380, 2018. doi: 10.1371/journal.pcbi.1006380.

15. Dann Huh and Johan Paulsson. Non-genetic heterogeneity from stochastic partitioning at cell division. Nature Genetics, 43(2):95–100, 2011. doi: 10.1038/ng.729.

16. Dann Huh and Johan Paulsson. Random partitioning of molecules at cell division. Proceedings of the National Academy of Sciences, 108(36):15004–15009, 2011. doi: 10.1073/pnas.1013171108.

17. Adam James Waite, Nicholas W. Frankel, and Thierry Emonet. Behavioral Variability and Phenotypic Diversity in Bacterial Chemotaxis. Annual Review of Biophysics, 47(Volume 47, 2018):595–616, 2018. doi: 10.1146/annurev-biophys-062215-010954.

18. Evren U. Azeloglu and Ravi Iyengar. Signaling Networks: Information Flow, Computation, and Decision Making. Cold Spring Harbor Perspectives in Biology, 7(4):a005934, 2015. doi: 10.1101/cshperspect.a005934.

19. J. Christian J. Ray, Jeffrey J. Tabor, and Oleg A. Igoshin. Non-transcriptional regulatory processes shape transcriptional network dynamics. Nature Reviews Microbiology, 9(11): 817–828, 2011. doi: 10.1038/nrmicro2667.

20. Andrew A. Bridges, Jojo A. Prentice, Ned S. Wingreen, and Bonnie L. Bassler. Signal Transduction Network Principles Underlying Bacterial Collective Behaviors. Annual Review of Microbiology, 76(1):235–257, 2022. doi: 10.1146/annurev-micro-042922-122020.

21. Sahand Hormoz, Nicolas Desprat, and Boris I. Shraiman. Inferring epigenetic dynamics from kin correlations. Proceedings of the National Academy of Sciences, 112(18):E2281–E2289, 2015. doi: 10.1073/pnas.1504407112.

22. Pedro Pessoa, Juan Andres Martinez, Vincent Vandenbroucke, Frank Delvigne, and Steve Pressé. Inherited or produced? Inferring protein production kinetics when protein counts are shaped by a cell’s division history. 2025. doi: 10.48550/arXiv.2506.09374.

23. Nitzan Rosenfeld, Jonathan W. Young, Uri Alon, Peter S. Swain, and Michael B. Elowitz. Gene Regulation at the Single-Cell Level. Science, 307(5717):1962–1965, 2005. doi: 10.1126/science.1106914.

24. Ido Golding, Johan Paulsson, Scott M. Zawilski, and Edward C. Cox. Real-Time Kinetics of Gene Activity in Individual Bacteria. Cell, 123(6):1025–1036, 2005. doi: 10.1016/j.cell.2005.09.031.

25. Franz M. Weinert, Robert C. Brewster, Mattias Rydenfelt, Rob Phillips, and Willem K. Kegel. Scaling of Gene Expression with Transcription-Factor Fugacity. Physical Review Letters, 113(25):258101, 2014. doi: 10.1103/PhysRevLett.113.258101.

26. Brian Munsky, Gregor Neuert, and Alexander van Oudenaarden. Using Gene Expression Noise to Understand Gene Regulation. Science, 336(6078):183–187, 2012. doi: 10.1126/science.1216379.

27. Daniel L. Jones, Robert C. Brewster, and Rob Phillips. Promoter architecture dictates cell-to-cell variability in gene expression. Science, 346(6216):1533–1536, 2014. doi: 10.1126/science.1255301.

28. Douglas E. Weidemann, James Holehouse, Abhyudai Singh, Ramon Grima, and Silke Hauf. The minimal intrinsic stochasticity of constitutively expressed eukaryotic genes is sub-Poissonian. Science Advances, 9(32):eadh5138, 2023. doi: 10.1126/sciadv.adh5138.

29. D. R. Cox. Some Statistical Methods Connected with Series of Events. Journal of the Royal Statistical Society: Series B (Methodological), 17(2):129–157, 1955. doi: 10.1111/j.2517-6161.1955.tb00188.x.

30. Bharath Sunchu and Clemens Cabernard. Principles and mechanisms of asymmetric cell division. Development, 147(13):dev167650, 2020. doi: 10.1242/dev.167650.

31. Ali Kinkhabwala, Anton Khmelinskii, and Michael Knop. Analytical model for macromolecular partitioning during yeast cell division. BMC Biophysics, 7(1):10, 2014. doi: 10.1186/s13628-014-0010-6.

32. Matthias Christen, Hemantha D. Kulasekara, Beat Christen, Bridget R. Kulasekara, Lucas R. Hoffman, and Samuel I. Miller. Asymmetrical Distribution of the Second Messenger c-di-GMP upon Bacterial Cell Division. Science, 328(5983):1295–1297, 2010. doi: 10.1126/science.1188658.

33. Johan Paulsson. Models of stochastic gene expression. Physics of Life Reviews, 2(2): 157–175, 2005. doi: 10.1016/j.plrev.2005.03.003.

34. Eran Eden, Naama Geva-Zatorsky, Irina Issaeva, Ariel Cohen, Erez Dekel, Tamar Danon, Lydia Cohen, Avi Mayo, and Uri Alon. Proteome Half-Life Dynamics in Living Human Cells. Science, 331(6018):764, 2011. doi: 10.1126/science.1199784.

35. Bryson D Bennett, Elizabeth H Kimball, Melissa Gao, Robin Osterhout, Stephen J Van Dien, and Joshua D Rabinowitz. Absolute metabolite concentrations and implied enzyme active site occupancy in Escherichia coli. Nature Chemical Biology, 5(8):593–599, 2009. doi: 10.1038/nchembio.186.

36. Hugo Dourado, Matteo Mori, Terence Hwa, and Martin J. Lercher. On the optimality of the enzyme–substrate relationship in bacteria. PLOS Biology, 19(10):e3001416, 2021. doi: 10.1371/journal.pbio.3001416.

37. Regine Hengge, Angelika Gründling, Urs Jenal, Robert Ryan, and Fitnat Yildiz. Bacterial Signal Transduction by Cyclic Di-GMP and Other Nucleotide Second Messengers. Journal of Bacteriology, 198(1):15–26, 2016. doi: 10.1128/JB.00331-15.

38. P. A. P. Moran. NOTES ON CONTINUOUS STOCHASTIC PHENOMENA. Biometrika, 37 (1-2):17–23, 1950. doi: 10.1093/biomet/37.1-2.17.

39. Caelan Brooks, Meiyi Yao, Jake T. McCool, Alan Gillman, Gürol M. Süel, Andrew Mugler, and Joseph W. Larkin. Computational model of fractal interface formation in bacterial biofilms. Physical Review E, 112(6):064408, 2025. doi: 10.1103/2zm9-r3qs.

40. Harry Nunns and Lea Goentoro. Signaling pathways as linear transmitters. eLife, 7:e33617, 2018. doi: 10.7554/eLife.33617.

41. Nicholas E. Phillips, Aleksandra Mandic, Saeed Omidi, Felix Naef, and David M. Suter. Memory and relatedness of transcriptional activity in mammalian cell lineages. Nature Communications, 10(1):1208, 2019. doi: 10.1038/s41467-019-09189-8.

42. Michael L. Simpson, Chris D. Cox, Michael S. Allen, James M. McCollum, Roy D. Dar, David K. Karig, and John F. Cooke. Noise in biological circuits. Wiley Interdisciplinary Reviews: Nanomedicine and Nanobiotechnology, 1(2):214–225, 2009. doi: 10.1002/wnan.22.

43. Johan Paulsson, Otto G. Berg, and Måns Ehrenberg. Stochastic focusing: Fluctuation-enhanced sensitivity of intracellular regulation. Proceedings of the National Academy of Sciences, 97(13):7148–7153, 2000. doi: 10.1073/pnas.110057697.

44. Mark D. McDonnell and Derek Abbott. What Is Stochastic Resonance? Definitions, Misconceptions, Debates, and Its Relevance to Biology. PLOS Computational Biology, 5(5): e1000348, 2009. doi: 10.1371/journal.pcbi.1000348.

45. Johan Paulsson. Summing up the noise in gene networks. Nature, 427(6973):415–418, 2004. doi: 10.1038/nature02257.

46. Juan M. Pedraza and Alexander van Oudenaarden. Noise Propagation in Gene Networks. Science, 307(5717):1965–1969, 2005. doi: 10.1126/science.1109090.

47. Dionysios Barmpoutis and Richard M. Murray. Noise Propagation in Biological and Chemical Reaction Networks. 2011. doi: 10.48550/arXiv.1108.2538.

48. Christina Kim, Gregory J. Seedorf, Steven H. Abman, and Douglas P. Shepherd. Heterogeneous response of endothelial cells to insulin-like growth factor 1 treatment is explained by spatially clustered sub-populations. Biology Open, 8(11):bio045906, 2019. doi: 10.1242/bio.045906.

49. Karl Francis and Bernhard O. Palsson. Effective intercellular communication distances are determined by the relative time constants for cyto/chemokine secretion and diffusion. Proceedings of the National Academy of Sciences, 94(23):12258–12262, 1997. doi: 10.1073/pnas.94.23.12258.

50. Jordi van Gestel, Tasneem Bareia, Bar Tenennbaum, Alma Dal Co, Polina Guler, Nitzan Aframian, Shani Puyesky, Ilana Grinberg, Glen G. D’Souza, Zohar Erez, Martin Ackermann, and Avigdor Eldar. Short-range quorum sensing controls horizontal gene transfer at micron scale in bacterial communities. Nature Communications, 12(1):2324, 2021. doi: 10.1038/s41467-021-22649-4.

51. Simon van Vliet, Christoph Hauert, Kyle Fridberg, Martin Ackermann, and Alma Dal Co. Global dynamics of microbial communities emerge from local interaction rules. PLOS Computational Biology, 18(3):e1009877, 2022. doi: 10.1371/journal.pcbi.1009877.

52. Junchao Shi, Qi Chen, Xin Li, Xiudeng Zheng, Ying Zhang, Jie Qiao, Fuchou Tang, Yi Tao, Qi Zhou, and Enkui Duan. Dynamic transcriptional symmetry-breaking in pre-implantation mammalian embryo development revealed by single-cell RNA-seq. Development, 142(20): 3468–3477, 2015. doi: 10.1242/dev.123950.

53. Zibo Chen and Michael B. Elowitz. Programmable protein circuit design. Cell, 184(9):2284–2301, 2021. doi: 10.1016/j.cell.2021.03.007.

54. Xiaoyu Yang, Jason W. Rocks, Kaiyi Jiang, Andrew J. Walters, Kshitij Rai, Jing Liu, Jason Nguyen, Scott D. Olson, Pankaj Mehta, James J. Collins, Nichole M. Daringer, and Caleb J. Bashor. Engineering synthetic phosphorylation signaling networks in human cells. Science, 387(6729):74–81, 2025. doi: 10.1126/science.adm8485.

55. Alma Dal Co, Simon van Vliet, and Martin Ackermann. Emergent microscale gradients give rise to metabolic cross-feeding and antibiotic tolerance in clonal bacterial populations. Philosophical Transactions of the Royal Society B: Biological Sciences, 374(1786):20190080, 2019. doi: 10.1098/rstb.2019.0080.

56. Daniel T. Gillespie. Exact stochastic simulation of coupled chemical reactions. The Journal of Physical Chemistry, 81(25):2340–2361, 1977. doi: 10.1021/j100540a008.

57. H E Kubitschek and J A Friske. Determination of bacterial cell volume with the Coulter Counter. Journal of Bacteriology, 168(3):1466–1467, 1986. doi: 10.1128/jb.168.3.1466-1467.1986.

58. Charles R. Harris, K. Jarrod Millman, Stéfan J. van der Walt, Ralf Gommers, Pauli Virtanen, David Cournapeau, Eric Wieser, Julian Taylor, Sebastian Berg, Nathaniel J. Smith, Robert Kern, Matti Picus, Stephan Hoyer, Marten H. van Kerkwijk, Matthew Brett, Allan Haldane, Jaime Fernández del Río, Mark Wiebe, Pearu Peterson, Pierre Gérard-Marchant, Kevin Sheppard, Tyler Reddy, Warren Weckesser, Hameer Abbasi, Christoph Gohlke, and Travis E. Oliphant. Array programming with NumPy. Nature, 585(7825):357–362, 2020. doi: 10.1038/s41586-020-2649-2.

59. Pauli Virtanen, Ralf Gommers, Travis E. Oliphant, Matt Haberland, Tyler Reddy, David Cournapeau, Evgeni Burovski, Pearu Peterson, Warren Weckesser, Jonathan Bright, Stéfan J. van der Walt, Matthew Brett, Joshua Wilson, K. Jarrod Millman, Nikolay Mayorov, Andrew R. J. Nelson, Eric Jones, Robert Kern, Eric Larson, C. J. Carey, İlhan Polat, Yu Feng,Eric W. Moore, Jake VanderPlas, Denis Laxalde, Josef Perktold, Robert Cimrman, Ian Henriksen, E. A. Quintero, Charles R. Harris, Anne M. Archibald, Antônio H. Ribeiro, Fabian Pedregosa, and Paul van Mulbregt. SciPy 1.0: fundamental algorithms for scientific computing in Python. Nature Methods, 17(3):261–272, 2020. doi: 10.1038/s41592-019-0686-2.

60. John D. Hunter. Matplotlib: A 2D Graphics Environment. Computing in Science & Engineering, 9(3):90–95, 2007. doi: 10.1109/MCSE.2007.55.

61. Gary Bradski. The Opencv Library. Dr. Dobb’s Journal of Software Tools, 120:122–125, 2000.

62. Robert Foreman and Roy Wollman. Mammalian gene expression variability is explained by underlying cell state. Molecular Systems Biology, 16(2):e9146, 2020. doi: 10.15252/msb.20199146.

63. Dan Davidi, Elad Noor, Wolfram Liebermeister, Arren Bar-Even, Avi Flamholz, Katja Tummler, Uri Barenholz, Miki Goldenfeld, Tomer Shlomi, and Ron Milo. Global characterization of in vivo enzyme catalytic rates and their correspondence to in vitro k _cat_measurements. Proceedings of the National Academy of Sciences, 113(12):3401–3406, 2016. doi:10.1073/pnas.1514240113.

64. Boumediene Soufi, Karsten Krug, Andreas Harst, and Boris Macek. Characterization of the E. coli proteome and its modifications during growth and ethanol stress. Frontiers in Microbiology, 6, 2015.

65. R. Milo, S. Shen-Orr, S. Itzkovitz, N. Kashtan, D. Chklovskii, and U. Alon. Network Motifs: Simple Building Blocks of Complex Networks. Science, 298(5594):824–827, 2002. doi:10.1126/science.298.5594.824.

66. Uri Alon. Network motifs: theory and experimental approaches. Nature Reviews Genetics, 8(6):450–461, 2007. doi: 10.1038/nrg2102.

67. John J. Tyson and Béla Novák. Functional Motifs in Biochemical Reaction Networks. Annual Review of Physical Chemistry, 61(1):219–240, 2010. doi: 10.1146/annurev.physchem.012809.103457.

68. Mathieu Cloutier and Edwin Wang. Dynamic modeling and analysis of cancer cellular network motifs. Integrative Biology, 3(7):724–732, 2011. doi: 10.1039/c0ib00145g.

69. Long Cai, Nir Friedman, and X. Sunney Xie. Stochastic protein expression in individual cells at the single molecule level. Nature, 440(7082):358–362, 2006. doi: 10.1038/nature04599.

70. Daniel Zenklusen, Daniel R. Larson, and Robert H. Singer. Single-RNA counting reveals alternative modes of gene expression in yeast. Nature Structural & Molecular Biology, 15 (12):1263–1271, 2008. doi: 10.1038/nsmb.1514.

71. Alexander M Jones, Jonas ÅH Danielson, Shruti N ManojKumar, Viviane Lanquar, Guido Grossmann, and Wolf B Frommer. Abscisic acid dynamics in roots detected with genetically encoded FRET sensors. eLife, 3:e01741, 2014. doi: 10.7554/eLife.01741.

72. Stephen Cooper and Charles E. Helmstetter. Chromosome replication and the division cycle of Escherichia coli Br. Journal of Molecular Biology, 31(3):519–540, 1968. doi: 10.1016/0022-2836(68)90425-7.

73. Suckjoon Jun, Fangwei Si, Rami Pugatch, and Matthew Scott. Fundamental principles in bacterial physiology—history, recent progress, and the future with focus on cell size control: a review. Reports on Progress in Physics, 81(5):056601, 2018. doi: 10.1088/1361-6633/aaa628.

74. R. G. Eagon. Pseudomonas natriegens, a marine bacterium with a generation time of less than 10 minutes. Journal of Bacteriology, 83(4):736–737, 1962. doi: 10.1128/jb.83.4.736-737.1962.

75. Xiongfeng Dai, Zichu Shen, Yiheng Wang, and Manlu Zhu. Sinorhizobium meliloti, a Slow-Growing Bacterium, Exhibits Growth Rate Dependence of Cell Size under Nutrient Limitation. mSphere, 3(6):e00567–18, 2018. doi: 10.1128/mSphere.00567-18.

76. M. Schuster, M. L. Urbanowski, and E. P. Greenberg. Promoter specificity in Pseudomonas aeruginosa quorum sensing revealed by DNA binding of purified LasR. Proceedings of the National Academy of Sciences, 101(45):15833–15839, 2004. doi: 10.1073/pnas.0407229101.

77. Jae Kyoung Kim and John J. Tyson. Misuse of the Michaelis–Menten rate law for protein interaction networks and its remedy. PLOS Computational Biology, 16(10):e1008258, 2020. doi: 10.1371/journal.pcbi.1008258.

78. Werner Sandmann. Sequential estimation for prescribed statistical accuracy in stochastic simulation of biological systems. Mathematical Biosciences, 221(1):43–53, 2009. doi: 10.1016/j.mbs.2009.06.006.

79. Philipp Thomas, Hannes Matuschek, and Ramon Grima. How reliable is the linear noise approximation of gene regulatory networks? BMC Genomics, 14(4):S5, 2013. doi: 10.1186/1471-2164-14-S4-S5.

80. James E. Ferrell and Sang Hoon Ha. Ultrasensitivity part III: cascades, bistable switches, and oscillators. Trends in Biochemical Sciences, 39(12):612–618, 2014. doi: 10.1016/j.tibs.2014.10.002.

81. A Goldbeter and D E Koshland. An amplified sensitivity arising from covalent modification in biological systems. Proceedings of the National Academy of Sciences, 78(11):6840–6844, 1981. doi: 10.1073/pnas.78.11.6840.

82. Carlos Gomez-Uribe, George C. Verghese, and Leonid A. Mirny. Operating Regimes of Signaling Cycles: Statics, Dynamics, and Noise Filtering. PLOS Computational Biology, 3 (12):e246, 2007. doi: 10.1371/journal.pcbi.0030246.

83. Huijing Wang and J. Christian J. Ray. Dynamical predictors of an imminent phenotypic switch in bacteria. Physical Biology, 14(4):045007, 2017. doi: 10.1088/1478-3975/aa7870.

84. Ann M. Stock, Victoria L. Robinson, and Paul N. Goudreau. Two-Component Signal Transduction. Annual Review of Biochemistry, 69(Volume 69, 2000):183–215, 2000. doi: 10.1146/annurev.biochem.69.1.183.

85. Rong Gao and Ann M. Stock. Biological Insights from Structures of Two-Component Proteins. Annual Review of Microbiology, 63(Volume 63, 2009):133–154, 2009. doi: 10.1146/annurev.micro.091208.073214.

86. Felipe Trajtenberg, Juan A Imelio, Matías R Machado, Nicole Larrieux, Marcelo A Marti, Gonzalo Obal, Ariel E Mechaly, and Alejandro Buschiazzo. Regulation of signaling directionality revealed by 3D snapshots of a kinase:regulator complex in action. eLife, 5:e21422, 2016. doi: 10.7554/eLife.21422.

87. Eric Batchelor and Mark Goulian. Robustness and the cycle of phosphorylation and dephosphorylation in a two-component regulatory system. Proceedings of the National Academy of Sciences, 100(2):691–696, 2003. doi: 10.1073/pnas.0234782100.

88. Sheng Jian Cai and Masayori Inouye. EnvZ-OmpR Interaction and Osmoregulation in Escherichia coli *. Journal of Biological Chemistry, 277(27):24155–24161, 2002. doi: 10.1074/jbc.M110715200.

89. Tim Miyashiro and Mark Goulian. High stimulus unmasks positive feedback in an autoregulated bacterial signaling circuit. Proceedings of the National Academy of Sciences, 105 (45):17457–17462, 2008. doi: 10.1073/pnas.0807278105.

90. Won-Sik Yeo, Igor Zwir, Henry V. Huang, Dongwoo Shin, Akinori Kato, and Eduardo A. Groisman. Intrinsic Negative Feedback Governs Activation Surge in Two-Component Regulatory Systems. Molecular Cell, 45(3):409–421, 2012. doi: 10.1016/j.molcel.2011.12.027.

