## Supplemental Figures and Appendices for "Memory from variability: Heritable short-term cellular memory emerges from stochastic biochemical reaction networks"

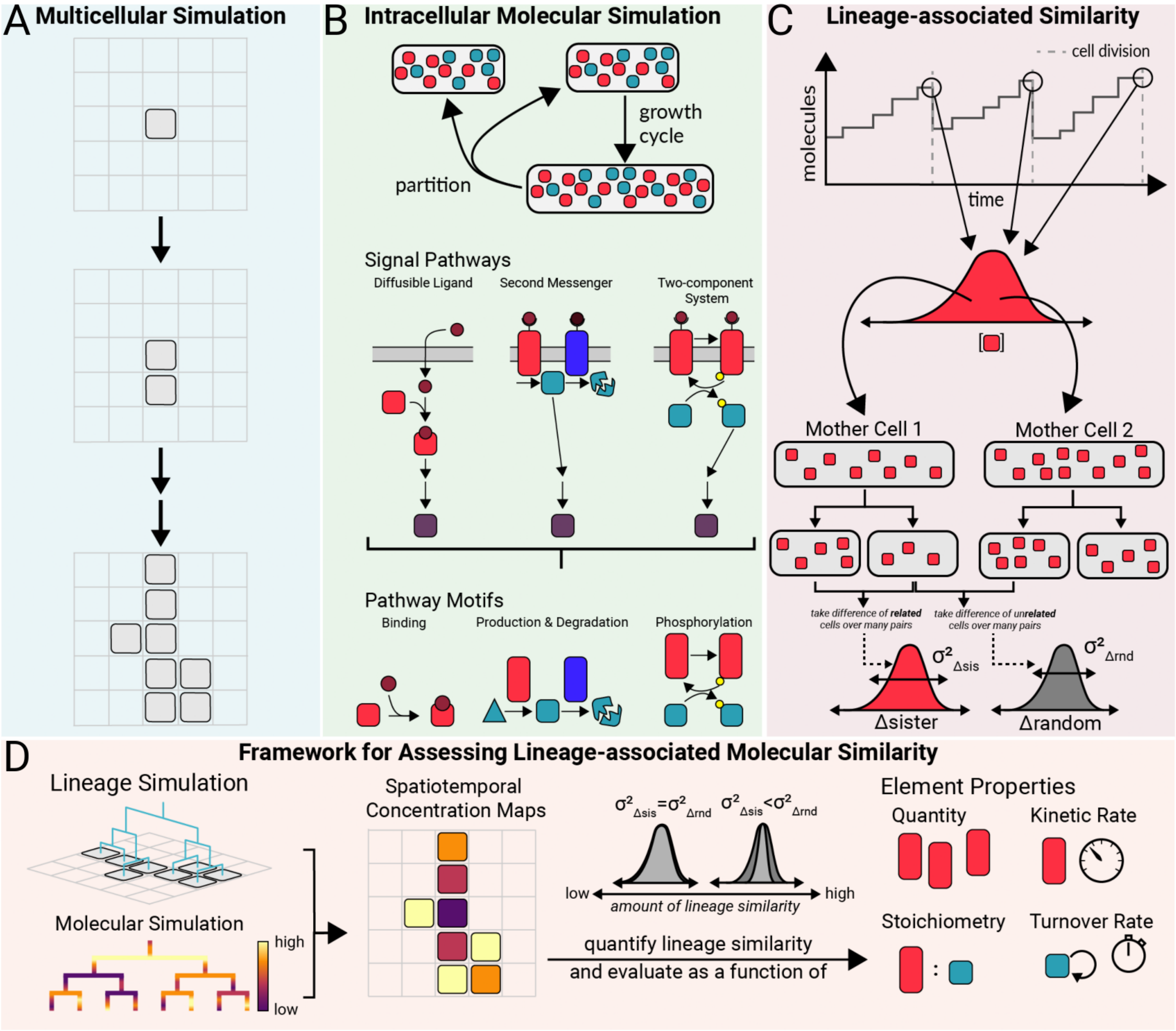


**Fig. S1. Framework for lineage-associated similarity | (A)** Simulations describe a single cell growing into a small multicellular collective over time. **(B)** During the multicellular simulation, cells grow, producing more molecules, and, upon division, partition the molecules between the resulting daughter cells. The amount of molecules in each cell is dictated by signaling pathway kinetics, which consist of individual pathway motifs. **(C)** Calculation of the lineage-associated similarity metric. **(D)** The complete framework determines kinetic factors that influence the magnitude and duration of lineage-associated similarity.

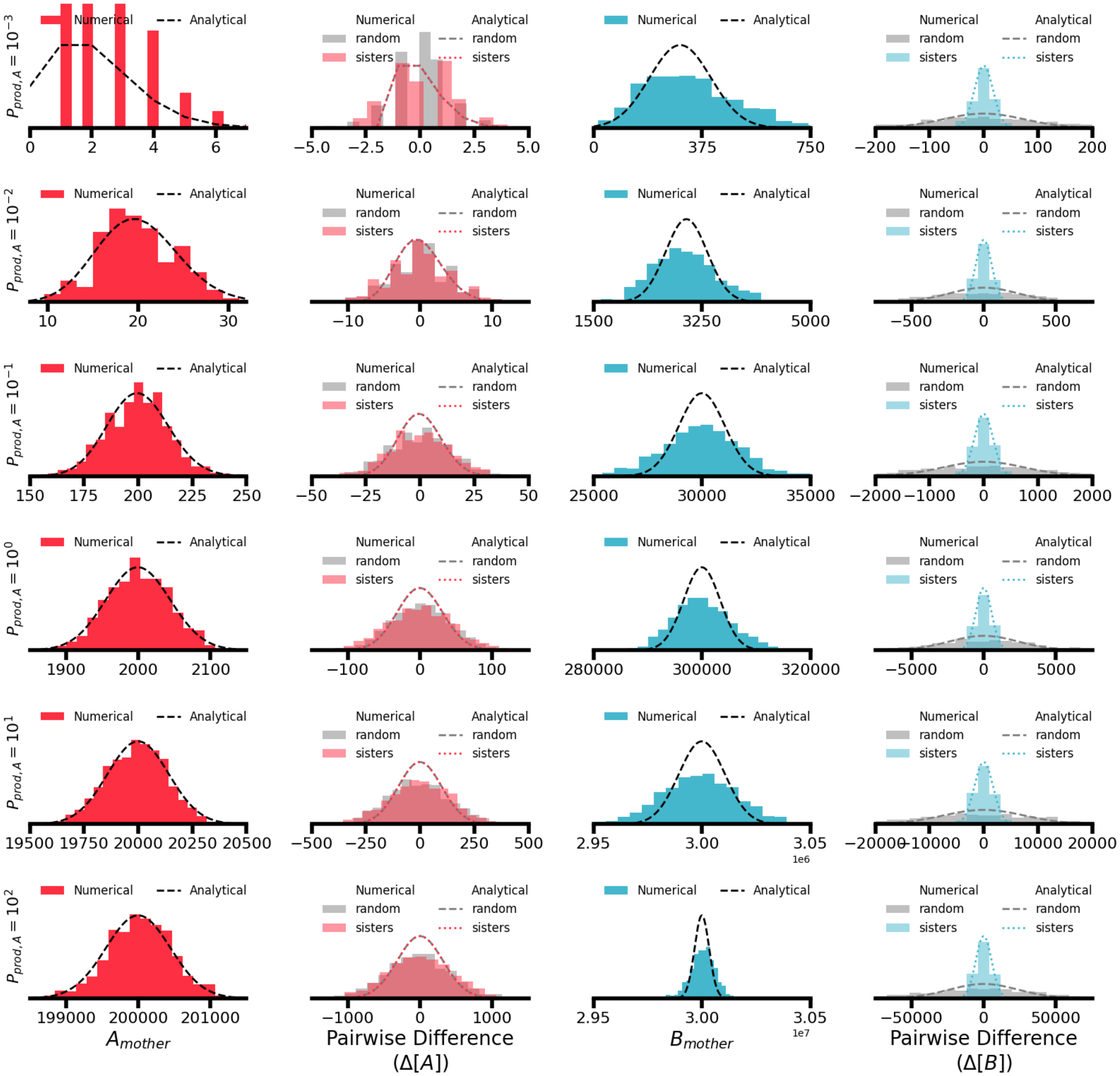


**Fig. S2. Distributions of molecules in the saturated production motif over a range of enzyme amounts | First column:** Number of enzyme A molecules in mother cells across six orders of magnitude of production probabilities (rows), fit to a Poisson distribution (black dashed line). **Second column:** Difference distributions of enzyme A molecules between random pairs (gray) and sister pairs (pink), along with Poisson fits (dashed and dotted lines). **Third column:** Number of product molecule B in mother cells, fit to a negative binomial distribution (black dashed line). **Fourth column:** Difference distributions of product molecule B between random pairs (gray) and sister pairs (cyan). A negative binomial distribution parameterized with the analytically derived mean and variance is used as an estimate of the distribution of differences in molecule B (dashed lines). See SI Appendix G for all simulation parameter values.

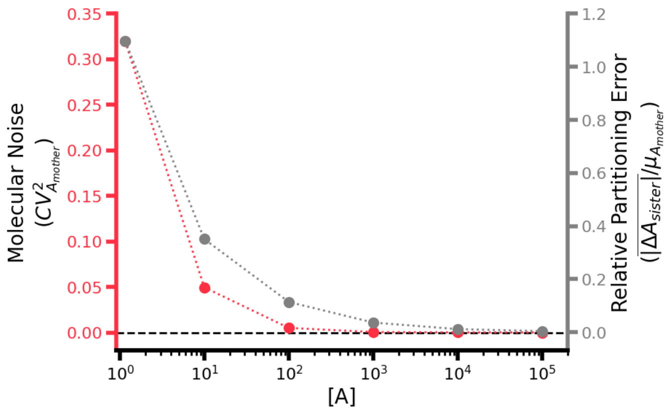


**Fig. S3. Relative errors that arise during a single cell generation and partitioning scale with molecule amount** | As the amount of molecule A increases (due to an increasing production rate $P_{prod,A}$), the molecular noise in the concentration of A (as measured by the coefficient of variation squared, left axis) and the relative partition error (as measured by the mean of the absolute value of the difference in A between sister cells scaled by the average amount of A, right axis) both decrease. This result explains how, even in "low-noise regimes" with a high level of molecule A, molecule A does not exhibit any lineage-associated similarity. See SI Appendix G for all simulation parameter values.

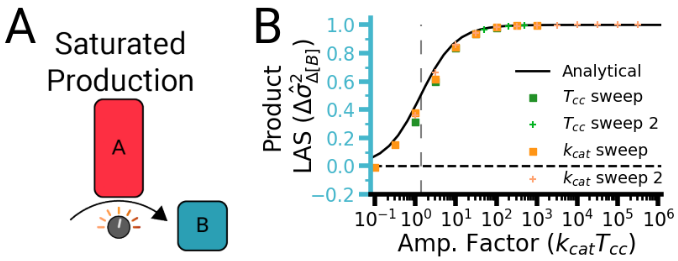


**Fig. S4. Full amplification factor sweeps** | **(A)** Schematic of the saturated production motif. **(B)** Lineage-associated similarity measure for the product molecule B as a function of the amplification factor ($k_{cat,A}T_{cc}$). Initial sweeps ($T_{cc}$ sweep and $k_{cat,A}$ sweep) were performed with values lower than the standard parameter values to fully explore the analytical prediction (black line). Subsequent sweeps ($T_{cc}$ sweep 2 and $k_{cat,A}$ sweep 2) were performed with standard parameter values for the remainder of the simulations (see SI Appendix C, Part 1). See SI Appendix G for all simulation parameter values.

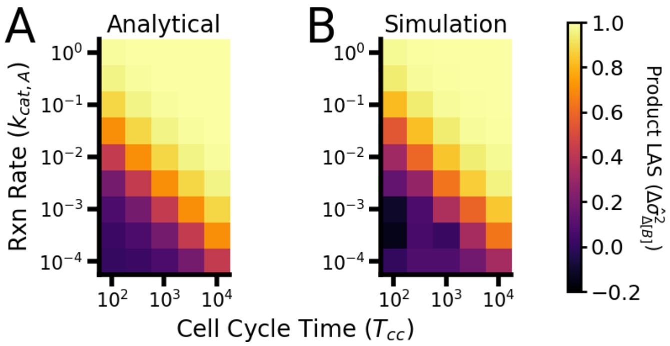


**Fig. S5. Comparison of the extent of lineage-associated similarity as a function of parameters for the saturated production circuit** | **(A)** Predicted extent of lineage-associated similarity for the product molecule B across different cell cycle times (x-axis) and maximal reaction rates (y-axis). **(B)** Numerically calculated extent of lineage-associated similarity for different amounts of the producing enzyme A. The extent of lineage-associated similarity is independent of the amount of A and is instead a function of reactions per cell cycle, a product of the reaction rate $k_{cat,A}$ and cell cycle time $T_{cc}$. See SI Appendix G for all simulation parameter values.

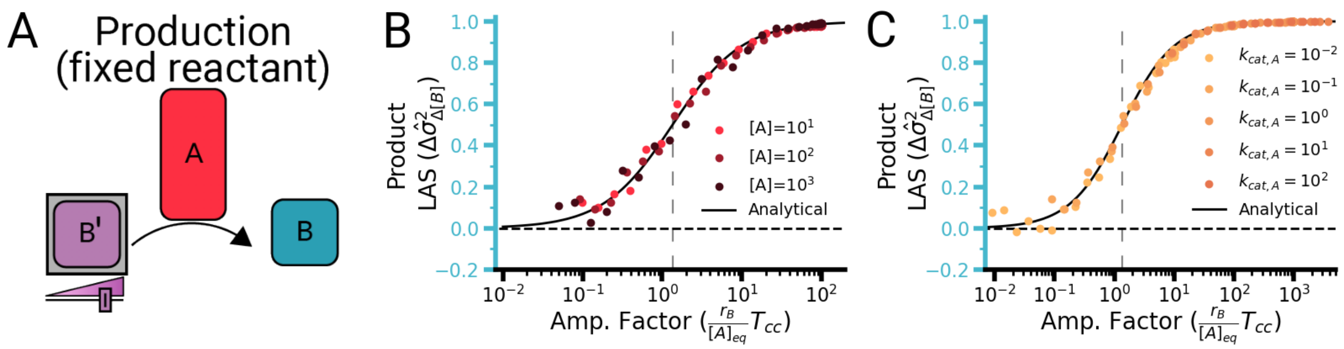


**Fig. S6. Amplification factor sweeps for the production motif with fixed reactant** | **(A)** Schematic depicting the production motif with fixed reactant, in which a constant pool of reactant ($B'$) is converted into a product molecule ($B$) by an enzyme ($A$). **(B)** Lineage-associated similarity of the product molecule ($B$) quantified using the normalized difference of variances (Equation 1) as a function of the amplification factor and different enzyme concentrations (different shades of red dots). The solid black line is the analytical solution (Equation 6), and the vertical dashed line indicates the inflection point of the analytical solution. **(C)** Lineage-associated similarity of the product molecule ($B$) quantified using the normalized difference of variances (Equation 1) as a function of the amplification factor and different maximal reaction rates (different shades of orange dots). The solid black line is the analytical solution (Equation 6), and the vertical dashed line indicates the inflection point of the analytical solution. n=1000 pairs of cells for each similarity calculation. See SI Appendix G for all simulation parameter values.

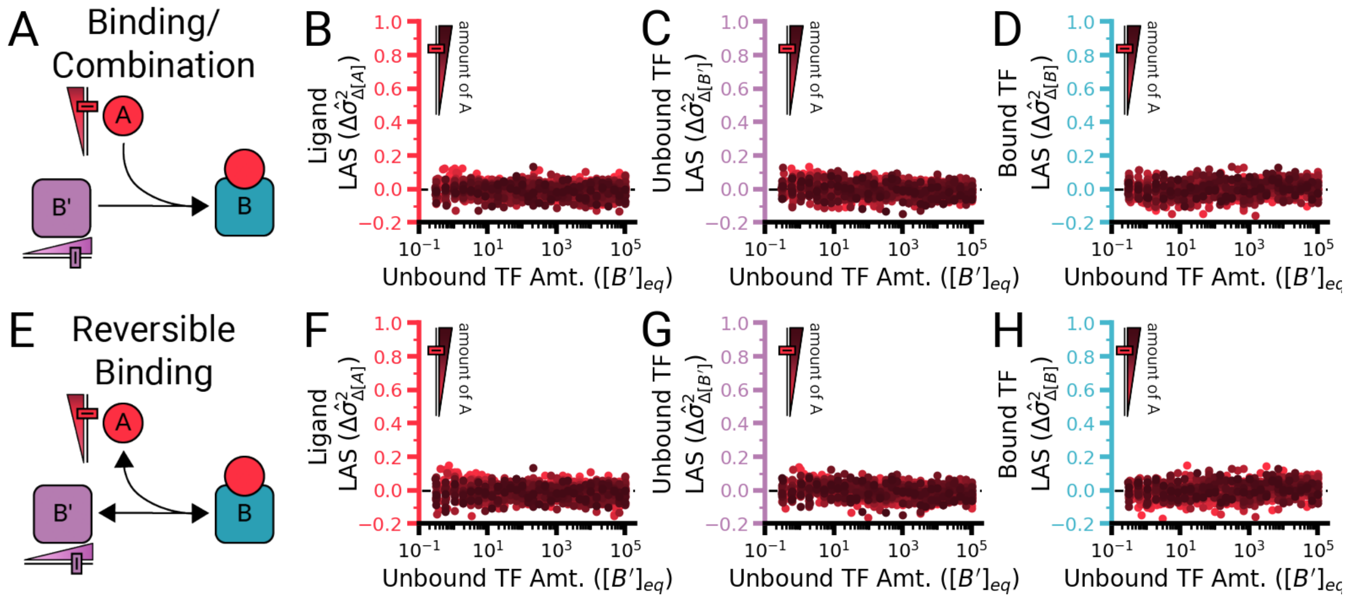


**Fig. S7. Nonsaturating binding motifs do not produce lineage-associated similarity | (A)** Schematic illustrating the irreversible binding motif, in which a ligand ($A$) binds to an unbound transcription factor ($B'$) to produce a bound transcription factor complex ($B$). **(B–D)** Lineage-associated similarity of the irreversible binding molecules (**(B)** ligand $A$, **(C)** unbound transcription factor (TF) $B'$, **(D)** bound transcription factor $B$) over orders of magnitude of unbound transcription factor (x-axis) and ligand (different colors). **(E)** Schematic of the reversible binding motif, in which the bound transcription factor complex ($B$) can unbind, restoring free ligand ($A$) and unbound transcription factor ($B'$). **(E–G)** Lineage-associated similarity of all molecules in the reversible binding motif over various amounts of unbound transcription factor (x-axis) and ligand (different colored scatter plots). See SI Appendix G for all simulation parameter values.

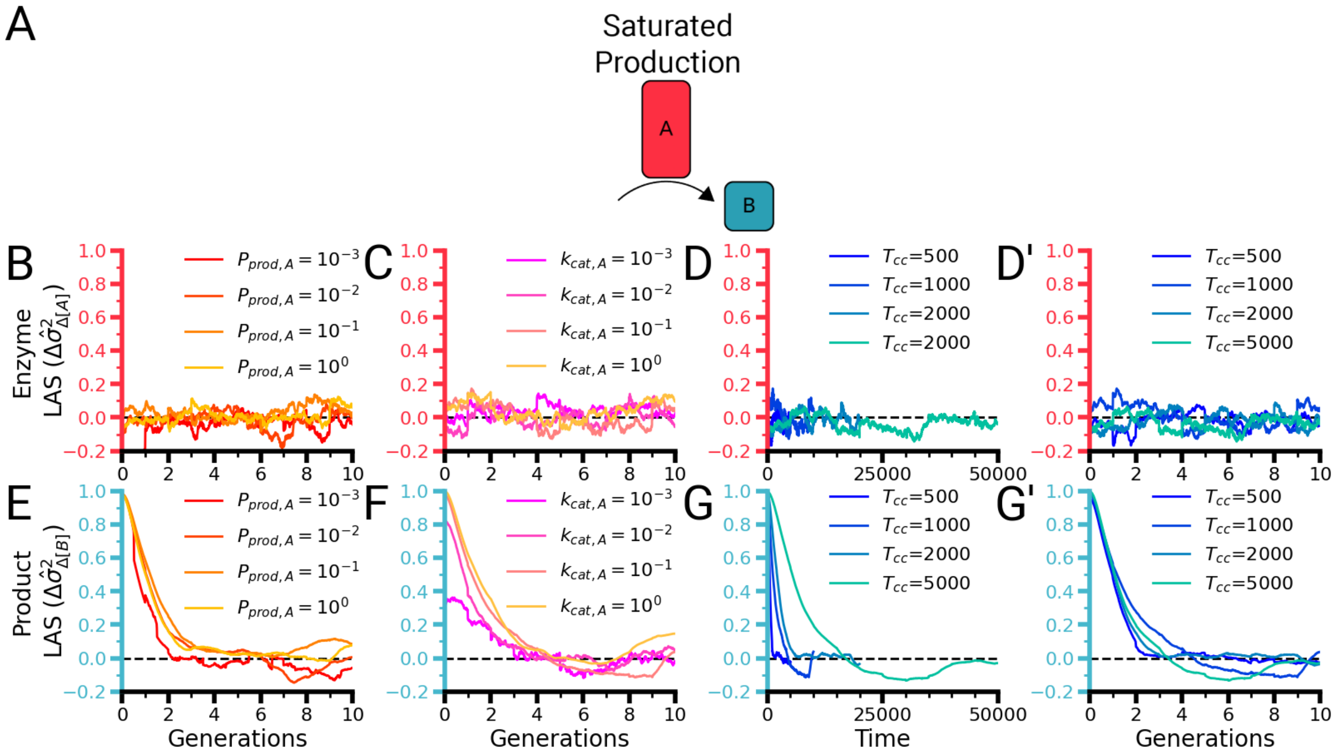

**Fig. S8. Duration of lineage-associated similarity in the production-only motif is consistent across all parameter ranges | (A)** Schematic of the production motif. An enzyme B converts a substrate molecule A into a product molecule B. **(B)** Normalized difference of variances for the concentration of enzyme A over time across varying production rates of A. **(C)** Normalized difference of variances for the concentration of enzyme A over time across varying reaction rates of A. **(D)** Normalized difference of variances for the concentration of the enzyme A over time across varying cell cycle times. **(E)** Normalized difference of variances for the concentration of the product molecule B over time across varying production rates of A. **(F)** Normalized difference of variances for the concentration of the product molecule B over time across varying reaction rates of A. **(G)** Normalized difference of variances for the concentration of the product molecule B over time across varying cell cycle times. **(D')** Normalized difference of variances for the concentration of enzyme A over time across varying cell cycles normalized to the cell cycle number. **(G')** Normalized difference of variances for the concentration of product molecule B over time across varying cell cycles normalized to the cell cycle number. While the lineage-associated similarity duration is different in absolute time, it remains the same in generational time. See SI Appendix G for all simulation parameter values.

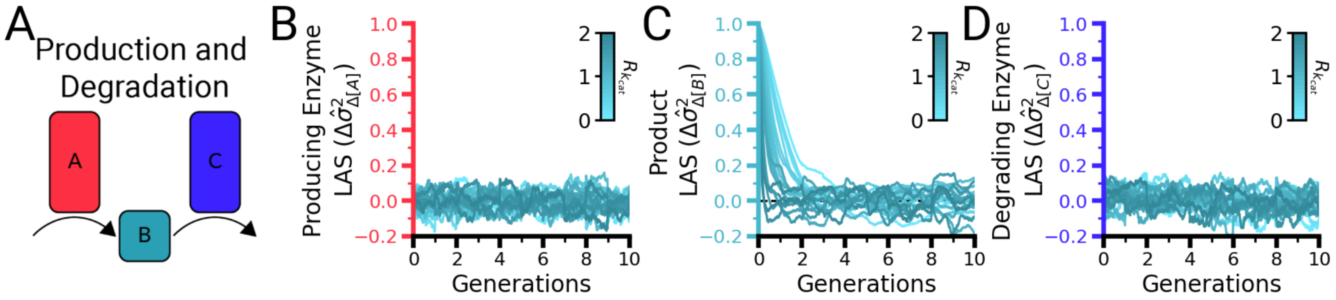


**Fig. S9. Full lineage-associated similarity data for the $R_{k_{cat}}$ parameter sweep** | **(A)** Schematic of the production and degradation motif, in which a producing enzyme A generates a product molecule B at saturation while a degrading enzyme C degrades B by Michaelis–Menten kinetics. **(B)** Normalized difference of variance of producing enzyme A across all values of $R_{k_{cat}}$. **(C)** Normalized difference of variance of product molecule B across all values of $R_{k_{cat}}$. As the value of $R_{k_{cat}}$ increases, the molecular turnover rate increases and the duration of lineage-associated similarity in the concentration of B decreases. **(D)** Normalized difference of variance of degrading enzyme C across all values of $R_{k_{cat}}$. See SI Appendix G for all simulation parameter values.

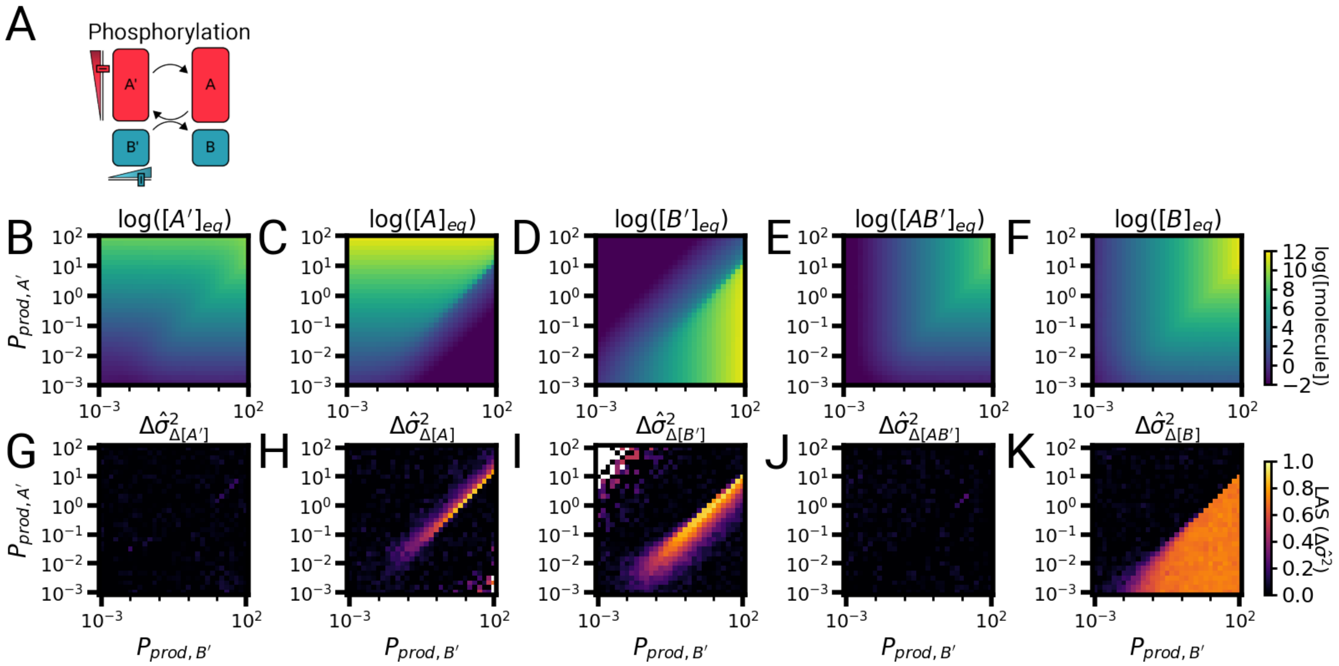


**Fig. S10. Lineage-associated similarity is present in a phosphorylation motif at saturation | (A)** Schematic of the phosphorylation motif with monofunctional sensor histidine kinase. The sensor histidine kinase ($A'$) autophosphorylates, producing phosphorylated sensor histidine kinase ($A$), which then binds to the unphosphorylated response regulator ($B'$), forming a complex ($AB'$) before transphosphorylating to produce phosphorylated response regulator ($B$). **(B–F)** Concentrations of each component molecule across the parameter sweep of production probability of the sensor histidine kinase ($P_{prod,A'}$, y-axis) and response regulator ($P_{prod,B'}$, x-axis). **(G–K)** Lineage-associated similarity of each component of the phosphorylation motif across a production probability parameter sweep. Note that the blank pixels in the heatmaps represent parameter combinations that resulted in a division error when the lineage-associated similarity was calculated. See SI Appendix G for all simulation parameter values.

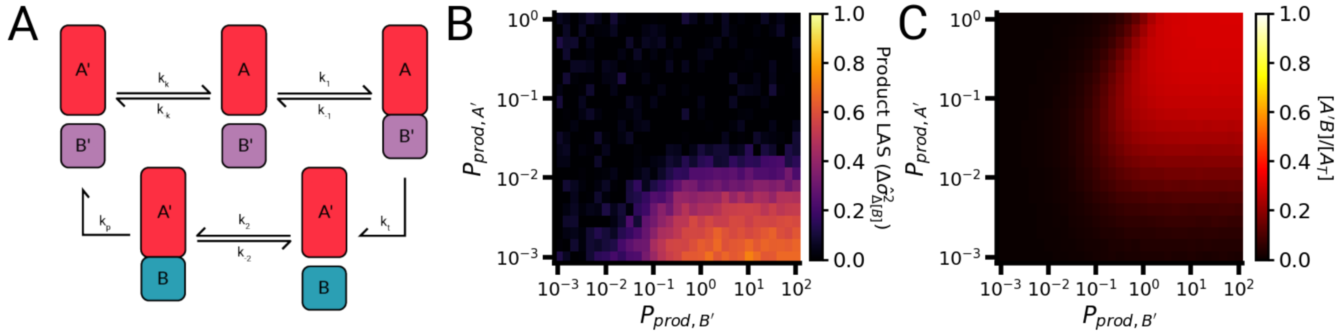


**Fig. S11. Phosphorylation motif with dual kinase–phosphatase activity displays saturation-dependent lineage-associated similarity and product quenching** | **(A)** Schematic of the phosphorylation motif with dual kinase–phosphatase activity. Unphosphorylated sensor histidine kinase ($A'$) autophosphorylates to produce phosphorylated sensor histidine kinase ($A$). Phosphorylated sensor histidine kinase ($A$) then binds with unphosphorylated response regulator ($B'$) to produce the bound complex $A'B$. Transphosphorylation produces unphosphorylated sensor histidine kinase ($A'$) and phosphorylated response regulator ($B$). These species then bind to produce the bound unphosphorylated sensor histidine kinase and phosphorylated response regulator complex ($AB'$), which, upon phosphatase activity, reproduces unphosphorylated sensor histidine kinase ($A'$) and unphosphorylated response regulator ($B'$). **(B)** Normalized difference of variances for phosphorylated response regulator ($B$) over various amounts of total sensor histidine kinase ($P_{prod,A}$) and response regulator ($P_{prod,B}$). **(C)** Fraction of sensor histidine kinase found in the bound unphosphorylated sensor histidine kinase–phosphorylated response regulator ($A'B$) complex. See SI Appendix G for all simulation parameter values.

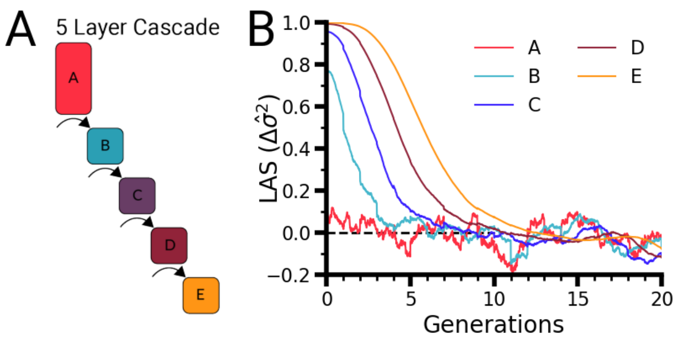


**Fig. S12. Additional cascade layers add generations of similarity** | **(A)** Schematic of a five-layer cascade, where each layer catalyzes the formation of the next layer (SI Appendix C.15). **(B)** Lineage-associated similarity of each molecule in the cascade over time. Each layer adds an additional 2–3 generations of similarity, leading to a total of ten generations of similarity in the final molecule E. Note that, due to the increasing layers, this motif is computationally expensive and was therefore run with parameters smaller than the values used throughout the remainder of this work. See SI Appendix G for all simulation parameter values.

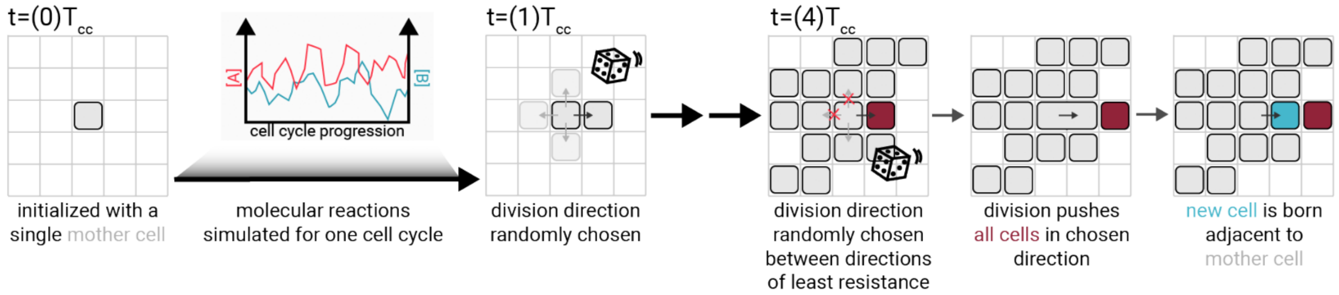


**Fig. S13. Visualization of the spatial simulation algorithm** | Spatial simulations are seeded with a single mother cell. Between divisions, the molecular reactions are simulated with the same Gillespie algorithm as before (see SI Appendix C.16). When the cell generation completes, a division direction is randomly selected out of the four cardinal directions. After many rounds of division ($t = (4)T_{cc}$), a small colony has formed. Now upon division, the division direction is selected from among the cardinal directions with the lowest number of obstructing cells (here, the up and left directions have two cells and are eliminated in favor of the right or down directions, which have only one cell. The division direction is then randomly chosen from the remaining directions). Upon division, the cell "elongates" and pushes all cells (red cell) in that direction to make room for the new daughter cell (cyan cell). This same algorithm is repeated for every cell division during the time course of the simulation.

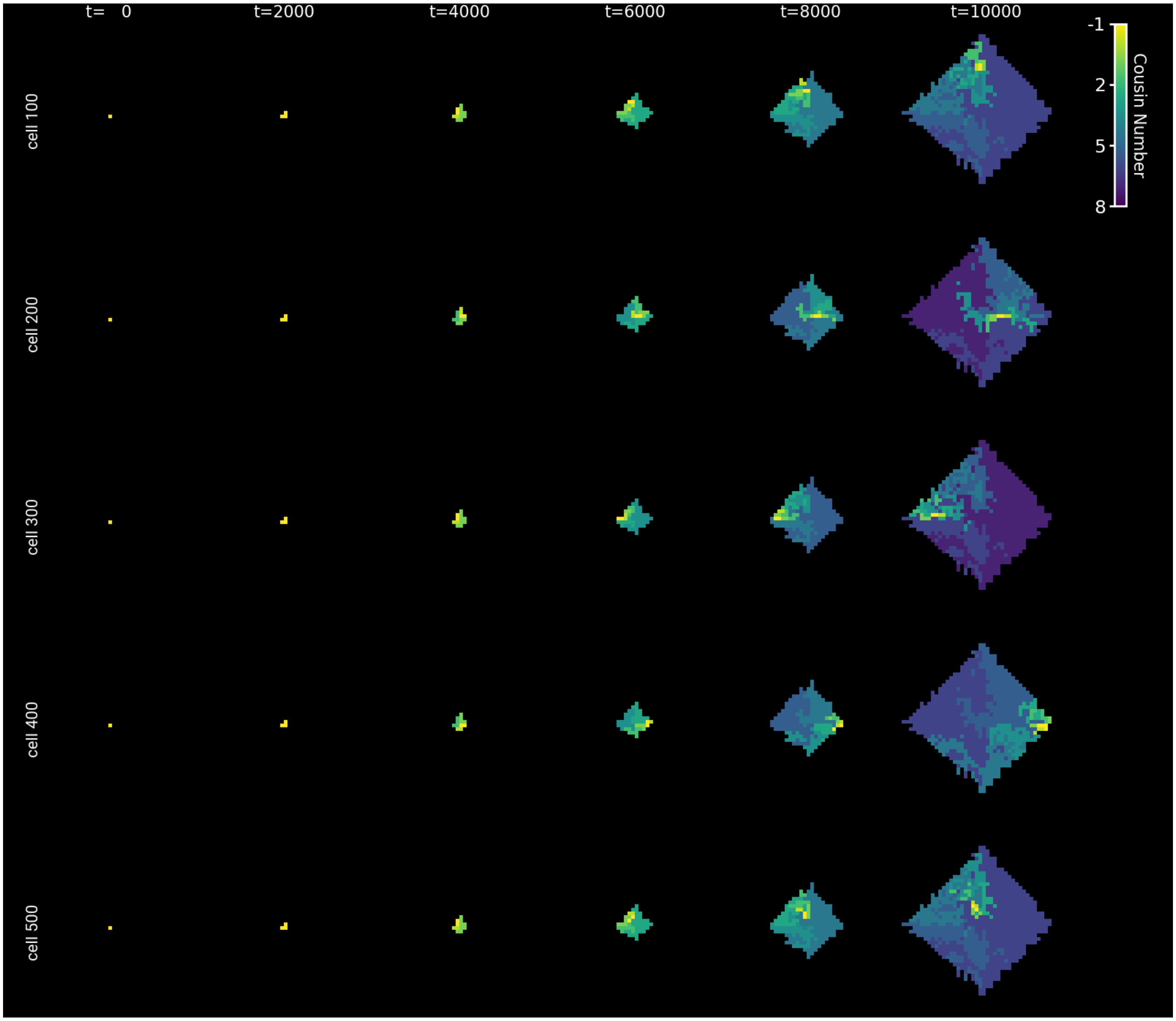


**Fig. S14. Cousin maps for different reference cells** | A cousin map is calculated as a function of simulation time and a reference cell. All other cells are colored based on their cousin number with respect to the reference cell. Here, multiple reference cells are used to generate different cousin maps (rows) over simulation time (columns).

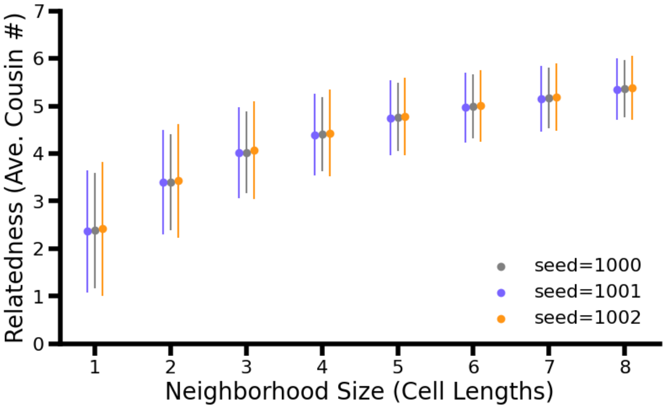


**Fig. S15. Relatedness curves are consistent over different spatial simulations** | Average cousin number as a function of neighborhood relatedness for three different spatial simulations seeded with different random seeds. The relationship between neighborhood size and relatedness is well conserved over different seeds of the spatial simulation. n=1000 cells for the analyzed frame of each simulation.

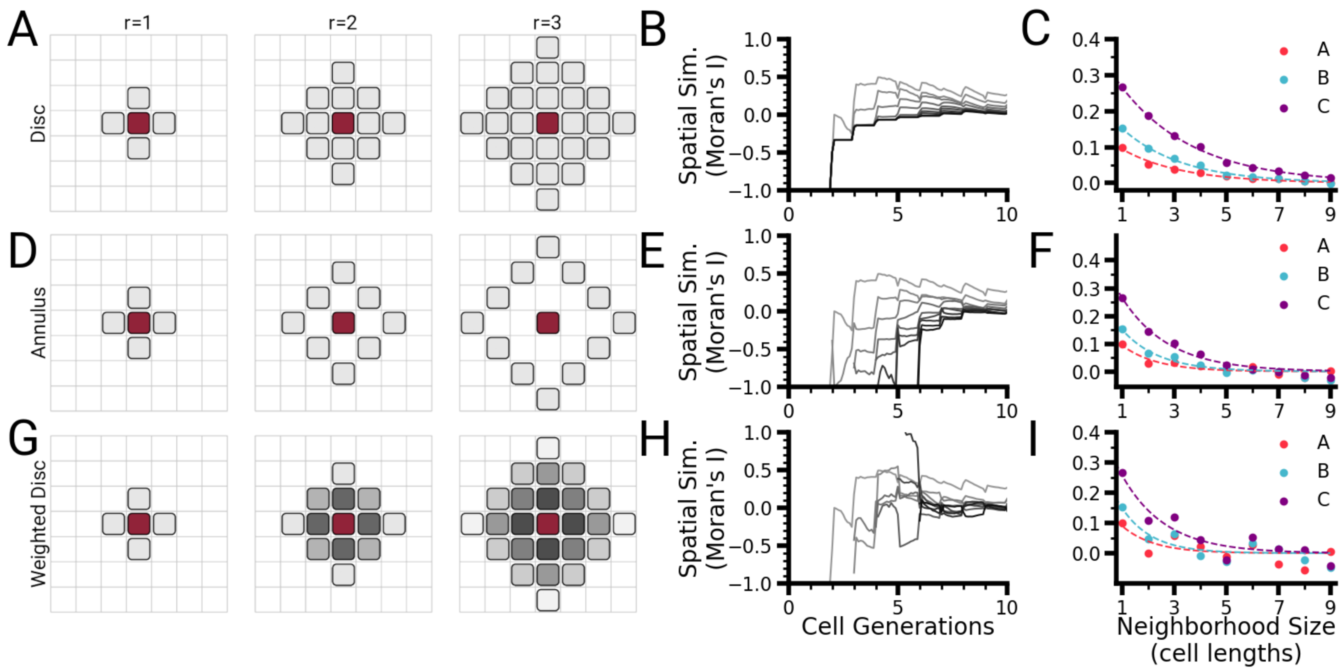


**Fig. S16. A binary disc provides the most effective neighborhood weight definition for spatial autocorrelation calculations** | **(A)** Schematic depicting the definition of the binary disc neighborhood as a function of neighborhood radius for a reference cell (red cell). **(B)** Moran's I quantification as a function of time for different radius values (different gray lines). After the colony grows to a sufficient size, the values level out. **(C)** Moran's I as a function of neighborhood radius for the final time point of the simulation for the saturated production motif. Lines are exponential fits intended as guides for the eye. **(D)** Schematic depicting the definition of the binary annulus neighborhood as a function of neighborhood radius for a reference cell (red cell). **(E)** Moran's I quantification as a function of time for different radius values (different gray lines). Similar to the disc scenario, as the colony reaches a sufficient size, the values level out. **(F)** Moran's I as a function of neighborhood radius for the final time point of the simulation. Product molecule B has a higher value than enzyme A, but the fits are not as cohesive as those observed for the disc definition. **(G)** Schematic depicting the definition of the weighted disc neighborhood as a function of neighborhood radius for a reference cell (red cell). While the shape is the same as the binary disc (row 1), the values placed on each neighbor are a function of their distance from the reference cell. **(H)** Moran's I as a function of time (x-axis) and neighborhood radius (gray lines) for the weighted disc neighborhood. **(I)** Moran's I as a function of neighborhood radius for the final time point of the simulation for the weighted disc neighborhood. Values are noisier with respect to exponential fits than either the disc or annulus definition. n=1000 cells in the final time point of the simulation. See SI Appendix G for all simulation parameter values.

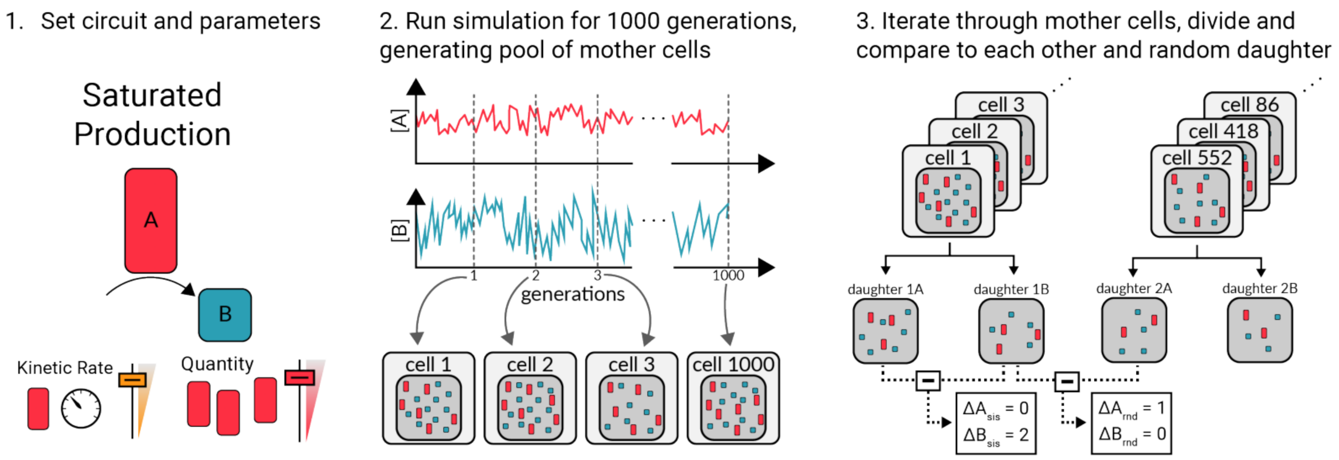


**Fig. S17. Seed cell simulation schematic** | To generate the populations required for making cell comparisons, the following algorithm is followed: (1) A circuit and parameters are chosen. (2) A single cell is simulated for 1000 generations, generating 1000 possible mother cells with different combinations of molecules. (3) The 1000 possible mother cells (Step 3, left) are paired with a randomly selected mother cell from the same pool (Step 3, right). The first mother cell is divided, producing two daughter cells. The difference in molecule number between the two daughters is calculated for each molecule ($\Delta A_{sis}$ and $\Delta B_{sis}$). The random mother cell is also divided, and the difference in molecule numbers between a daughter from the first cell and one of the daughter cells from the random mother is calculated ($\Delta A_{rnd}$ and $\Delta B_{rnd}$).

#### SI Appendix A: Lineage-associated similarity metrics

Throughout the main text, the metrics used to quantify lineage-associated similarity originate from distributions of comparisons of molecule concentrations between sister and random pairs of cells (Figure 2B and C for the variance of pairwise differences and Figures 2 F-I, 3A, and 4 for the LAS index). We further extended the use of the LAS index when analyzing the temporal dynamics of this similarity (Figures 3C and D, 5, and 6C). This new lineage-associated similarity metric was used because it mirrors the exact question we are interested in: are related cells more similar (i.e. have a smaller variance in pairwise differences) than random cells? However, we also find it useful to consider the LAS index's relationship with the Pearson correlation coefficient, as the Pearson correlation coefficient has been used by others to consider if related cells are similar (6). The relationship between our metric for lineage-associated similarity and the Pearson correlation coefficient is evident from a closer examination of their mathematical expressions. The full expression for the LAS index of molecule A over time can be written as follows:

$$\Delta\hat{\sigma}^2_{\Delta[A]}(t) = \frac{\sigma^2_{\Delta rnd}(t) - \sigma^2_{\Delta sis}(t)}{\sigma^2_{\Delta rnd}(t)} \quad \textbf{(8)}$$

$$= 1 - \frac{\sigma^2_{\Delta sis}(t)}{\sigma^2_{\Delta rnd}(t)} \quad \textbf{(9)}$$

$$= 1 - \frac{\sum_{i=0}^{n=N}([A]_{rel1,i,t} - [A]_{rel2,i,t})^2}{\sum_{i=0}^{n=N}([A]_{rel1,i,t} - [A]_{rnd,i,t})^2}, \quad \textbf{(10)}$$

where $N$ is the total number of cell pairs examined (for most of the calculations present in this paper, 1000 pairs each of related and unrelated cells). $[A]_{rel1,i,t}$ is the concentration of molecule A in the first cell of related pair $i$ at time $t$, $[A]_{rel2,i,t}$ is the concentration of molecule A in the second sister cell of pair $i$ at time $t$, and $[A]_{rnd,i,t}$ is the concentration of molecule A in an unrelated cell of pair $i$ at time $t$. Note that we compare related cells and not sister cells in the time-series version of this metric because in subsequent generations, we are comparing cousins, second cousins, etc.

The time-series Pearson correlation coefficient indicates if two quantities vary together (covariance) in a unitless normalized metric. In the context of this metric, similarity is if amount of molecule A varies together in related cells. The Pearson coefficient for related cells can be compared to the coefficient for random cells to assess if related cells behave differently than random cells. Specifically, the Pearson correlation coefficient should be computed for the amount of molecule A in related cells as follows:

$$r_{[A],sis}(t) = \frac{\sum_{i=0}^{n=N}([A]_{rel1,i,t} - [\bar{A}]_{rel1,t})([A]_{rel2,i,t} - [\bar{A}]_{rel2,t})}{N\sqrt{\frac{\sum_{i=0}^{n=N}([A]_{rel1,i,t} - [\bar{A}]_{rel1,t})^2}{N}\frac{\sum_{i=0}^{n=N}([A]_{rel2,i,t} - [\bar{A}]_{rel2,t})^2}{N}}}. \quad \textbf{(11)}$$

In this expression, $[\bar{A}]_{rel1,t}$ is the average concentration of A across all cells designated as cell one in pairs of related cells at time $t$, and $[\bar{A}]_{rel2,t}$ is the average concentration of A across all cells designated as cell two in pairs of related cells at time $t$.

Both our lineage-associated similarity metric (Figure S18F) and the Pearson correlation coefficient for related cells (Figure S18G) capture the decay of similarity in a molecular species over time in related cells.

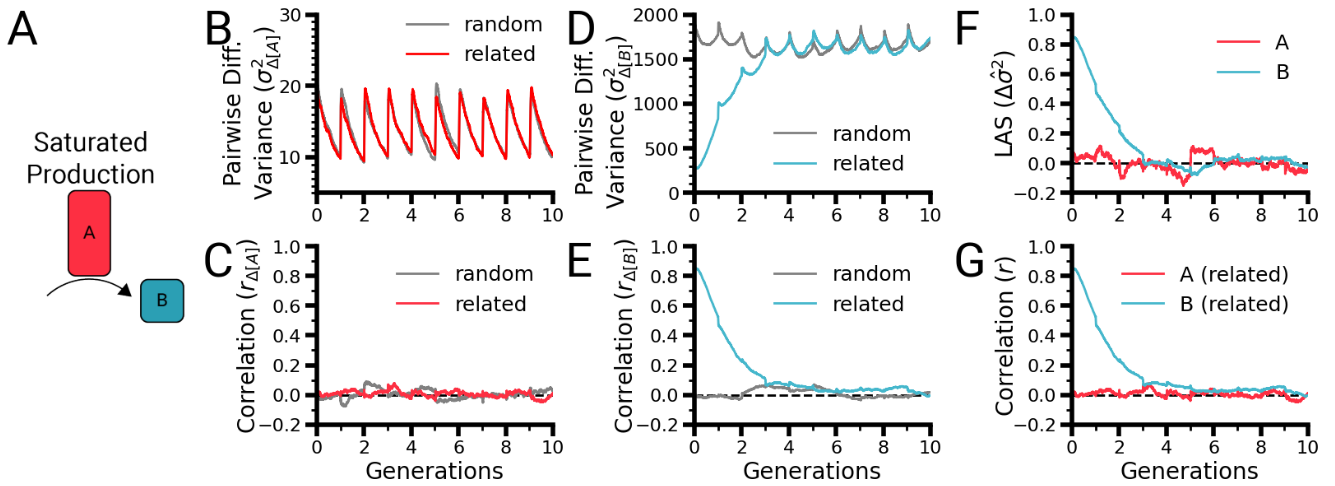


**Fig. S18. Comparison between lineage-associated similarity metrics** | **(A)** Schematic of the saturated production motif used for metric comparison analysis. We simulated 1000 pairs of cells with the saturated production motif for ten generations. The molecular concentration data over time for related pairs and random pairs of cells for enzyme A were analyzed using the variance of pairwise differences as shown in Figure 2B and C **(B)** and the time-series Pearson correlation coefficient **(C)**. Similarly, the concentration traces of the product molecule B were analyzed using the variance of pairwise differences **(D)** and the time-series Pearson correlation coefficient **(E)**. **(F)** and **(G)** show the lineage-associated similarity of each molecule over ten generations, with the lineage-associated similarity metric used in **(F)** and the Pearson correlation coefficient for related pairs of cells used in **(G)**. See SI Appendix G for all simulation parameter values.

#### SI Appendix B: Conceptual framework for how molecular levels are modulated in cells during growth and division over multiple generations

To model how molecular concentrations are shaped during a single cell generation and then affected by cell division, we use the following simple and experimentally informed rules:

1. Biochemical species in our framework, such as proteins and small molecules, are produced in two possible ways:
   (a) Biochemical species are consumed and new molecules are produced by a reaction. These molecules are created with a production probability dictated by kinetic biochemical reaction equations that we have parameterized with rate constants ($k_{cat}$ and similar rate constants) typical of bacteria (see SI Appendix C.1).
   (b) All other molecules are produced with a fixed probability (i.e., a Poisson process with a rate parameter denoted as $P_{prod}$). This model is appropriate for the production of molecules and is in line with experimental measurements for many gene products (26, 27, 62).
2. Cells grow at a constant rate over a preset cell cycle time (set by the parameter $T_{cc}$), during which molecules are produced and consumed as described by Rule 1.
3. Cells divide physically in half, and each molecular species is binomially partitioned (p = 0.5) between the daughter cells.

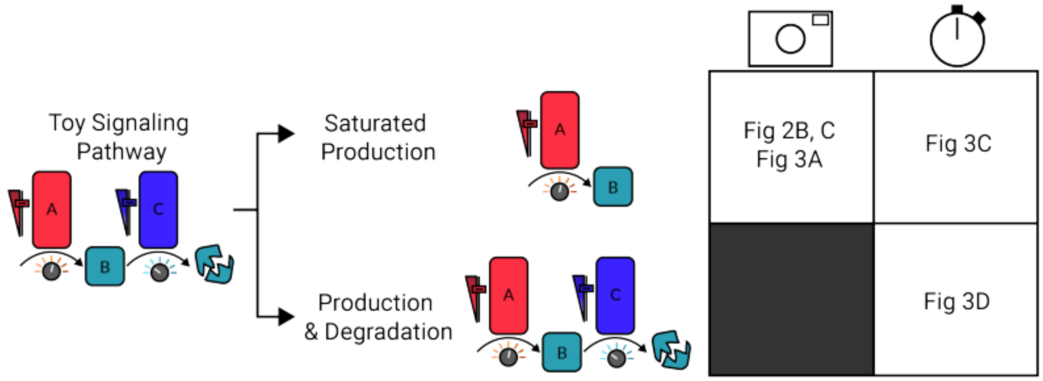


**Fig. S19. Toy signaling pathway model breakout and locations** | To gain an intuition about the signaling pathway properties that lead to lineage-associated similarity, we started with a toy signaling pathway, where an enzyme A produces a product molecule B, which is then degraded by a second enzyme C. We simulated two forms of the pathway: a saturated production circuit, which examines only the production of molecule B by enzyme A, and the full production & degradation version, which includes the degradation by molecule C. For saturated production, we examined snapshot analyses in Figures 2 and 3 and temporal analyses in Figure 3. For the production & degradation version, we examined temporal analyses in Figure 3.

To finalize this framework, we must identify the basic biochemical reactions that compose complex signaling networks so that we can fully explore how different signaling network architectures may impact similarity. Sufficient data are available for bacteria and their signaling pathways to build and parameterize the models of (27, 63, 64). The unbranched structures of common bacterial signaling pathways (20) make them conducive to building simple models for analyzing inheritance patterns. Furthermore, some major mammalian signaling pathways exhibit input–output linearity over biologically realistic parameter ranges (40). Taken together, these linear bacterial pathways represent potentially good approximations of more complex mammalian regulatory networks, allowing our results to be generalized across kingdoms of life. We considered three critical classes of bacterial intracellular signaling pathways (20): a diffusible ligand circuit, a second messenger system, and a two-component system. A network analysis approach (65–68) suggests that these pathways can be decomposed into five biochemical reaction motifs shared across the pathways: gene expression, binding, production, degradation, and phosphorylation. By mathematically modeling each of these reaction motifs, we can fully explore how these basic biochemical reactions affect molecular concentrations of related cells down cellular lineages.

To extend our framework beyond the first two generations (mother and daughters), after the division creating the first two daughter cells, we continued to simulate the same intracellular reactions and additional divisions for ten generations.

#### SI Appendix C: Simulation model construction

**Part 1: Parameter ranges.** In the simplest state, a producing enzyme is expressed by a constitutive promoter and, while this promoter is in an active state, transcript production is stochastic (69). mRNA transcripts have been shown to follow Poisson statistics (70), and for many constitutive promoters, this description can also be applied to gene production (26, 71). We parameterize the system by setting a production rate from the enzyme promoter, $P_{prod,A}$, the cell cycle duration, and the maximal reaction rate of the enzyme A, $k_{cat,A}$. The steady-state concentration of enzyme A is simply the product of the production rate and the cell cycle time (SI Appendix C.3). We set the cell cycle time to 1000 time units, comparable to the number of seconds in the *E. coli* cell cycle during exponential growth (72). With this setup, our model assumes balanced growth for all signaling molecules considered (73). To determine if the signal pathway property of quantity affects lineage-associated similarity (Figure S1, Element Properties), we scaled the production rate such that the protein concentration spans from $10^0$ to $10^5$ nM, matching the range of protein copy numbers in *E. coli* (64). Rate constants of $10^{-3} - 10^3 s^{-1}$ were analyzed, in line with typical $k_{cat}$ values for enzymes from *E. coli* (63). For cell cycle times, we took inspiration from *Pseudomonas natriegens*, the fastest recorded growing bacteria, with a cell cycle time under 10 min (74), and set the lower bound of $T_{cc}$ to 500 s. For the upper bound, we were inspired by the 2- to 6-h doubling time of *Sinorhizobium meliloti* (75) and set the maximum screened value of $T_{cc}$ to 10,000 s.

**Part 2: Saturated production motif.** In the saturated production motif (Figures 2 and 3), A is the enzyme that produces the product molecule B.

$$prod_A = P_{prod,A}, \tag{1}$$
$$prod_B = k_{cat,A}A, \tag{2}$$
$$R_{total} = prod_A + prod_B, \tag{3}$$

where $P_{prod,A}$ is the production rate of enzyme A and $k_{cat,A}$ is the maximal reaction rate of enzyme A.

**Part 3: Production motif with fixed reactant.** In the production motif with fixed reactant, a reactant molecule $B'$ is converted into a product molecule $B$ by an enzyme $A$. One of the parameters of this simulation is a specified concentration of reactant $B'$. At each time step in the simulation, the reactant amount is set by this parameter multiplied by the volume of the cell at that time step. This amount of reactant is then used for the Gillespie algorithm:

$$prod_A = P_{prod,A}, \tag{4}$$
$$prod_B = \frac{k_{cat,A}}{2}\left(K_M + A + B' - \sqrt{(K_M + A + B')^2 - 4AB'}\right), \tag{5}$$
$$R_{total} = prod_A + prod_B, \tag{6}$$

where $P_{prod,A}$ is the production probability of enzyme A, $k_{cat,A}$ is the maximal reaction rate of enzyme A, and $K_M$ is the half-saturation constant.

**Part 4: Production motif.** In the production motif (shown in Figure 4), $A$ is the enzyme, $B'$ is the reactant, and $B$ is the product molecule.

$$prod_A = P_{prod,A}, \tag{7}$$
$$prod_{B'} = P_{prod,B'}, \tag{8}$$
$$prod_B = \frac{k_{cat,A}}{2}\left(K_M + A + B' - \sqrt{(K_M + A + B')^2 - 4AB'}\right), \tag{9}$$
$$R_{total} = prod_A + prod_{B'} + prod_B, \tag{10}$$

where $P_{prod,A}$ is the production rate of enzyme A, $P_{prod,B'}$ is the production rate of the reactant $B'$, $k_{cat,A}$ is the maximal reaction rate of enzyme A, and $K_M$ is the half-saturation constant of enzyme A.

**Part 5: Irreversible binding motif.** In the irreversible binding motif, $A$ is the diffusible ligand, $B'$ is the unbound transcription factor, and $B$ is the bound transcription factor complex. This motif takes inspiration from irreversible ligand-binding transcription factors, such as the LasR family from *Pseudomonas aeruginosa* (76).

$$prod_A = P_{prod,A}, \tag{11}$$
$$prod_{B'} = P_{prod,B'}, \tag{12}$$
$$prod_B = k_{bind}AB', \tag{13}$$
$$R_{total} = prod_A + prod_{B'} + prod_B, \tag{14}$$

where $P_{prod,A}$ is the production rate of the diffusible ligand, $P_{prod,B'}$ is the production rate of the unbound transcription factor, and $k_{bind}$ is the rate constant for forming the bound transcription factor complex $B$.

**Part 6: Reversible binding motif.** In the reversible binding motif, $A$ is the diffusible ligand, $B'$ is the unbound transcription factor, and $B$ is the bound transcription factor complex.

$$prod_A = P_{prod,A}, \tag{15}$$
$$prod_{B'} = P_{prod,B'}, \tag{16}$$
$$prod_B = k_{bind}AB', \tag{17}$$
$$deg_B = k_{unbind}B, \tag{18}$$
$$R_{total} = prod_A + prod_{B'} + prod_B + deg_B, \tag{19}$$

where $P_{prod,A}$ is the production rate of the diffusible ligand, $P_{prod,B}$ is the production rate of the unbound transcription factor, $k_{bind}$ is the reaction constant for forming the bound transcription factor complex $B$, and $k_{unbind}$ is the unbinding rate constant for the bound complex $B$.

**Part 7: Phosphorylation motif.** In the phosphorylation motif (Figure S10), $A'$ is the unphosphorylated sensor histidine kinase, $A$ is the phosphorylated sensor histidine kinase, $B'$ is the unphosphorylated response regulator, $AB'$ is the bound complex of phosphorylated sensor histidine kinase and unphosphorylated response regulator, and $B$ is the phosphorylated response regulator.

$$prod_{A'} = P_{prod,A'}, \tag{20}$$
$$prod_A = k_aA', \tag{21}$$
$$prod_{B'} = P_{prod,B'}, \tag{22}$$
$$prod_{A'B} = k_bAB', \tag{23}$$
$$prod_B = k_t(AB'), \tag{24}$$
$$R_{total} = prod_{A'} + prod_A + prod_{B'} + prod_{A'B} + prod_B, \tag{25}$$

where $P_{prod,A'}$ is the production rate of the sensor histidine kinase, $k_a$ is the autophosphorylation rate of the sensor histidine kinase, $P_{prod,B'}$ is the production rate of the response regulator, $k_b$ is the binding rate constant of the phosphorylated sensor histidine kinase and unphosphorylated response regulator, and $k_t$ is the reaction rate for the transphosphorylation of the response regulator by the sensor histidine kinase.

By analyzing the production rate of $B$, we derived its production as a function of total A and unphosphorylated B as follows. The production rate of phosphorylated response regulator can be described as follows:

$$\frac{dB}{dt} = k_t(AB'), \tag{26}$$

where $k_t$ is the transphosphorylation rate constant and $(AB')$ is the concentration of bound $AB'$ complexes. We thus need to solve for $B$ as a function of total A and total B. To do so, we will apply a strategy similar to the derivation of the enzyme production equation (77). To start, we examine the rate equation for the $AB'$ complex:

$$\frac{dAB'}{dt} = k_b AB' - k_t(AB'). \tag{27}$$

We define $A_T$ as the total amount of A in the system (unphosphorylated, phosphorylated, and bound in the $AB'$ complex) and $B_T$ as the amount of unphosphorylated B.

$$A_T \equiv A' + A + (AB'), \tag{28}$$
$$B_T \equiv B' + (AB'). \tag{29}$$

We can then solve these expressions for $A$ and $B'$, respectively:

$$A = A_T - A' - (AB'), \tag{30}$$
$$B' = B_T - (AB'), \tag{31}$$

and substitute them into Equation 27:

$$\frac{dB}{dt} = k_b(A_T - A' - (AB'))(B_T - (AB')) - k_t(AB'). \tag{32}$$

We can then set this equation to the steady state:

$$0 = k_b(A_T - A' - (AB'))(B_T - (AB')) - k_t(AB'). \tag{33}$$

Finally, we must obtain A in terms of $A_T$, $B_T$, and $B$. To do so, we start with its rate equation:

$$\frac{dA'}{dt} = -k_a A' + k_t(AB') \tag{34}$$

and solve for its value at the steady state:

$$0 = -k_a A' + k_t(AB'), \tag{35}$$
$$k_a A' = k_t(AB'), \tag{36}$$
$$A' = \frac{k_t}{k_a}(AB'). \tag{37}$$

We can then substitute this expression into our steady-state equation for C:

$$0 = k_b(A_T - \frac{k_t}{k_a}(AB') - (AB'))(B_T - (AB')) - k_t(AB'), \tag{38}$$

which simplifies as follows:

$$0 = k_b(A_T - (1 + \frac{k_t}{k_a})(AB'))(B_T - (AB')) - k_t(AB'). \tag{39}$$

For clarity, we define $d \equiv 1 + \frac{k_t}{k_a}$. By substituting $d$ and expanding the expression, we obtain the following:

$$0 = k_b(A_T - d(AB'))(B_T - (AB')) - k_t(AB') \tag{40}$$
$$0 = k_b(A_T B_T - A_T(AB') - dB_T(AB') + d(AB')^2) - k_t(AB') \tag{41}$$
$$0 = k_b A_T B_T - k_b A_T(AB') - k_b dB_T(AB') + k_b d(AB')^2 - k_t(AB') \tag{42}$$
$$0 = k_b d(AB')^2 - (k_b A_T + k_b dB_T + k_t)(AB') + k_b A_T B_T. \tag{43}$$

Applying the quadratic formula gives us a solution for $(AB')(A_T, B_T)$:

$$(AB')(A_T, B_T) = \frac{(k_b A_T + k_b dB_T + k_t) - \sqrt{(k_b A_T + k_b dB_T + k_t)^2 - 4k_b^2 dA_T B_T}}{2k_b d}, \tag{44}$$

which we can then substitute into Equation 26:

$$\frac{dB}{dt} = \frac{k_t}{2k_b d}\left(k_b A_T + k_b dB_T + k_t - \sqrt{(k_b A_T + k_b dB_T + k_t)^2 - 4k_b^2 dA_T B_T}\right). \tag{45}$$

This expression describes the production rate of $B$ in terms of total A ($A_T$) and unphosphorylated B ($B_T$).

**Part 8: Bifunctional two-component system.** The bifunctional two-component system (Figure S11) is composed of an unphosphorylated sensor histidine kinase ($A'$) that reversibly autophosphorylates (producing $A$) and, in turn, reversibly binds to an unphosphorylated response regulator ($B'$), producing a bound complex ($AB'$). The phosphate group is then irreversibly transferred, restoring the unphosphorylated sensor histidine kinase ($A'$) and a phosphorylated response regulator ($B$). Distinguishing this motif from its monofunctional counterpart (SI Appendix C.7), the unphosphorylated sensor histidine kinase can then reversibly bind to the phosphorylated response regulator, creating a new complex ($A'B$), before acting as a phosphatase and irreversibly dephosphorylating the response regulator, restoring the unphosphorylated response regulator ($B'$). The events in the bifunctional two-component system are as follows:

$$prod_{A'} = P_{prod,A'}, \tag{46}$$
$$prod_{B'} = P_{prod,B'}, \tag{47}$$
$$phos_{A'} = k_k A', \tag{48}$$
$$dephos_A = k_{rev,k} A, \tag{49}$$
$$AB'_{bind} = k_1 AB', \tag{50}$$
$$AB'_{unbind} = k_{rev,1}(AB'), \tag{51}$$
$$phos_{B'} = k_t(AB'), \tag{52}$$
$$A'B_{bind} = k_2 A'B, \tag{53}$$
$$A'B_{unbind} = k_{rev,2}(A'B), \tag{54}$$
$$dephos_B = k_p(A'B), \tag{55}$$

where $P_{prod,A'}$ is the production probability of the unphosphorylated sensor histidine kinase, $P_{prod,B'}$ is the production probability of the unphosphorylated response regulator, $k_k$ is the sensor histidine kinase autophosphorylation rate constant, $A'$ is the amount of unphosphorylated sensor histidine kinase, $k_{rev,k}$ is the sensor histidine kinase autodephosphorylation rate constant, $A$ is the amount of phosphorylated sensor histidine kinase, $k_1$ is the binding rate constant of phosphorylated sensor histidine kinase and unphosphorylated response regulator, $B'$ is the amount of unphosphorylated response regulator, $k_{rev,1}$ is the unbinding rate constant of the bound complex of phosphorylated sensor histidine kinase and unphosphorylated response regulator, $AB'$ is the amount of the bound complex of phosphorylated sensor histidine kinase and unphosphorylated response regulator, $k_t$ is the transphosphorylation rate constant, $k_2$ is the binding rate constant of unphosphorylated sensor histidine kinase and phosphorylated response regulator, $B$ is the amount of phosphorylated response regulator, $k_{rev,2}$ is the unbinding constant of the complex of unphosphorylated sensor histidine kinase and phosphorylated response regulator, $A'B$ is the amount of the bound complex of unphosphorylated sensor histidine kinase and phosphorylated response regulator, and $k_p$ is the sensor histidine kinase phosphatase rate constant.

**Part 9: Production and degradation motif.** In the production and degradation motif (shown in Figure S9), A is the producing enzyme, B is the signal molecule, and C is the degrading enzyme:

$$prod_A = P_{prod,A}, \tag{56}$$
$$prod_C = P_{prod,C}, \tag{57}$$
$$prod_B = k_{cat,A}A, \tag{58}$$
$$deg_B = \frac{k_{cat,C}}{2}\left(K_M + B + C - \sqrt{(K_M + B + C)^2 - 4BC}\right), \tag{59}$$
$$R_{total} = prod_A + prod_C + prod_B + deg_B, \tag{60}$$

where $P_{prod,A}$ is the production rate of the producing enzyme A, $P_{prod,C}$ is the production rate of the degrading enzyme C, $k_{cat,A}$ is the reaction rate of the enzyme A, $k_{cat,C}$ is the maximal reaction rate of the enzyme C, and $K_M$ is the half-saturation constant of the enzyme C.

**Part 10: Diffusible ligand transcription factor pathway.** The diffusible ligand transcription factor pathway (Figure 5A) is composed of a binding motif followed by a saturated production motif, where $B'$ is the unbound transcription factor, A is the ligand, B is the bound transcription factor, and C is the protein product:

$$prod_A = P_{prod,A}, \tag{61}$$
$$prod_{B'} = P_{prod,B'}, \tag{62}$$
$$prod_B = k_{bind}AB', \tag{63}$$
$$prod_C = k_{prod}C, \tag{64}$$
$$R_{total} = prod_A + prod_{B'} + prod_B + prod_C, \tag{65}$$

where $P_{prod,A}$ is the diffusion rate of the ligand, $P_{prod,B'}$ is the production rate of the unbound transcription factor $B'$, $k_{bind}$ is the reaction constant for forming the bound transcription factor B, and $k_{prod}$ is the transcription rate for producing C.

**Part 11: Second messenger pathway.** The second messenger pathway (Figure 5C) is composed of a production and degradation motif followed by a saturated production motif, where A is the producing enzyme, C is the degrading enzyme, B is the signal molecule, and D is the protein product:

$$prod_A = P_{prod,A}, \tag{66}$$
$$prod_C = P_{prod,C}, \tag{67}$$
$$prod_B = k_{cat,A}A, \tag{68}$$
$$deg_B = \frac{k_{cat,B}}{2}\left(K_M + B + C - \sqrt{(K_M + B + C)^2 - 4BC}\right), \tag{69}$$
$$prod_C = k_{prod}B, \tag{70}$$
$$R_{total} = prod_A + prod_C + prod_B + deg_B + prod_C, \tag{71}$$

where $P_{prod,A}$ is the production rate of the producing enzyme A, $P_{prod,C}$ is the production rate of the degrading enzyme C, $k_{cat,A}$ is the reaction rate of the enzyme A, $k_{cat,B}$ is the maximal reaction rate of the enzyme B, $K_M$ is the half-saturation constant of the enzyme B, and $k_{prod}$ is the transcription rate for producing D.

**Part 12: Two-component system pathway.** The two-component system pathway (Figure 5E) is composed of a phosphorylation motif followed by a saturated production motif, where $A'$ is the unphosphorylated sensor histidine kinase, $A$ is the phosphorylated sensor histidine kinase, $B'$ is the unphosphorylated response regulator, $B$ is the phosphorylated response regulator, and $C$ is the protein product:

$$prod_{A'} = P_{prod,A'}, \tag{72}$$
$$prod_A = k_a A', \tag{73}$$
$$prod_{B'} = P_{prod,B'}, \tag{74}$$
$$prod_{AB'} = k_{bind} AB', \tag{75}$$
$$prod_B = k_t(AB'), \tag{76}$$
$$prod_C = k_{prod} B, \tag{77}$$
$$R_{total} = prod_{A'} + prod_A + prod_{B'} + prod_{AB'} + prod_B + prod_C, \tag{78}$$

where $P_{prod,A'}$ is the production rate of the sensor histidine kinase, $k_a$ is the autophosphorylation rate of the sensor histidine kinase, $P_{prod,B'}$ is the production rate of the response regulator, $k_{bind}$ is the binding constant for forming the complex of phosphorylated sensor histidine kinase and unphosphorylated response regulator, $k_t$ is the reaction rate for the transphosphorylation of the response regulator by the sensor histidine kinase, and $k_{prod}$ is the transcription rate for producing C.

**Part 13: Unsaturable phosphorylation monocycle.** The unsaturable phosphorylation monocycle (Figure S20B) is composed of a reactant molecule ($B'$) that is converted into a product molecule ($B$) by one enzyme ($A$, representing a kinase) and converted back by a second enzyme ($C$, representing a phosphatase), where each conversion is described by mass-action kinetics. The events in the pathway are as follows:

$$prod_A = P_{prod,A}, \tag{79}$$
$$prod_{B'} = P_{prod,B'}, \tag{80}$$
$$prod_C = P_{prod,C}, \tag{81}$$
$$prod_B = k_A B' A, \tag{82}$$
$$deg_B = k_C BC, \tag{83}$$

where $P_{prod,A}$ is the production probability of enzyme A, $P_{prod,B'}$ is the production probability of the substrate $B'$, $P_{prod,C}$ is the production probability of the enzyme C, $k_A$ is the binding constant for $A$ and $B'$, and $k_C$ is the binding constant for $C$ and $B$.

**Part 14: Saturable phosphorylation monocycle.** The saturable phosphorylation monocycle (Figure S20C) is composed of a reactant molecule ($B'$) that is converted into a product molecule ($B$) by one enzyme ($A$, representing a kinase) and converted back by a second enzyme ($C$, representing a phosphatase), where each conversion is described by enzyme kinetics. The events in the pathway are as follows:

$$prod_A = P_{prod,A}, \tag{84}$$
$$prod_{B'} = P_{prod,B'}, \tag{85}$$
$$prod_C = P_{prod,C}, \tag{86}$$
$$form_B = \frac{k_{cat,A}}{2}\left(K_{M,A} + A + B' - \sqrt{(K_{M,A} + A + B')^2 - 4AB'}\right), \tag{87}$$
$$form_{B'} = \frac{k_{cat,C}}{2}\left(K_{M,C} + C + B - \sqrt{(K_{M,C} + C + B)^2 - 4CB}\right), \tag{88}$$
$$R_{total} = prod_A + prod_B + prod_C + form_B + form_{B'}, \tag{89}$$

where $P_{prod,A}$ is the production probability of enzyme A, $P_{prod,B'}$ is the production probability of reactant B, $P_{prod,C}$ is the production probability of enzyme C, $k_{cat,A}$ is the maximal reaction rate of enzyme A, $K_{M,A}$ is the half-saturation constant of enzyme A, $k_{cat,C}$ is the maximal reaction rate of enzyme C, and $K_{M,C}$ is the half-saturation constant of enzyme C.

**Part 15: Five-layer cascade.** The five-layer cascade (Figure S12A) is composed of an initial enzyme A that catalyzes the production of molecule B, which in turn catalyzes the production of molecule C, which in turn catalyzes the production of molecule D, which in turn catalyzes the production of molecule E. The events in the pathway are as follows:

$$prod_A = P_{prod,A}, \quad (90)$$
$$prod_B = k_{cat,A}A, \quad (91)$$
$$prod_C = k_{cat,B}B, \quad (92)$$
$$prod_D = k_{cat,C}C, \quad (93)$$
$$prod_E = k_{cat,D}D, \quad (94)$$

where $P_{prod,A}$ is the production probability of molecule A, $k_{cat,A}$ is the maximal reaction rate of molecule A, $k_{cat,B}$ is the maximal reaction rate of molecule B, $k_{cat,C}$ is the maximal reaction rate of molecule C, and $k_{cat,D}$ is the maximal reaction rate of molecule D.

**Part 16: Gillespie algorithm and cell growth.** Once the sum of the event probabilities has been calculated, the event is selected using a random number spanning a uniform distribution between 0 and $R_{total}$. The length of the time step between events is then given by Equation 95:

$$\tau = (1/R_{total}) * \ln 1/r_1, \quad (95)$$

where $r_1$ is a random number pulled from a uniform random distribution spanning 0 to 1. After the time duration has been selected, the new cell volume is calculated, assuming linear growth:

$$V_{new} = V_{old} + \frac{\tau}{T_{cc}}, \quad (96)$$

where $T_{cc}$ is the cell cycle time. This process is repeated until $V > 2$, upon which the cell is divided. In the binomial partition method, different binomial random variables are pulled to partition each molecular agent. In the asymmetric partition model, a preset percentage is used to divide all molecules. The process then repeats for an arbitrary number of cell generations. The use of the exact Gillespie algorithm, while computationally expensive, ensures that our noise measurements are accurate for a sufficient sample size (78) and avoids linear approximations of noise that may otherwise underestimate the amount of variance in biologically realistic parameter ranges with few molecules (79).

#### SI Appendix D: Proof of variance equations

**Part 1: Effect of binomial partitioning on variance.** Let $X$ denote some molecular species (either enzyme A or product molecule B). The initial number of molecules, $X_0$, is a binomial partition of the number present in the mother before cell division, $X_M$. We thus have the following:

$$P(X_0|X_M) = Bin(X_M, 1/2). \tag{1}$$

By direct calculation,

$$\begin{aligned} \langle X_0^2 \rangle_{X_0} =& \langle X_0^2 \rangle_{X_0, X_M} && (2) \\ =& \langle \langle X_0^2 \rangle_{X_0|X_M} \rangle_{X_M} && (3) \\ =& \left\langle \frac{X_M}{4} + \left(\frac{X_M}{2}\right)^2 \right\rangle_{X_M} && (4) \\ =& \frac{1}{4}(\langle X_M \rangle_{X_M} + \langle X_M^2 \rangle_{X_M}). && (5) \end{aligned}$$

Moreover, we note that

$$\langle X_0 \rangle = \frac{1}{2}\langle X_M \rangle. \tag{6}$$

Thus,

$$\begin{aligned} Var(X_0) =& \langle X_0^2 \rangle_{X_0} - \langle X_0 \rangle_{X_0}^2 && (7) \\ =& \frac{1}{4}\left(\langle X_M^2 \rangle + \langle X_M \rangle\right) - \left(\frac{1}{2}\langle X_M \rangle\right)^2 && (8) \\ =& \frac{1}{4}(\langle X_M^2 \rangle - \langle X_M \rangle^2 + \langle X_M \rangle) && (9) \\ =& \frac{1}{4}(Var(X_M) + \langle X_M \rangle). && (10) \end{aligned}$$

**Part 2: Effect of Poisson signal accumulation.** During cell growth, new molecules are added at a stochastic rate. By definition,

$$\Delta X_t = X_t - X_0 \Longrightarrow X_t = X_0 + \Delta X_t, \tag{11}$$

where $X_t$ is the amount of molecule at time $t$ and $\Delta X_t$ is the amount added between time points $0$ and $t$. We have the following variance addition formula:

$$Var(X_t) = Var(X_0) + Var(\Delta X_t) + 2Cov(X_0, \Delta X_t). \tag{12}$$

Note that at steady state, we expect $X_M$ and $X_T$ to have the same statistics, where $X_T$ denotes the concentration at time $T_{cc}$, as the molecular distribution at the time of cell division eventually becomes constant across generations. In particular,

$$\langle X_T \rangle = \langle X_M \rangle \tag{13}$$

and

$$Var(X_T) = Var(X_M). \tag{14}$$

**Part 3: Statistics for the variance of enzyme A.** For enzyme A, we have

$$Cov(A_0, \Delta A_T) = 0. \tag{15}$$

Moreover, $\Delta A_T$ is generated by a Poisson process with rate $P_{prod,A}$. Therefore,

$$Var(\Delta A_T) = \langle \Delta A_T \rangle = P_{prod,A} T_{cc}. \tag{16}$$

This implies (from Part 2) that

$$Var(A_T) = Var(A_0) + P_{prod,A} T_{cc}. \tag{17}$$

Similarly,

$$\langle A_T \rangle = \langle A_0 \rangle + \langle \Delta A_T \rangle \quad (18)$$
$$= \langle A_0 \rangle + P_{prod,A} T_{cc}, \quad (19)$$
$$(20)$$

which, based on Part 1, is as follows:

$$\langle A_T \rangle = \frac{1}{2} \langle A_T \rangle + P_{prod,A} T_{cc} \quad (21)$$
$$= 2 P_{prod,A} T_{cc} \quad (22)$$

with

$$\langle A_0 \rangle = P_{prod,A} T_{cc}. \quad (23)$$

From Part 1, we also have

$$Var(A_0) = \frac{1}{4}(Var(A_T) + 2P_{prod,A} T_{cc}) \quad (24)$$
$$= \frac{1}{4}(Var(A_0) + P_{prod,A} T_{cc} + 2P_{prod,A} T_{cc}) \quad (25)$$
$$3Var(A_0) = 3P_{prod,A} T_{cc} \quad (26)$$
$$Var(A_0) = P_{prod,A} T_{cc} \quad (27)$$

and

$$Var(A_T) = 2P_{prod,A} T_{cc}. \quad (28)$$

From these expressions, we can calculate the variance of differences between our sister and random pairs. For the sister pairs,

$$\Delta A_{sis} = A_{sis,1} - A_{sis,2} = A_0 - (A_T - A_0) = 2A_0 - A_T. \quad (29)$$

Given $\langle \Delta A_{sis} \rangle = 0$, we obtain the variance as

$$Var(\Delta A_{sis}) = \langle \Delta A_{sis}^2 \rangle = 4\langle A_0^2 \rangle - 4\langle A_0 A_T \rangle + \langle A_T^2 \rangle. \quad (30)$$

We first consider that

$$\langle A_0 A_T \rangle = \langle \langle A_0 \rangle_{A_0|A_T} A_T \rangle_{A_T} = \frac{1}{2} \langle A_T^2 \rangle_{A_T}. \quad (31)$$

Therefore,

$$Var(\Delta A_{sis}) = 4\langle A_0^2 \rangle - \langle A_T^2 \rangle. \quad (32)$$

By substituting this expression in our derived formulas, we find

$$Var(\Delta A_{sis}) = ((P_{prod,A} T_{cc})^2 + P_{prod,A} T_{cc}) - ((2P_{prod,A} T_{cc})^2 + 2P_{prod,A} T_{cc}) \quad (33)$$
$$= 2P_{prod,A} T_{cc}. \quad (34)$$

It follows that the variance of the two random cells is two times the variance of each individual cell:

$$Var(\Delta A_{rnd}) = 2Var(A_0), \quad (35)$$
$$Var(\Delta A_{rnd}) = 2P_{prod,A} T_{cc}. \quad (36)$$

Note that the variance of difference formulas for sister and random pairs are equivalent when considering the enzyme A.

**Part 4: Persistence of Poisson statistics.** Suppose that

$$P(X_M) = Poiss(\bar{X}_M) \tag{37}$$

and

$$P(X_0|X_M) = Bin(X_M, 1/2). \tag{38}$$

Then, $X_0$ ~Poiss$(\bar{X}_M/2)$. We proceed by direct calculation:

$$P(X_0) = \sum_{X_M} P(X_0, X_M) \tag{39}$$
$$= \sum_{X_M} P(X_M)P(X_0|X_M) \tag{40}$$
$$= \sum_{X_M=X_0}^{\infty} \frac{(\bar{X}_M)^{X_M} e^{-\bar{X}_M}}{X_M!} \frac{X_M!(1/2)^{X_M}}{(X_M - X_0)!X_0!} \tag{41}$$
$$= \frac{e^{-\bar{X}_M}}{X_0!} \sum_{X_M=X_0}^{\infty} \frac{(\bar{X}_M/2)^{X_M}}{(X_M - X_0)!} \tag{42}$$
$$= \frac{e^{-\bar{X}_M}}{X_0!} \sum_{k=0}^{\infty} \frac{(\bar{X}_M/2)^{X_0+k}}{k!} \tag{43}$$
$$= \frac{e^{-\bar{X}_M}}{X_0!} \left(\frac{\bar{X}_M}{2}\right)^{X_0} e^{\bar{X}_M/2} \tag{44}$$
$$= \frac{(\bar{X}_M/2)^{X_0} e^{-\bar{X_M}/2}}{X_0!} \tag{45}$$
$$= Poiss(\bar{X}_M/2). \tag{46}$$

Thus, a Poisson random variable is still Poisson after it is binomially partitioned. As a corollary,

$$P(A_t) = Poiss(P_{prod,A}T_{cc} + P_{prod,A}t) \tag{47}$$

for all time points.

**Part 5: Calculating the mean of the product molecule B.** We write the following:

$$B_t = B_0 + \Delta B_t, \tag{48}$$

where $\Delta B_t$ is generated from a doubly-stochastic Poisson process with rate $k_{cat,A}A_t$. Thus,

$$\langle B_t \rangle = \langle B_0 \rangle + \langle \Delta B_t \rangle \tag{49}$$
$$= \langle B_0 \rangle + k_{cat,A} \int_0^t dt' A_{t'} \tag{50}$$
$$= \langle B_0 \rangle + k_{cat,A} \int_0^t dt' (P_{prod,A}T_{cc} + P_{prod,A}t') \tag{51}$$
$$= \langle B_0 \rangle + k_{cat,A}P_{prod,A}T_{cc}t + \frac{1}{2}k_{cat,A}P_{prod,A}t^2. \tag{52}$$

By setting $t = T_{cc}$ and noting that $\langle B_0 \rangle = \frac{1}{2}\langle B_T \rangle$, we obtain

$$\langle B_T \rangle = \left\langle \frac{B_T}{2} \right\rangle + \frac{3}{2} k_{cat,A} P_{prod,A} T_{cc}^2 \tag{53}$$

$$= 3 k_{cat,A} P_{prod,A} T_{cc}^2, \tag{54}$$

$$\langle B_0 \rangle = \frac{3}{2} k_{cat,A} P_{prod,A} T_{cc}^2, \tag{55}$$

$$\langle B_T \rangle = \frac{3}{2} k_{cat,A} P_{prod,A} T_{cc}^2 + k_{cat,A} P_{prod,A} T_{cc} t + \frac{1}{2} k_{cat,A} P_{prod,A} t^2. \tag{56}$$

**Part 6: Variance of accumulated product molecule B.** We next aim to calculate $Var(\Delta B_T)$. As a first step, we calculate

$$\langle \Delta B_T^2 \rangle = \langle \langle \Delta B_T^2 \rangle_{\Delta B_T | A_{1:T}} \rangle_{A_{1:T}}. \tag{57}$$

Because $\Delta B_T$ is Poisson when conditioned on $A_{1:T}$, we have

$$\langle \Delta B_T^2 \rangle = \left\langle k_{cat,A} \int_0^{T_{cc}} dt A_t + \left( k_{cat,A} \int_0^{T_{cc}} dt A_t \right)^2 \right\rangle_{A_{1:T}} \tag{58}$$

$$= k_{cat,A} \int_0^{T_{cc}} dt \langle A_t \rangle_{A_{1:T}} + k_{cat,A}^2 \int_0^{T_{cc}} dt \int_0^{T_{cc}} dt' \langle A_t A_{t'} \rangle_{A_{1:T}}. \tag{59}$$

By relabeling the integrated variables such that $t \leq t'$, we can write

$$A_{t'} = A_t + (\Delta A)_{(t,t')}, \tag{60}$$

where $(\Delta A)_{(t,t')}$ is the number of molecules of enzyme A added between $t$ and $t'$. Therefore,

$$\langle \Delta B_T^2 \rangle = k_{cat,A} \int_0^{T_{cc}} dt \langle A_t \rangle + k_{cat,A}^2 \int_0^{T_{cc}} dt \int_0^{t} dt' \langle A_t A_{t'} \rangle + k_{cat,A}^2 \int_0^{T_{cc}} dt \int_t^{T_{cc}} dt' \langle A_t A_{t'} \rangle \tag{61}$$

$$\begin{aligned} &= k_{cat,A} \int_0^{T_{cc}} dt \langle A_t \rangle + k_{cat,A}^2 \int_0^{T_{cc}} dt \int_0^{t} dt' \langle (A_{t'} + \Delta A_{(t',t)}) A_{t'} \rangle \\ &+ k_{cat,A}^2 \int_0^{T_{cc}} dt \int_t^{T_{cc}} dt' \langle A_t (A_t + \Delta A_{(t,t')}) \rangle \end{aligned} \tag{62}$$

$$\begin{aligned} &= k_{cat,A} \int_0^{T_{cc}} dt (P_{prod,A} T_{cc} + P_{prod,A} t) + k_{cat,A}^2 \int_0^{T_{cc}} dt \int_0^{t} dt' (\langle A_{t'}^2 \rangle + \langle \Delta A_{(t',t)} \rangle \langle A_{t'} \rangle) \\ &+ k_{cat,A}^2 \int_0^{T_{cc}} dt \int_t^{T_{cc}} dt' (\langle A_t^2 \rangle + \langle A_t \rangle \langle \Delta A_{(t,t')} \rangle). \end{aligned} \tag{63}$$

The first integral is

$$I_1 = k_{cat,A} \left( P_{prod,A} T_{cc}^2 + \frac{1}{2} P_{prod,A} T_{cc}^2 \right) \tag{64}$$

$$= \frac{3}{2} k_{cat,A} P_{prod,A} T_{cc}^2. \tag{65}$$

The second integral is

$$
\begin{aligned}
I_2 =& k_{cat,A}^2 \int_0^{T_{cc}} dt \int_0^t dt'(P_{prod,A}T_{cc} + P_{prod,A}t' + P_{prod,A}^2(T_{cc}+t')^2 + P_{prod,A}(t-t')P_{prod,A}(T_{cc}+t')) &\textbf{(66)}\\
=& k_{cat,A}^2 \int_0^{T_{cc}} dt \int_0^t dt'(P_{prod,A}T_{cc} + P_{prod,A}^2T_{cc}^2 + P_{prod,A}^2 tT_{cc} \\
&+ t'(P_{prod,A} + 2P_{prod,A}^2T_{cc} + P_{prod,A}^2 t - P_{prod,A}^2T_{cc}) + (t')^2(P_{prod,A}^2 - P_{prod,A}^2)) &\textbf{(67)}\\
=& k_{cat,A}^2 \int_0^{T_{cc}} dt(P_{prod,A}T_{cc}t + P_{prod,A}^2T_{cc}^2 t + P_{prod,A}^2T_{cc}t^2 + \frac{1}{2}P_{prod,A}t^2 \\
&+ P_{prod,A}^2T_{cc}t^2 + \frac{1}{2}P_{prod,A}^2t^3 - \frac{1}{2}P_{prod,A}^2T_{cc}t^2) &\textbf{(68)}\\
=& k_{cat,A}^2 \int_0^{T_{cc}} dt\left(t(P_{prod,A}T_{cc} + P_{prod,A}^2T_{cc}^2) + t^2\left(\frac{3}{2}P_{prod,A}^2T_{cc} + \frac{1}{2}P_{prod,A}\right) + \frac{1}{2}t^3P_{prod,A}^2\right) &\textbf{(69)}\\
=& k_{cat,A}^2\left(\frac{1}{2}P_{prod,A}T_{cc}^3 + \frac{1}{2}P_{prod,A}^2T_{cc}^4 + \frac{1}{2}P_{prod,A}^2T_{cc}^4 + \frac{1}{6}P_{prod,A}T_{cc}^3 + \frac{1}{8}P_{prod,A}^2T_{cc}^4\right) &\textbf{(70)}\\
=& k_{cat,A}^2\left(\frac{2}{3}P_{prod,A}T_{cc}^3 + \frac{9}{8}P_{prod,A}^2T_{cc}^4\right). &\textbf{(71)}
\end{aligned}
$$

The third integral is

$$
\begin{aligned}
I_3 =& k_{cat,A}^2 \int_0^{T_{cc}} dt \int_t^{T_{cc}} dt'(P_{prod,A}T_{cc} + P_{prod,A}t + P_{prod,A}^2(T_{cc}+t)^2 + P_{prod,A}(T_{cc}+t)P_{prod,A}(t'-t) &\textbf{(72)}\\
=& k_{cat,A}^2 \int_0^{T_{cc}} dt \int_t^{T_{cc}} dt'(P_{prod,A}(T_{cc}+t) + P_{prod,A}^2(T_{cc}+t)^2 - P_{prod,A}^2(T_{cc}+t)t + P_{prod,A}^2(T_{cc}+t)t') &\textbf{(73)}\\
=& k_{cat,A}^2 \int_0^{T_{cc}} dt((P_{prod,A}(T_{cc}+t)(T_{cc}-t) + P_{prod,A}^2(T_{cc}+t)T_{cc}(T_{cc}-t) \\
&+ \frac{1}{2}P_{prod,A}^2(T_{cc}+t)T_{cc}^2 - \frac{1}{2}P_{prod,A}^2(T_{cc}+t)t^2) &\textbf{(74)}\\
=& k_{cat,A}^2 \int_0^{T_{cc}} dt(P_{prod,A}T_{cc}^2 + \frac{3}{2}P_{prod,A}^2T_{cc}^3 + t(\frac{1}{2}P_{prod,A}^2T_{cc}^2) \\
&+ t^2(-P_{prod,A} - P_{prod,A}^2T_{cc} - \frac{1}{2}P_{prod,A}^2T_{cc}) + t^3(-\frac{1}{2}P_{prod,A}^2)) &\textbf{(75)}\\
=& k_{cat,A}^2\left(P_{prod,A}T_{cc}^3 + \frac{3}{2}P_{prod,A}^2T_{cc}^4 + \frac{1}{4}P_{prod,A}^2T_{cc}^4 - \frac{1}{3}P_{prod,A}T_{cc}^3 - \frac{1}{2}P_{prod,A}^2T_{cc}^4 - \frac{1}{8}P_{prod,A}^2T_{cc}^4\right) &\textbf{(76)}\\
=& k_{cat,A}^2\left(\frac{2}{3}P_{prod,A}T_{cc}^3 + \frac{9}{8}P_{prod,A}^2T_{cc}^4\right). &\textbf{(77)}
\end{aligned}
$$

Note that, as expected, $I_2 = I_3$. Putting these terms together, we obtain the following:

$$
\langle \Delta B_T^2 \rangle = \frac{3}{2}k_{cat,A}P_{prod,A}T_{cc}^2 + \frac{4}{3}k_{cat,A}^2P_{prod,A}T_{cc}^3 + \frac{9}{4}k_{cat,A}^2P_{prod,A}^2T_{cc}^4, \qquad \textbf{(78)}
$$

$$
\begin{aligned}
Var(\Delta B_T) =& \langle \Delta B_T^2 \rangle - \left(\frac{3}{2}k_{cat,A}P_{prod,A}T_{cc}^2\right)^2 &\textbf{(79)}\\
=& \frac{3}{2}k_{cat,A}P_{prod,A}T_{cc}^2 + \frac{4}{3}k_{cat,A}^2P_{prod,A}T_{cc}^3. &\textbf{(80)}
\end{aligned}
$$

**Part 7: Signal molecule covariance statistics.** The final quantity to compute is $Cov(B_0, \Delta B_T)$. This quantity is nonzero because both terms are correlated with the amount of enzyme in the mother cell. However, the terms become independent when conditioned on $A_0$:

$$P(B_0, \Delta B_T) = \sum_{A_0} P(B_0, \Delta B_T, A_0) \quad (81)$$

$$= \sum_{A_0} P(A_0) P(B_0, \Delta B_T | A_0) \quad (82)$$

$$= \sum_{A_0} P(A_0) P(B_0 | A_0) P(\Delta B_T | A_0). \quad (83)$$

This implies that

$$Cov(B_0, \Delta B_T) = \langle \langle \delta B_0 \rangle_{B_0|A_0} \langle \delta \Delta B_T \rangle_{\Delta B_T|A_0} \rangle_{A_0}, \quad (84)$$

where the $\delta$ notation indicates the difference of the random variable from its mean. We note that

$$P(\Delta B_T | A_t) = Poiss\left(k_{cat,A} \int_0^{T_{cc}} dt A_t\right) \quad (85)$$

$$= Poiss\left(k_{cat,A} \int_0^{T_{cc}} dt (A_0 + \Delta A_t)\right), \quad (86)$$

$$P(\Delta B_T | A_0) = \sum_{\Delta A_t} P(\Delta B_T, \Delta A_t | A_0) \quad (87)$$

$$= \sum_{\Delta A_t} P(\Delta A_t) P(\Delta B_T | A_0, \Delta A_t). \quad (88)$$

We then obtain

$$\langle \delta \Delta B_T \rangle_{\Delta B_T|A_0} = k_{cat,A} \int_0^{T_{cc}} dt \left( (A_0 + \langle \Delta A_t \rangle) - \frac{3}{2} k_{cat,A} P_{prod,A} T_{cc}^2 \right) \quad (89)$$

$$= k_{cat,A} A_0 T_{cc} + k_{cat,A} \int_0^{T_{cc}} dt \left( P_{prod,A} t - \frac{3}{2} k_{cat,A} P_{prod,A} T_{cc}^2 \right) \quad (90)$$

$$= k_{cat,A} A_0 T_{cc} + \frac{1}{2} k_{cat,A} P_{prod,A} T_{cc}^2 - \frac{3}{2} k_{cat,A} P_{prod,A} T_{cc}^2 \quad (91)$$

$$= k_{cat,A} A_0 T_{cc} - k_{cat,A} P_{prod,A} T_{cc}^2. \quad (92)$$

To compute $\langle B_0 \rangle_{B_0|A_0}$, we note the linear relationship between the $A$ and $B$ variables, which translates into a linear relationship between the means:

$$\langle B_0 \rangle_{B_0|A_0} = \frac{1}{2} \langle B_T \rangle_{B_T|A_0}, \quad (93)$$

$$\langle A_t \rangle_{A_t|A_0} = \langle A_T - \Delta A_{(t,T)} \rangle_{A_T, \Delta A_{(t,T)}|A_0} \quad (94)$$

$$= 2A_0 - P_{prod,A}(T_{cc} - t), \quad (95)$$

$$\langle B_T \rangle_{B_T|A_0} = k_{cat,A} \int_0^t dt' \langle A_{t'} \rangle_{A'_t|A_0} \tag{96}$$

$$= k_{cat,A} \int_0^t dt' (2A_0 - P_{prod,A} T_{cc} + P_{prod,A} t') \tag{97}$$

$$= 2k_{cat,A} A_0 t - k_{cat,A} P_{prod,A} T_{cc} t + \frac{1}{2} k_{cat,A} P_{prod,A} T_{cc}^2 \tag{98}$$

$$= 2k_{cat,A} A_0 T_{cc} - \frac{1}{2} k_{cat,A} P_{prod,A} T_{cc}^2. \tag{99}$$

Therefore,

$$\langle B_0 \rangle_{B_0|A_0} = k_{cat,A} A_0 T_{cc} - \frac{1}{4} k_{cat,A} P_{prod,A} T_{cc}^2 \tag{100}$$

and

$$\langle \delta B_0 \rangle_{B_0|A_0} = \langle B_0 \rangle_{B_0|A_0} - \frac{3}{2} k_{cat,A} P_{prod,A} T_{cc}^2 \tag{101}$$

$$= k_{cat,A} A_0 T_{cc} - \frac{7}{4} k_{cat,A} P_{prod,A} T_{cc}^2. \tag{102}$$

Combining these expressions yields the following:

$$Cov(B_0, \Delta B_T) = \left\langle \left( k_{cat,A} A_0 T_{cc} - \frac{7}{4} k_{cat,A} P_{prod,A} T_{cc}^2 \right) \left( k_{cat,A} A_0 T_{cc} - k_{cat,A} P_{prod,A} T_{cc}^2 \right) \right\rangle_{A_0} \tag{103}$$

$$= \frac{7}{4} k_{cat,A}^2 P_{prod,A}^2 T_{cc}^4 - k_{cat,A}^2 P_{prod,A} T_{cc}^3 \langle A_0 \rangle - \frac{7}{4} k_{cat,A}^2 P_{prod,A} T_{cc}^3 \langle A_0 \rangle + k_{cat,A}^2 T_{cc}^2 \langle A_0^2 \rangle \tag{104}$$

$$= k_{cat,A}^2 T_{cc}^2 (P_{prod,A} T_{cc} + P_{prod,A}^2 T_{cc}^2) - k_{cat,A}^2 P_{prod,A}^2 T_{cc}^4 \tag{105}$$

$$= k_{cat,A}^2 P_{prod,A} T_{cc}^3. \tag{106}$$

**Part 8: Putting it all together.** From Part 1,

$$Var(B_0) = \frac{1}{4} (Var(B_T) + \langle B_T \rangle) \tag{107}$$

$$= \frac{1}{4} (Var(B_T) + 3k_{cat,A} P_{prod,A} T_{cc}^2). \tag{108}$$

Substituting this into Equation 12 from Part 2,

$$Var(B_T) = \frac{1}{4} (Var(B_T) + 3k_{cat,A} P_{prod,A} T_{cc}^2) + \frac{3}{2} k_{cat,A} P_{prod,A} T_{cc}^2 + \frac{4}{3} k_{cat,A}^2 P_{prod,A} T_{cc}^3 + 2k_{cat,A}^2 P_{prod,A} T_{cc}^3 \tag{109}$$

$$\frac{3}{4} Var(B_T) = \frac{9}{4} k_{cat,A} P_{prod,A} T_{cc}^2 + \frac{10}{3} k_{cat,A}^2 P_{prod,A} T_{cc}^3 \tag{110}$$

$$Var(B_T) = 3k_{cat,A} P_{prod,A} T_{cc}^2 + \frac{40}{9} k_{cat,A}^2 P_{prod,A} T_{cc}^3. \tag{111}$$

This implies, by direct substitution, that

$$Var(B_0) = \frac{1}{4}(3k_{cat,A}P_{prod,A}T_{cc}^2 + \frac{40}{9}k_{cat,A}^2 P_{prod,A}T_{cc}^3 + 3k_{cat,A}P_{prod,A}T_{cc}^2) \quad (112)$$

$$= \frac{3}{2}k_{cat,A}P_{prod,A}T_{cc}^2 + \frac{10}{9}k_{cat,A}^2 P_{prod,A}T_{cc}^3. \quad (113)$$

Note that each variance expression looks like a Poisson variance plus an excess variance that can be attributed to the doubly-stochastic part.

**Part 9: Sister cell variance.** Finally, we calculate the variance of the difference between sister cells:

$$\Delta B_{sis} = B_{sis,1} - B_{sis,2} = B_0 - (B_T - B_0) = 2B_0 - B_T. \quad (114)$$

Clearly,

$$\langle \Delta B_{sis} \rangle = 0. \quad (115)$$

Therefore,

$$Var(\Delta B_{sis}) = \langle \Delta B_{sis}^2 \rangle = 4\langle B_0^2 \rangle - 4\langle B_0 B_T \rangle + \langle B_T^2 \rangle. \quad (116)$$

A component that we have not yet considered is

$$\langle B_0 B_T \rangle = \langle \langle B_0 \rangle_{B_0|B_T} B_T \rangle_{B_T} = \frac{1}{2}\langle B_T^2 \rangle_{B_T}. \quad (117)$$

Therefore,

$$Var(\Delta B_{sis}) = 4\langle B_0^2 \rangle - \langle B_T^2 \rangle. \quad (118)$$

By substituting Equations 114 and 115 from Part 8, we find

$$Var(\Delta B_{sis}) = 4\left(\left(\frac{3}{2}k_{cat,A}P_{prod,A}T_{cc}^2\right)^2 + \frac{3}{2}\left(k_{cat,A}P_{prod,A}T_{cc}^2 + \frac{10}{9}k_{cat,A}^2 P_{prod,A}T_{cc}^3\right)\right)$$
$$- \left(\left(\frac{3}{2}k_{cat,A}P_{prod,A}T_{cc}^2\right)^2 + \frac{3}{2}\left(k_{cat,A}P_{prod,A}T_{cc}^2 + \frac{10}{9}k_{cat,A}^2 P_{prod,A}T_{cc}^3\right)\right) \quad (119)$$

$$Var(\Delta B_{sis}) = 3k_{cat,A}P_{prod,A}T_{cc}^2. \quad (120)$$

**Part 10: Random pair variance.** It follows that the variance of two random cells would be two times the variance of each individual cell:

$$Var(\Delta B_{rnd}) = 2Var(B_0) \quad (121)$$

$$= 3k_{cat,A}P_{prod,A}T_{cc}^2 + \frac{20}{9}k_{cat,A}^2 P_{prod,A}T_{cc}^3. \quad (122)$$

#### SI Appendix E: Ultrasensitive reactions lead to ultrasensitivity in lineage-associated similarity

Having shown that reaction rate saturation is required for lineage-associated similarity, we next investigated how the saturation-related phenomenon of zero-order ultrasensitivity (80) affects the relationship between saturation and similarity among related cells in our framework. We evaluated two models of the canonical zero-order ultrasensitivity reaction motif. We first evaluated the phosphorylation monocycle (81), in which an unphosphorylated substrate ($B'$) is phosphorylated by a kinase ($A$) into a phosphorylated form ($B$) and dephosphorylated back to its unphosphorylated state by a phosphatase ($C$) (Figure S20A). Note that this motif differs from the monofunctional phosphorylation motif (SI Appendix C.7, Figure S10) and bifunctional phosphorylation motif (SI Appendix C.8, Figure S11) by having the enzyme A always be in an active state, whereas the phosphorylation motifs require autophosphorylation of enzyme A. Our evaluation of a phosphorylation monocycle modeled with mass-action kinetics (SI Appendix C.13), which is known to not exhibit zero-order ultrasensitivity (82), exhibited a graded response of lineage-associated similarity in the phosphorylated substrate ($B$) as the total amount of substrate increased (Figure S20B).

We next examined a saturable version of the phosphorylation monocycle (SI Appendix C.14). Here, we found that the lineage-associated similarity of the product amount with respect to the total substrate exhibited a steep transition when the amount of the enzyme matched the half-saturation constant (Figure S20C, red lines). When the enzyme was not saturated, the transitions were more shallow (Figure S20C, blue lines), indicating that saturation in a phosphorylation monocycle with enzyme kinetics leads to ultrasensitivity in lineage-associated similarity.

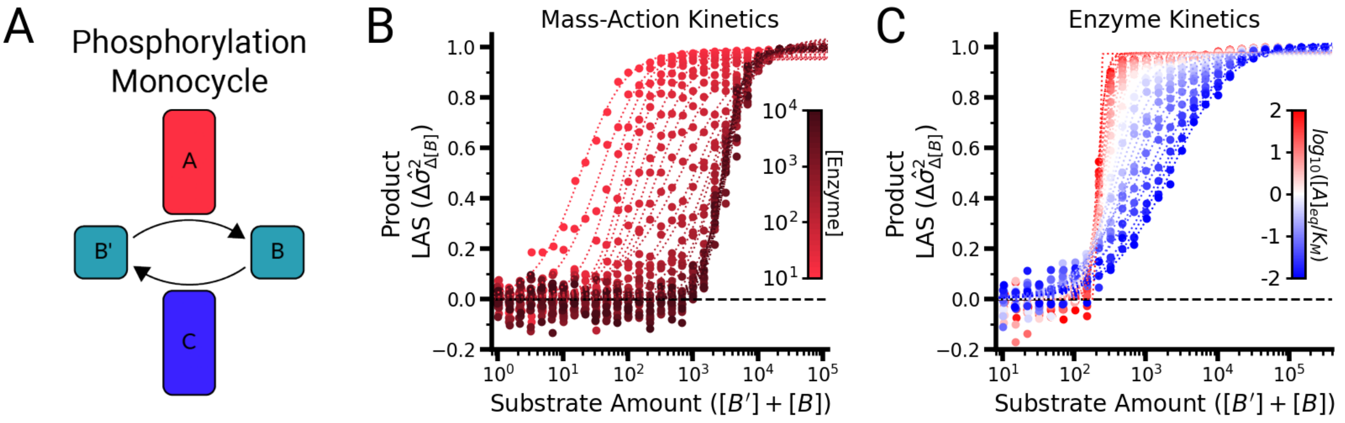


**Fig. S20. Saturation of the substrate and enzyme leads to ultrasensitive behavior of lineage-associated similarity | (A)** Schematic of the saturable phosphorylation monocycle motif, where substrate $B'$ is converted into product $B$ by enzyme $A$ and converted back by enzyme $C$. **(B)** Lineage-associated similarity of the product molecule ($B$) as a function of the total substrate amount ($[B'] + [B]$) for various concentrations of enzyme simulated with mass-action kinetics. **(C)** Lineage-associated similarity of the product molecule ($B$) as a function of the total substrate amount ($[B'] + [B]$) for various enzyme half-saturation constants ($K_{M,A}$ and $K_{M,C}$) simulated with enzyme kinetics. See SI Appendix G for all simulation parameter values.

#### SI Appendix F: Multiple mechanisms near and at saturation confer lineage-associated similarity in stochastic biochemical reactions

To understand how biologically realistic assumptions about saturation create similarity in both reactants and products, we focused on the relationship between the amount of reactant and the amount of enzyme. We chose this comparison because, while the saturating amount of reactant per enzyme scales nonlinearly with enzyme and reactant concentration, it still provides a guide for the frequency at which a reactant molecule would encounter an enzyme. Examining how similarity is affected by this ratio, we found that reactant similarity is present when the equilibrium reactant-to-enzyme ratio surpasses 1:1 (Figure 4C), corresponding to the start of the physiologically realistic regime (36). As shown previously for binomially partitioned intracellular molecules, lineage-associated similarity indicates that the reactant's concentration distribution has a wider variance than a Poisson distribution. A convenient metric for comparing a distribution's variance and a Poissonian distribution is the Fano factor, with Poisson processes exhibiting a Fano factor of 1. We observe maximal Fano factor values in the reactant's distribution just above a saturation ratio of 1 (defined as the mean concentration of reactant divided by the half-saturation constant amount of enzyme) (Figure S21B), corresponding to a peak in the similarity curve with respect to the substrate-to-enzyme ratio (Figure 4C). This increase in noise peaking at values just above the half-saturation constant for stochastically produced reactants has been observed elsewhere (83), reinforcing that our framework is consistent with other theoretical studies.

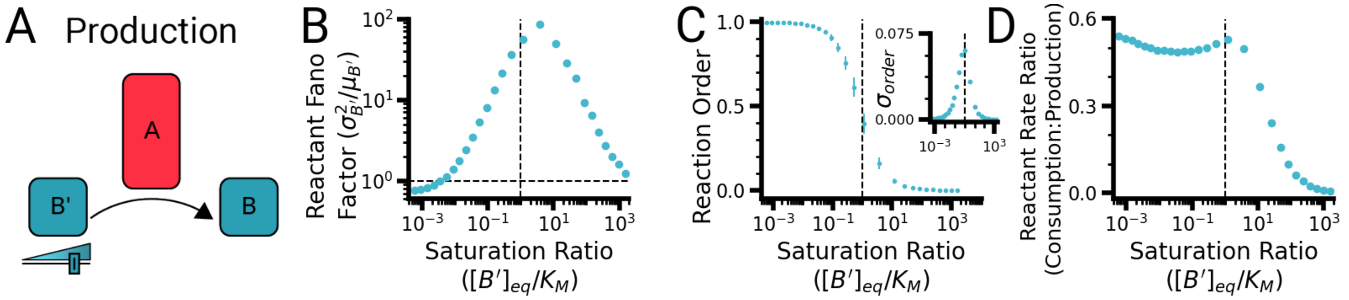


**Fig. S21. Reaction order variance drives lineage-associated similarity in the reactant molecule** | **(A)** Schematic depicting the production motif, in which a reactant molecule ($B'$) is converted into a product molecule ($B$) by an enzyme ($A$). **(B)** Reactant Fano factor as a function of the saturation ratio, defined as the reactant concentration divided by the half-saturation constant. The vertical dashed line indicates a saturation ratio of 1, and the horizontal dashed line indicates a Fano factor of 1, corresponding to a Poisson distribution. **(C)** Reaction order as a function of the saturation ratio. The vertical dashed line indicates a saturation ratio of 1, and the error bars indicate the standard deviation. **Inset:** Standard deviation of the reaction order as a function of the saturation ratio. **(D)** Reactant rate ratio, defined as the reaction rate divided by the production rate, as a function of saturation ratio. The vertical dashed line indicates a saturation ratio of 1. See SI Appendix G for all simulation parameter values.

We suggest that the increased variance in the reactant concentration distribution driving the Fano factor peak prior to saturation arises from reactant order switching (Figure S21B). Specifically, as the availability of reactant increases and the reaction order decreases from 1, the stochastic reaction begins to frequently switch its exact reaction order (Figure S21C), creating variable amounts of competition between the production and consumption of the reactant. This varying rate of competition increases the variance in the reactant's possible concentration distribution to be super-Poissonian (Figure S21B). This variance in reaction order switching peaks just after a saturation ratio of 1 is reached and then begins to decline (Figure S21C, inset). However, the reactant lineage-associated similarity does not decrease as quickly as the reaction order variance: it decreases more slowly in the approaching- and at-saturation regime because the competition between production and consumption of the reactant continues to generate a super-Poissonian variance in possible amounts of reactant (Figure S21D). As the production of the reactant increases to the point at which the saturated reaction cannot keep up, the reactant lineage-associated similarity goes to zero, as we are now approximating the fully saturated production circuit (explored in Figure 2B, C, and G, and 3A).

What controls the amount of substrate required for lineage-associated similarity in the product? We know that the amount of similarity in a product molecule is set by the amplification factor when the reactant is not limiting (Figure 2E) but that the amount of reactant can limit the extent of similarity in the product (Figure 4B). This suggests a critical role for reaction rate saturation. We find that across a wide range of enzyme concentrations, a shift from a lack of similarity in the product molecule to similarity occurs at a saturation ratio of 1 (Figure 4D). We explored the steepness of this transition through the lens of ultrasensitivity using a phosphorylation monocycle motif (SI Appendix E). Experimental measurements of reactant concentrations up to half-saturation constants in *E. coli* indicate that most metabolic reactions occur at ratios of 1:1–10:1 (35, 36), (Figure 4D, shaded area), suggesting that natural systems operate in ranges where similarity in the product exists.

These findings suggest that a key requirement of a reaction to produce lineage-associated similarity is the capacity for the reaction to be saturated. We validated this finding by examining a reaction motif that does not saturate: a simple binding motif, where two molecules associate together to produce a bound product (SI Appendices B.5 and B.6 for irreversible and reversible binding motif analyses), and found no similarity in any component of the motif (Figure S7). To determine if this requirement of saturation for lineage-associated similarity was present in other reaction motifs, we also examined a phosphorylation cascade (SI Appendix C.7), inspired by bacterial two-component systems (84, 85). In a monofunctional sensor histidine kinase model, as is found in the membrane fluidity-sensing DesK:DesR complex in *Bacillus subtilis* (86), the production rate of the phosphorylated response regulator (akin to the product molecule B) saturates as a function of the amount of sensor histidine kinase (similar to enzyme A, see SI Appendix C.7), and lineage-associated similarity was similarly present when the motif was stoichiometrically saturated (Figure S10). In a more complex bifunctional sensor histidine kinase circuit (SI Appendix C.8) (87), similarity was also found at reactant saturation for low levels of sensor histidine kinase (Figure S11). Notably, natural ratios of sensor histidine kinases and response regulators suggest that most of these systems operate in a saturated regime (88–90), where lineage-associated similarity would exist.

#### SI Appendix G: Simulation parameter values

| Figure Panel | Motif | Parameter Values |
|---|---|---|
| Figure 2B & C | Saturated Production | $n_{cells} = 1000$<br>$T_{cc} = 1000$<br>$P_{prod,A} = 10^{-3} - 10^{2}$<br>$k_{cat,A} = 10^{-1}$ |
| Figure 2F | Single with Bias | $n_{cells} = 1000$<br>$T_{cc} = 1000$<br>$P_{prod,A} = 10^{-1}$<br>$biases = 0.5 - 1$ |
| Figure 2G | Saturated Production | $n_{cells} = 1000$<br>$T_{cc} = 1000$<br>$P_{prod,A} = 10^{-1}$<br>$k_{cat,A} = 10^{-1}$ |
| Figure 2H | Single with Bursting | $n_{cells} = 1000$<br>$T_{cc} = 1000$<br>$P_{prod,A} = 10^{0}/burstSize$<br>$burstSize = 1 - 20$ |
| Figure 2I | Single with Varying Cell Cycle Time | $n_{cells} = 1000$<br>$T_{cc} = 1000$<br>$P_{prod,A} = 10^{-1}$<br>$\sigma_{T_{cc}} = 10^{-3} - 10^{3}$ |
| Figure 3A | Saturated Production | $Sweepk_{cat}$<br>$n_{cells} = 1000$<br>$T_{cc} = 1000$<br>$P_{prod,A} = 10^{-2}$<br>$k_{cat,A} = 10^{-4} - 10^{0}$<br><br>$SweepT_{cc}$<br>$n_{cells} = 1000$<br>$T_{cc} = 10^{2} - 10^{4}$<br>$P_{prod,A} = 10^{-2}$<br>$k_{cat,A} = 10^{-2}$<br><br>$Sweepk_{cat}\ 2$<br>$n_{cells} = 1000$<br>$T_{cc} = 1000$<br>$P_{prod,A} = 10^{-1}$<br>$k_{cat,A} = 10^{-3} - 10^{3}$<br><br>$SweepT_{cc}2$<br>$n_{cells} = 1000$<br>$T_{cc} = 500 - 10000$<br>$P_{prod,A} = 10^{-1}$<br>$k_{cat,A} = 10^{-1}$ |
| Figure 3C | Production and Degradation | $n_{cells} = 1000$<br>$T_{cc} = 1000$<br>$P_{prod,A} = 10^{-1}$<br>$P_{prod,C} = 10^{-1}$<br>$k_{cat,A} = 0.5$<br>$k_{cat,C} = 0.9$<br>$K_M = 5000$ |

| Figure Panel | Motif | Parameter Values |
| --- | --- | --- |
| Figure 3D | Production and Degradation | $n_{cells} = 1000$<br>$T_{cc} = 1000$<br>$P_{prod,A} = 10^{-1}$<br>$P_{prod,C} = 10^{-1}$<br>$k_{cat,A} = 0.5 - 1$<br>$k_{cat,C} = 0.9 - 1.9$<br>$K_M = 5000$ |
| Figure 4B | Unsaturated Production | $n_{cells} = 1000$<br>$T_{cc} = 1000$<br>$P_{prod,A} = 10^{-1}$<br>$k_{cat,A} = 10^{-1}$<br>$K_M = 10^3$<br>$P_{prod,B} = 10^{-2} - 10^4$ |
| Figure 4C & D | Unsaturated Production | $n_{cells} = 1000$<br>$T_{cc} = 1000$<br>$P_{prod,A} = 10^{-2} - 10^1$<br>$k_{cat,A} = 10^{-1}$<br>$K_M = 10^3$<br>$P_{prod,B} = 10^{-2} - 10^4$ |
| Figure 5B | Diffusible Transcription Factor | $n_{cells} = 1000$<br>$n_{cycles} = 10$<br>$T_{cc} = 1000$<br>$P_{prod,A} = 10^{-1}$<br>$P_{prod,B} = 10^{-1}$<br>$k_{bind} = 10^{-3}$<br>$k_{prod} = 10^{-1}$ |
| Figure 5D | cdG Circuit | $n_{cells} = 1000$<br>$n_{cycles} = 10$<br>$T_{cc} = 1000$<br>$P_{prod,A} = 10^{-1}$<br>$P_{prod,B} = 10^{-1}$<br>$k_{cat,A} = 10^{-1}$<br>$k_{cat,B} = 10^{-1}$<br>$K_M = 5000$<br>$k_{prod} = 10^{-1}$ |
| Figure 5F | Two-Component System | $n_{cells} = 1000$<br>$n_{cycles} = 10$<br>$T_{cc} = 1000$<br>$P_{prod,A} = 10^{-1}$<br>$P_{prod,B} = 10^1$<br>$k_a = 10^{-2}$<br>$k_b = 10^{-4}$<br>$k_t = 10^{-2}$<br>$k_4 = 10^{-2}$ |
| Figure 6B | Cascade | $max_{cells} = 2^{10}$<br>$P_{prod,A} = 10^{-1}$<br>$k_{cat,A} = 10^{-2}$<br>$k_{cat,B} = 10^{-2}$<br>$T_{cc} = 1000$<br>$\sigma_{T_{cc}} = 10$ |
| Figure 6C | Cascade | $n_{cells} = 1000$<br>$n_{cycles} = 10$<br>$T_{cc} = 1000$<br>$P_{prod,A} = 10^{-1}$<br>$k_{cat,A} = 10^{-1}$<br>$k_{cat,B} = 10^{-1}$ |

| Figure Panel | Motif | Parameter Values |
|---|---|---|
| Figure 6D | Cascade | $max_{cells} = 2^{10}$<br>$P_{prod,A} = 10^{-1}$<br>$k_{cat,A} = 10^{-2}$<br>$k_{cat,B} = 10^{-2}$<br>$T_{cc} = 1000$<br>$\sigma_{T_{cc}} = 10$ |
| Figure S2 | Saturated Production | $n_{cells} = 1000$<br>$T_{cc} = 1000$<br>$P_{prod,A} = 10^{-3} - 10^{2}$<br>$k_{cat,A} = 10^{-1}$ |
| Figure S3 | Saturated Production | $n_{cells} = 1000$<br>$T_{cc} = 1000$<br>$P_{prod,A} = 10^{-3} - 10^{2}$<br>$k_{cat,A} = 10^{-1}$ |
| Figure S4 | Saturated Production | $Sweepk_{cat}$<br>$n_{cells} = 1000$<br>$T_{cc} = 1000$<br>$P_{prod,A} = 10^{-2}$<br>$k_{cat,A} = 10^{-4} - 10^{0}$<br><br>$SweepT_{cc}$<br>$n_{cells} = 1000$<br>$T_{cc} = 10^{2} - 10^{4}$<br>$P_{prod,A} = 10^{-2}$<br>$k_{cat,A} = 10^{-2}$<br><br>$Sweepk_{cat}$ 2<br>$n_{cells} = 1000$<br>$T_{cc} = 1000$<br>$P_{prod,A} = 10^{-1}$<br>$k_{cat,A} = 10^{-3} - 10^{3}$<br><br>$SweepT_{cc}2$<br>$n_{cells} = 1000$<br>$T_{cc} = 500 - 10000$<br>$P_{prod,A} = 10^{-1}$<br>$k_{cat,A} = 10^{-1}$ |
| Figure S5 | Saturated Production | $n_{cells} = 1000$<br>$T_{cc} = 10^{2} - 10^{4}$<br>$P_{prod,A} = 10^{-2}$<br>$k_{cat,A} = 10^{-4} - 10^{0}$ |
| Figure S6B | Fixed Reactant | $n_{cells} = 1000$<br>$T_{cc} = 1000$<br>$P_{prod,A} = 10^{-2} - 10^{0}$<br>$k_{cat,A} = 10^{-1}$<br>$K_M = 1000$<br>$P_{prod,B} = 10^{-3} - 10^{3}$ |
| Figure S6C | Fixed Reactant | $n_{cells} = 1000$<br>$T_{cc} = 1000$<br>$P_{prod,A} = 10^{-1}$<br>$k_{cat,A} = 10^{-2} - 10^{0}$<br>$K_M = 1000$<br>$P_{prod,B} = 10^{-3} - 10^{3}$ |

| Figure Panel | Motif | Parameter Values |
|---|---|---|
| Figure S7B-D | Binding | $n_{cells} = 2000$<br>$T_{cc} = 1000$<br>$P_{prod,A} = 10^-3 - 10^2$<br>$P_{prod,B} = 10^{-3} - 10^2$<br>$k_{bind} = 10^{-3}$ |
| Figure S7F-H | Reversible Binding | $n_{cells} = 2000$<br>$T_{cc} = 1000$<br>$P_{prod,A} = 10^{-3} - 10^2$<br>$P_{prod,B} = 10^{-3} - 10^2$<br>$k_1 = 10^{-3}$<br>$k_2 = 10^{-5}$ |
| Figure S8B & E | Saturated Production | $n_{cells} = 1000$<br>$n_{cycles} = 10$<br>$T_{cc} = 1000$<br>$P_{prod,A} = 10^{-3} - 10^0$<br>$k_{cat,A} = 10^{-1}$ |
| Figure S8C & F | Saturated Production | $n_{cells} = 1000$<br>$n_{cycles} = 10$<br>$T_{cc} = 1000$<br>$P_{prod,A} = 10^{-1}$<br>$k_{cat,A} = 10^{-3} - 10^0$ |
| Figure S8D & G | Saturated Production | $n_{cells} = 1000$<br>$n_{cycles} = 10$<br>$T_{cc} = 500 - 5000$<br>$P_{prod,A} = 10^{-1}$<br>$k_{cat,A} = 10^{-1}$ |
| Figure S9 | Production and Degradation | $n_{cells} = 1000$<br>$T_{cc} = 1000$<br>$P_{prod,A} = 10^{-1}$<br>$P_{prod,C} = 10^{-1}$<br>$k_{cat,A} = 0.5 - 1$<br>$k_{cat,C} = 0.9 - 1.9$<br>$K_M = 5000$ |
| Figure S10 | Phosphorylation Cascade | $n_{cells} = 2000$<br>$T_{cc} = 1000$<br>$P_{prod,A} = 10^-3 - 10^2$<br>$P_{prod,B} = 10^{-3} - 10^2$<br>$k_a = 10^{-2}$<br>$k_b = 10^{-4}$<br>$k_t = 10^{-2}$ |
| Figure S11 | Phosphorylation Cascade Full | $n_{cells} = 2000$<br>$T_{cc} = 1000$<br>$P_{prod,A} = 10^{-3} - 10^2$<br>$P_{prod,B} = 10^{-3} - 10^2$<br>$k_k = 10^{-2}$<br>$k_{rev,k} = 10^{-2}$<br>$k_1 = 10^{-4}$<br>$k_{rev,1} = 10^{-2}$<br>$k_2 = 10^{-4}$<br>$k_{rev,2} = 10^{-2}$<br>$k_t = 10^{-2}$<br>$k_p = 10^{-2}$ |

| Figure Panel | Motif | Parameter Values |
|---|---|---|
| Figure S12 | Cascade (5) | $n_{cells} = 1000$<br>$n_{cycles} = 20$<br>$T_{cc} = 1000$<br>$P_{prod,A} = 5 * 10^{-3}$<br>$k_{cat,A} = 5 * 10^{-3}$<br>$k_{cat,B} = 5 * 10^{-3}$<br>$k_{cat,C} = 5 * 10^{-3}$<br>$k_{cat,D} = 5 * 10^{-3}$ |
| Figure S14 | Cascade | $max_{cells} = 2^{10}$<br>$P_{prod,A} = 10^{-1}$<br>$k_{cat,A} = 10^{-2}$<br>$k_{cat,B} = 10^{-2}$<br>$T_{cc} = 1000$<br>$\sigma_{T_{cc}} = 10$ |
| Figure S15 | Cascade | $max_{cells} = 2^{10}$<br>$P_{prod,A} = 10^{-1}$<br>$k_{cat,A} = 10^{-2}$<br>$k_{cat,B} = 10^{-2}$<br>$T_{cc} = 1000$<br>$\sigma_{T_{cc}} = 10$ |
| Figure S16 | Cascade | $max_{cells} = 2^{10}$<br>$P_{prod,A} = 10^{-1}$<br>$k_{cat,A} = 10^{-2}$<br>$k_{cat,B} = 10^{-2}$<br>$T_{cc} = 1000$<br>$\sigma_{T_{cc}} = 10$ |
| Figure S18 & G | Saturated Production | $n_{cells} = 1000$<br>$n_{cycles} = 10$<br>$T_{cc} = 1000$<br>$P_{prod,A} = 10^{-1}$<br>$k_{cat,A} = 10^{-1}$ |
| Figure S20B | Phosphorylation Monocycle | $n_{cells} = 2000$<br>$T_{cc} = 1000$<br>$P_{prod,A} = 10^{-3} - 10^{2}$<br>$P_{prod,B} = 10^{-3} - 10^{2}$<br>$P_{prod,C} = 10^{-1}$<br>$k_a = 10^{-1}$<br>$k_c = 10^{-1}$ |
| Figure S20C | Saturated Phosphorylation | $n_{cells=2000}$<br>$T_{cc} = 1000$<br>$P_{prod,A} = 10^{-1}$<br>$k_a = 10^{-2}$<br>$K_{M,A} = 10^{0} - 10^{4}$<br>$P_{prod,B} = 10^{-2} - 10^{3}$<br>$P_{prod,C} = 10^{-1}$<br>$k_c = 10^{-2}$<br>$K_{M,C} = 10^{0} - 10^{4}$ |
| Figure S21 | Unsaturated Production | $n_{cells} = 1000$<br>$T_{cc} = 1000$<br>$P_{prod,A} = 10^{-1}$<br>$k_{cat,A} = 10^{-1}$<br>$P_{prod,B} = 10^{-2} - 10^{3}$<br>$K_M = 1000$ |

### SI Appendix H: Code and Data Table

| Figure | Code File | Raw Data Files | Analysis Code | Analyzed Data File |
|---|---|---|---|---|
| Fig 2B, C | satprod_prodAsweep.py | satprod_prodAsweep_[0-5].pickle | | |
| Fig 2F | single_partition_bias_screen.py | asymmetric_bias_screen.pickle | | |
| Fig 2G | satprod_prodAsweep.py | satprod_prodAsweep_2.pickle | | |
| Fig 2H | satprod_burst_size_sweep.py | burstSize_prodA-[0-2].pickle | | |
| Fig 2I | single_var_cell_cycle_time.py | varTcc_diffs.pickle | | |
| Fig 3A | prodsat_sweep_kcat.py<br>prodsat_sweep_Tcc.py<br>prodsat_sweep_kcat_2.py<br>prodsat_sweep_Tcc_2.py | motifs_prodsat_kcatsweep.pickle<br>motifs_prodsat_Tccsweep.pickle<br>prodsat_kcatsweep_kcatA_[00-11].pickle<br>prodsat_Tccsweep_Tcc_[00-04].pickle | | |
| Fig 3C | proddeg_rkcat_sweep.py | Rkcatsweep2_0.pickle | | |
| Fig 3D | proddeg_rkcat_sweep.py | Rkcatsweep2_0.pickle<br>Rkcatsweep2_5.pickle<br>Rkcatsweep2_9.pickle<br>Rkcatsweep2_18.pickle | | |
| Fig 4B | produnsat_sweep_PprodB.py | prodBsweep5_PprodB_[00-28].pickle | produnsat_sweep_PprodB_aggregate.py | production_prodBsweep5.pickle |
| Fig 4C, D | produnsat_sweep_PprodA_PprodB.py | prodAprodBsweep1_PprodA_[00-15]_PprodB_[00-30].pickle | produnsat_sweep_PprodA_PprodB_aggregate.py | production_prodAprodBsweep1.pickle |
| Fig 5B | diffTF_time.py | motifs_diffTF3.pickle | | |
| Fig 5D | cdg_time.py | motifs_cdg_time.pickle | | |
| Fig 5F | tcs_time.py | motifs_tcs3.pickle | | |
| Fig 6B | cascade_spatial.py | cascade_10gen2.pickle | cascade_spatial_analyze.py | cascade_10gen2_molConcMaps.pickle |
| Fig 6C | cascade_time.py | motifs_cascade.pickle | cascade_time_analyze.py | cascade_time_normdvar.pickle |
| Fig 6D | cascade_spatial_moranI.py | cascade_10gen2_morIs_discdist_r[0-9].pickle | cascade_spatial_analyze.py | cascade_10gen_moranIs2.pickle |
| Fig S2 | satprod_prodAsweep.py | satprod_prodAsweep_[0-5].pickle | | |
| Fig S3 | satprod_prodAsweep.py | satprod_prodAsweep_[0-5].pickle | | |
| Fig S4 | prodsat_sweep_kcat.py<br>prodsat_sweep_Tcc.py<br>prodsat_sweep_kcat_2.py<br>prodsat_sweep_Tcc_2.py | motifs_prodsat_kcatsweep.pickle<br>motifs_prodsat_Tccsweep.pickle<br>prodsat_kcatsweep_kcatA_[00-11].pickle<br>prodsat_Tccsweep_Tcc_[00-04].pickle | | |
| Fig S5 | prodsat_sweep_kcatA_Tcc.py | motifs_prodsat_2dsweep_prodA_0_kcatA_[0-8].pickle | | |
| Fig S6B | fixedreactant_sweep_PprodA.py | fixedreactant4_PprodA_[00-02]_PprodB_[00-30].pickle | fixedreactant_sweep_PprodA_aggregate.py | fixedReactant4.pickle |
| Fig S6C | fixedreactant_sweep_kcatA.py | fixedreactant5_kcatA_[00-02]_PprodB_[00-30].pickle | fixedreactant_sweep_kcatA_aggregate.py | fixedReactant5.pickle |
| Fig S7B-D | bind_sweep_PprodA_PprodB.py | bind3_prodA_[00-30]_prodB_[00-30].pickle | bind_sweep_PprodA_PprodB_aggregate.py | bind3.pickle |
| Fig S7F-H | revbind_sweep_PprodA_PprodB.py | revbind4_prodA_[00-30]_prodB_[00-30].pickle | revbind_sweep_PprodA_PprodB_aggregate.py | revbind4.pickle |
| Fig S8B,E | satprod_time_sweep_PprodA.py | satprod_time_PprodAsweep_[0-3].pickle | | |
| Fig S8C, F | satprod_time_sweep_kcatA.py | satprod_time_kcatAsweep_[0-3].pickle | | |
| Fig S8D, D', G, G' | satprod_time_sweep_Tcc.py | satprod_time_Tccsweep_[0-3].pickle | | |
| Fig S9 | proddeg_rkcat_sweep.py | Rkcatsweep2_[0-19].pickle | | |
| Fig S10 | phosint.py | phos_int3_prodA_[00-30]_prodB_[00-30].pickle | phosint_aggregate.py | phos_int3.pickle |
| Fig S11 | phos2_sweep_PprodA_PprodB.py | phos2_0_prodA_[00-30]_prodB_[00-30].pickle | phos2_sweep_PprodA_PprodB_aggregate.py | phos2_0.pickle |
| Fig S12 | cascade6_time.py | cascade5_time.pickle | | |
| Fig S14 | cascade_spatial.py | cascade_10gen2.pickle | cascade_spatial_analyze.py | cascade_10gen2_cousinmaps_all.pickle |
| Fig S15 | cascade_spatial.py | cascade_10gen2.pickle | cascade_spatial_analyze.py | grid_[1000-1002]_relatedness.pickle |
| Fig S16 | cascade_spatial_moranI.py | "cascade_10gen2_morIs_discdist_r[0-9].pickle | cascade_10gen2_morIs_donut_r[0-9].pickle | cascade_10gen2_morIs_gausdist_r[0-9].pickle |
| Fig S18 | prodsat_time.py | prodsat_time.pickle | calc_correlation.py | correlationcoef.pickle |
| Fig S20B | phoscycle_sweep_PprodA_PprodB.py | phoscycle2_PprodA_[00-30]_PprodB_[00-30].pickle | phoscycle_sweep_PprodA_PprodB_aggregate.py | phoscycle2.pickle |
| Fig S20C | phossat_sweep_PprodB_Km.py | phossat3_prodB_[00-30]_km_[00-24].pickle | phossat_sweep PprodB_Km_aggregate.py | satphos3.pickle |
| Fig S21 | produnsat_order_sweep_PprodB.py | order4_PprodB_[00-27].pickle | produnsat_order_sweep_PprodB_aggregate.py | calcOrder4.pickle |
